# Patient-specific Boolean models of signaling networks guide personalized treatments

**DOI:** 10.1101/2021.07.28.454126

**Authors:** Arnau Montagud, Jonas Béal, Luis Tobalina, Pauline Traynard, Vigneshwari Subramanian, Bence Szalai, Róbert Alföldi, László Puskás, Alfonso Valencia, Emmanuel Barillot, Julio Saez-Rodriguez, Laurence Calzone

**Affiliations:** Institut Curie, PSL Research University, Paris, France; INSERM, U900, Paris, France; MINES ParisTech, PSL Research University, CBIO-Centre for Computational Biology, Paris, France; Barcelona Supercomputing Center (BSC), Barcelona, Spain; Faculty of Medicine, Joint Research Centre for Computational Biomedicine (JRC−COMBINE), RWTH Aachen University, 52074 Aachen, Germany; Semmelweis University, Faculty of Medicine, Department of Physiology, Budapest, Hungary; Astridbio Technologies Ltd., 6728 Szeged, Hungary; Institució Catalana de Recerca i Estudis Avançats (ICREA), 08010, Barcelona, Spain; Faculty of Medicine and Heidelberg University Hospital, Institute of Computational Biomedicine, Heidelberg University, Heidelberg, Germany

## Abstract

Prostate cancer is the second most occurring cancer in men worldwide. To better understand the mechanisms of tumorigenesis and possible treatment responses, we developed a mathematical model of prostate cancer which considers the major signalling pathways known to be deregulated.

We personalised this Boolean model to molecular data to reflect the heterogeneity and specific response to perturbations of cancer patients. 488 prostate samples were used to build patient-specific models and compared to available clinical data. Additionally, eight prostate cell-line-specific models were built to validate our approach with dose-response data of several drugs.

The effects of single and combined drugs were tested in these models under different growth conditions. We identified 15 actionable points of interventions in one cell-line-specific model whose inactivation hinders tumorigenesis. To validate these results, we tested nine small molecule inhibitors of five of those putative targets and found a dose-dependent effect on four of them, notably those targeting HSP90 and PI3K. These results highlight the predictive power of our personalized Boolean models and illustrate how they can be used for precision oncology.

## Introduction

Like most cancers, prostate cancer arises from mutations in single somatic cells that induce deregulations in processes such as proliferation, invasion of adjacent tissues and metastasis. Not all prostate patients respond to the treatments in the same way, depending on the stage and type of their tumour (Chen & Zhou, 2016) as well as to differences in their genetic and epigenetic profiles (Yang *et al*, 2018; Toth *et al*, 2019). The high heterogeneity of these profiles can be explained by a large number of interacting proteins and the complex cross-talks between the cell signalling pathways that can be altered in cancer cells. Because of this complexity, understanding the process of tumorigenesis and tumour growth would benefit from a systemic and dynamical description of the disease. At the molecular level, this can be tackled by a simplified mechanistic cell-wide model of protein interactions of the underlying pathways, dependent on external environmental signals.

Although continuous mathematical modelling has been widely used to study cellular biochemistry dynamics (e.g., ordinary differential equations) (Goldbeter, 2002; Tyson *et al*, 2019; Le Novère, 2015; Sible & Tyson, 2007; Kholodenko *et al*, 1995), this formalism does not scale up well to large signalling networks, due to the difficulty of estimating kinetic parameter values (Babtie & Stumpf, 2017). In contrast, the logical (or logic) modelling formalism represents a simpler mean of abstraction where the causal relationships between proteins (or genes) are encoded with logic statements and dynamical behaviours are represented by transitions between discrete states of the system (Thomas, 1973; Kauffman, 1969). In particular, Boolean models, the simplest implementation of logical models, describes each protein as a binary variable (ON/OFF). This framework is flexible, requires in principle no quantitative information, can be hence applied to large networks combining multiple pathways, and can also provide a qualitative understanding of molecular systems lacking mechanistic detailed information.

In the last years, logical and in particular Boolean modelling has successfully been used to describe the dynamics of human cellular signal transduction and gene regulations (Helikar *et al*, 2008; Calzone *et al*, 2010; Grieco *et al*, 2013; Flobak *et al*, 2015; Cho *et al*, 2016; Traynard *et al*, 2016) and their deregulation in cancer (Fumiã & Martins, 2013; Hu *et al*, 2015). Numerous applications of logical modelling have shown that this framework is able to delineate the main dynamical properties of complex biological regulatory networks (Faure *et al*, 2006; Abou-Jaoudé *et al*, 2011).

However, the Boolean approach is purely qualitative and does not consider real time of cellular events (half time of proteins, triggering of apoptosis, etc.). To cope with this issue, we developed the MaBoSS software to compute continuous Markov Chain simulations on the model state transition graph (STG), in which a model state is defined as a vector of nodes that are either active or inactive. In practice, MaBoSS associates transition rates for activation and inhibition of each node of the network, enabling it to account for different time scales of the processes described by the model. Given some initial conditions, MaBoSS applies a Monte-Carlo kinetic algorithm (or Gillespie algorithm) to the STG to produce time trajectories (Stoll *et al*, 2012, 2017) such that time evolution of the model state probabilities can be estimated.

Stochastic simulations can easily explore the model dynamics with different initial conditions by varying the probability of having a node active at the beginning of the simulations and by modifying the model such that it accounts for genetic and environmental perturbations (e.g., presence or absence of growth factors, or death receptors). For each case, the effect on the probabilities of selected read-outs can be measured (Cohen *et al*, 2015; Montagud *et al*, 2017).

When summarizing the biological knowledge into a network and translating it into logical terms, the obtained model is generic and cannot explain the differences and heterogeneity between patients’ responses to treatments. Models can be trained with dedicated perturbation experiments (Saez-Rodriguez *et al*, 2009; Dorier *et al*, 2016), but such data can only be obtained with non-standard procedures such as microfluidics from patients’ material (Eduati *et al*, 2020). To address this limitation, we developed a methodology to use different omics data that are more commonly available to personalise generic models to individual cancer patients or cell lines and verified that the obtained models correlated with clinical results such as patient survival information (Béal *et al*, 2019). In present work, we apply this approach to prostate cancer to suggest targeted therapy to patients based on their omics profile (Figure 1). We first built 488 patient- and eight cell line-prostate-specific models using data from The Cancer Genome Atlas (TCGA) and the Genomics of Drug Sensitivity in Cancer (GDSC) projects, respectively. Simulating these models with the MaBoSS framework, we identified points of intervention that diminish the probability of reaching pro-tumorigenic phenotypes. Lastly, we developed a new methodology to simulate drug effects on these data-tailored Boolean models and present a list of viable drugs and regimes that could be used on these patient-and cell-line-specific models for optimal results. Experimental validations were performed on the LNCaP prostate cell line with two predicted targets, confirming the predictions of the model.

**Figure 1:**
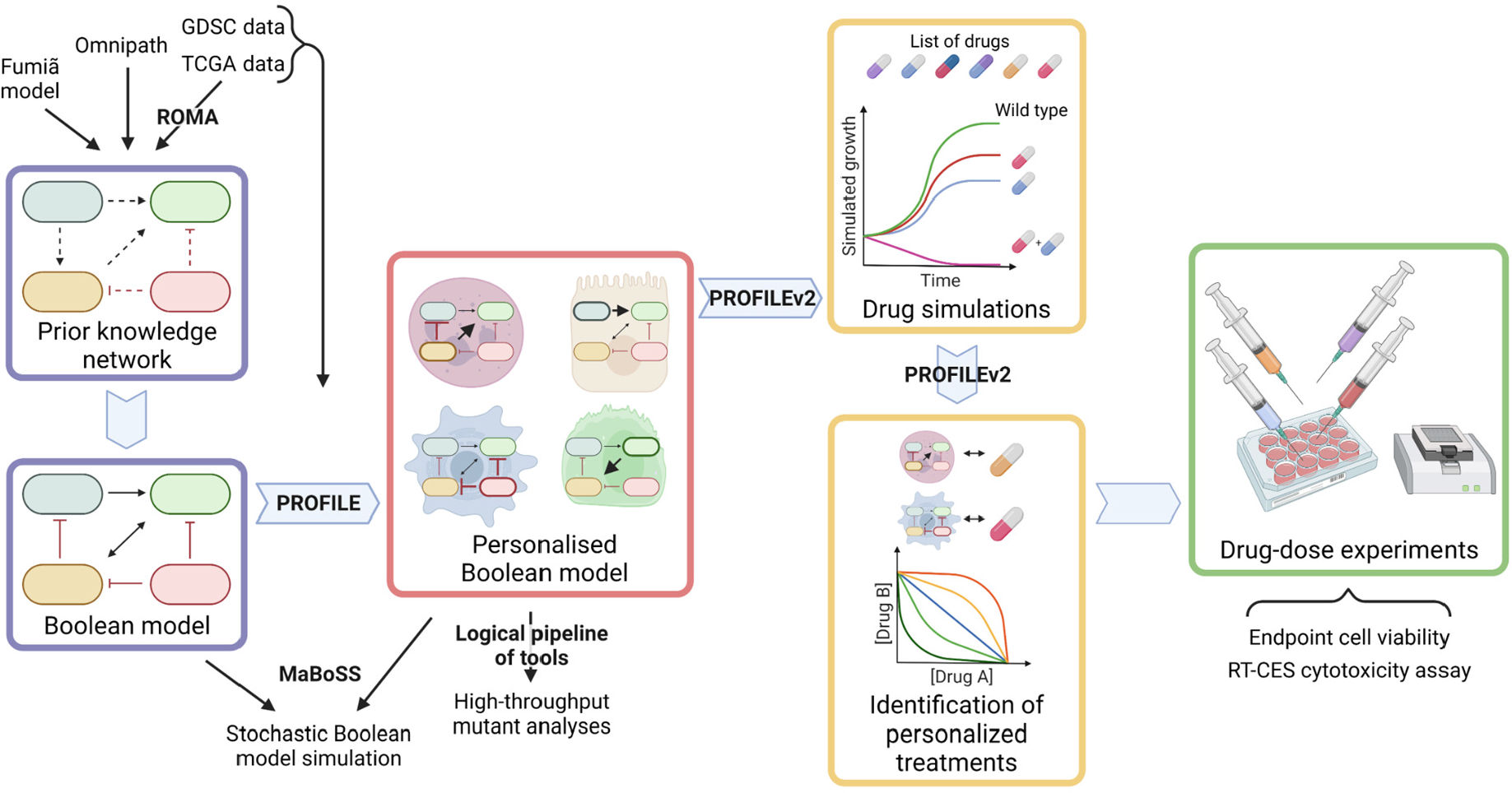
Workflow to build patient-specific Boolean models and to uncover personalized drug treatments from present work. We gathered data from Fumiã and Martins (2013) Boolean model, Omnipath (Türei *et al*, 2021) and pathways identified with ROMA (Martignetti *et al*, 2016) on the TCGA data to build a prostate-specific prior knowledge network. This network was manually converted into a prostate Boolean model that could be stochastically simulated using MaBoSS (Stoll *et al*, 2017) and tailored to different TCGA and GDSC datasets using our PROFILE tool to have personalized Boolean models. Then, we studied all the possible single and double mutants on these tailored models using our logical pipeline of tools (Montagud *et al*, 2017). Using these personalized models and our PROFILE_v2 tool presented in this work, we obtained tailored drug simulations and drug treatments for 488 TCGA patients and eight prostate cell lines. Lastly, we performed drug-dose experiments on a short list of candidate drugs that were particularly interesting in the LNCaP prostate cell line. Created with BioRender.com.

## Results

### Prostate Boolean model construction

A network of signalling pathways and genes relevant for prostate cancer progression was assembled to recapitulate the potential deregulations that lead to high-grade tumours. Dynamical properties were added onto this network to perform simulations, uncover therapeutic targets and explore drug combinations. The model was built upon a generic cancer Boolean model by Fumiã and Martins (2013), which integrates major signalling pathways, and their substantial cross-talks. The pathways include the regulation of cell death and proliferation in many tumours.

This initial generic network was extended to include prostate-cancer-specific genes (e.g., SPOP, AR, etc.), pathways identified using ROMA (Martignetti *et al*, 2016), OmniPath (Türei *et al*, 2021) and up-to-date literature. ROMA is applied on omics data, either transcriptomics or proteomics. In each pathway, the genes that contribute the most to the overdispersion are selected. ROMA was applied to the TCGA transcriptomics data using gene sets from cancer pathway databases (Appendix File, Figure S1). These results were used as guidelines to extend the network to fully cover the alterations found in prostate cancer patients. OmniPath was used to complete our network finding connections between the proteins of interest known to play a role in prostate and the ones identified with ROMA, and the list of genes already present in the model. The final network includes pathways such as androgen receptor, MAPK, Wnt, NFkB, PI3K/AKT, MAPK, mTOR, SHH, the cell cycle, the epithelial-mesenchymal transition (EMT), apoptosis and DNA damage pathways.

This network was then converted into a Boolean model where all variables can take two values: 0 (inactivate or absent) or 1 (activate or present). Our model aims at predicting prostate phenotypic behaviours for healthy and cancer cells in different conditions. Nine inputs that represent some of these physiological conditions of interest were considered: *EGF, FGF, TGF beta, Nutrients, Hypoxia, Acidosis, Androgen, TNF alpha* and *Carcinogen*. These input nodes have no regulation and their values are fixed for each simulation, representing the cell’s microenvironmental characteristics.

We defined six variables as output nodes that allow the integration of multiple phenotypic signals and simplify the analysis of the model. Two of these phenotypes represent the possible growth status of the cell: *Proliferation* and *Apoptosis*. *Apoptosis* is activated by Caspase 8 or Caspase 9, while *Proliferation* is activated by cyclins D and B (read-outs of the G1 and M phases, respectively). The *Proliferation* output is described in published models as specific stationary protein activation patterns, namely the following sequence of activation of cyclins:

Cyclin D, then Cyclin E, then Cyclin A, and finally Cyclin B (Traynard *et al*, 2016). Here, we considered a proper sequence when Cyclin D activates first, allowing the release of the transcriptional factor E2F1 from the inhibitory complex it was forming with RB (retinoblastoma protein), and then triggering a series of events leading to the activation of Cyclin B, responsible for the cell’s entry into mitosis (Appendix File, Figure S4). We also define several phenotypic outputs that are readouts of cancer hallmarks: *Invasion, Migration,* (bone) *Metastasis* and *DNA repair*. The final model accounts for 133 nodes and 449 edges (Figure 2, SuppFile 1, and in GINsim format at the address: http://ginsim.org/model/signalling-prostate-cancer).

**Figure 2:**
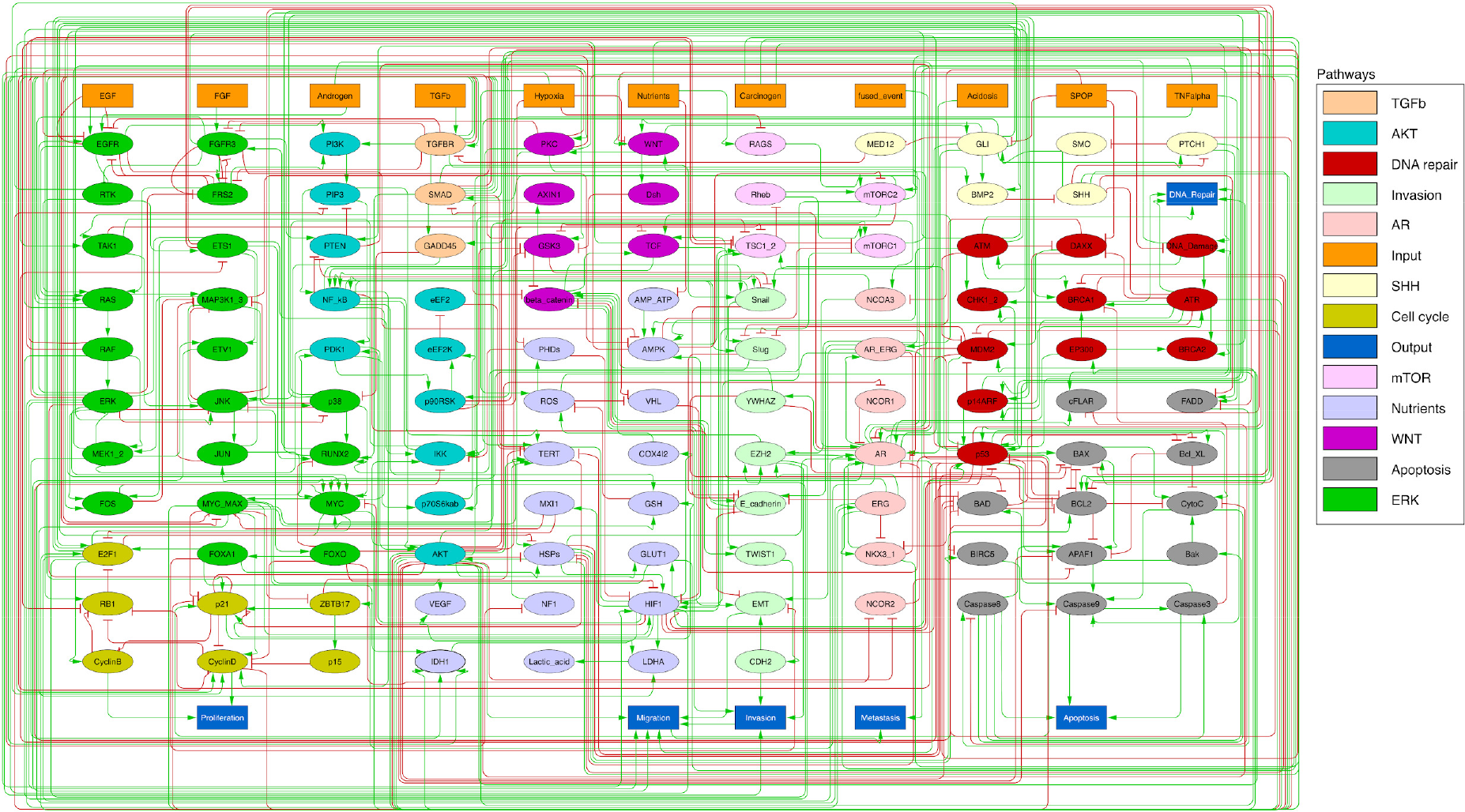
Prostate Boolean model used in present work. Nodes (ellipses) represent biological entities, and arcs are positive (green) or negative (red) influences of one entity on another one. Orange rectangles correspond to inputs (from left to right: EGF, FGF, TGFb, Nutrients, Hypoxia, Acidosis, Androgen, fused_event, TNFalpha, SPOP, Carcinogen) and dark blue rectangles to phenotypes (from left to right: Proliferation, Migration, Invasion, Metastasis, Apoptosis, DNA_repair), the read-outs of the model.

### Prostate Boolean model simulation

The model can be considered as a model of healthy prostate cells when no mutants (or fused genes) are present. We refer to this model as the wild type model. These healthy cells mostly exhibit quiescence (neither proliferation nor apoptosis) in the absence of any input (Figure 3A). When *Nutrients* and growth factors (*EGF* or *FGF*) are present, *Proliferation* is activated (Figure 3B). *Androgen* is necessary for AR activation and helps in the activation of *Proliferation*, even though it is not necessary when *Nutrients* or growth factors are present. Cell death factors (such as Caspase 8 or 9) trigger *Apoptosis* in the absence of *SPOP*, while *Hypoxia* and *Carcinogen* facilitate apoptosis, but are not necessary if cell death factors are present (Figure 3C).

**Figure 3:**
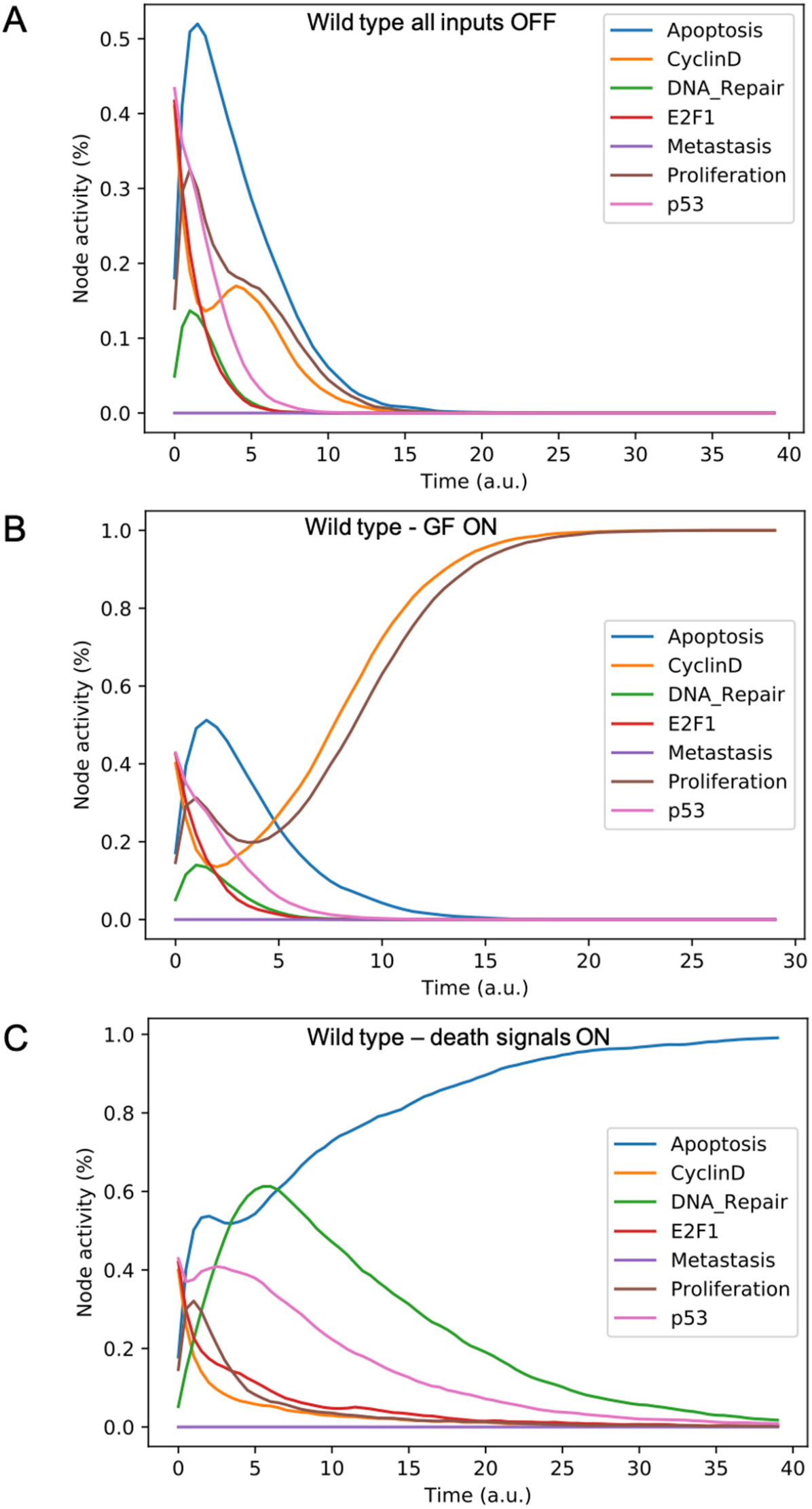
Prostate Boolean model MaBoSS simulations. (A) The model was simulated with all initial inputs set to 0 and all other variables random. All phenotypes are 0 at the end of the simulations, which should be understood as a quiescent state, where neither proliferation nor apoptosis are active. (B) The model was simulated with growth factors (*EGF* and *FGF*), *Nutrients* and *Androgen* ON. (C) The model was simulated with *Carcinogen*, *Androgen*, *TNFalpha*, *Acidosis*, and *Hypoxia* ON.

In our model, the progression towards metastasis is described as a stepwise process. *Invasion* is first activated by known pro-invasive proteins: either β-catenin (Francis *et al*, 2013) or a combination of *CDH2* (De Wever *et al*, 2004), *SMAD* (Daroqui *et al*, 2012) or *EZH2* (Ren *et al*, 2012). *Migration* is then activated by *Invasion* and *EMT* and with either *AKT* or *AR* (Castoria *et al*, 2011). Lastly, (bone) *Metastasis* is activated by *Migration* and one of three nodes: *RUNX2* (Altieri *et al*, 2009), *ERG* (Adamo & Ladomery, 2016) or ERG fused with TMPRSS2 (St John *et al*, 2012), FLI1, ETV1 or ETV4 (The Cancer Genome Atlas Research Network, 2015).

This prostate Boolean model was simulated stochastically using MaBoSS (Stoll *et al*, 2012, 2017) and validated by recapitulating known phenotypes of prostate cells under physiological conditions (Figure 3). In particular, we tested that combinations of inputs lead to non-aberrant phenotypes such as growth factors leading to apoptosis in wild type conditions; we also verified that the cell cycle events occur in a proper order: as CyclinD gets activated, RB1 is phosphorylated and turned OFF, allowing E2F1 to mediate the synthesis of CyclinB (see SuppFile 2 for the jupyter notebook and the simulation of diverse cellular conditions).

### Personalisation of the prostate Boolean model

#### Personalised TCGA prostate cancer patient Boolean models

We tailored the generic prostate Boolean model to a set of 488 TCGA prostate cancer patients (Appendix File, Figure S7) using our personalisation method (PROFILE, (Béal *et al*, 2019)), constructing 488 individual Boolean models, one for each patient. Personalised models were built using three types of data: discrete data such as mutations and copy number alterations (CNA) and continuous data such as RNAseq data. For discrete data, the nodes corresponding to the mutations or the CNA were forced to 0 or 1 according to the effect of alterations, based on *a priori* knowledge (i.e., if the mutation was reported to be activating or inhibiting the gene’s activity). For continuous data, the personalisation method modifies the value for the transition rates of model variables and their initial conditions to influence the probability of some transitions. This corresponds, in a biologically-meaningful way, to translating genetic mutations as lasting modifications making the gene independent of regulation, and to translating RNA expression levels as modulation of a signal but not changing the regulation rules (see Materials and Methods and in Appendix File, Figure S8-S12).

We assess the general behaviour of the individual patient-specific models by comparing the model outputs (i.e., probabilities to reach certain phenotypes) with clinical data. Here, the clinical data consist of a Gleason score associated with each patient, which in turn corresponds to the gravity of the tumour based on its appearance and the stage of invasion (Gleason, 1977, 1992; Chen & Zhou, 2016). We gathered output probabilities for all patient-specific models and confronted them to their Gleason scores. The phenotype *DNA_repair*, which can be interpreted as a sensor of DNA damage and genome integrity which could lead to DNA repair, seems to separate low and high Gleason scores (Figure 4A) confirming that DNA damage pathways are activated in patients (Marshall *et al*, 2019) but may not lead to the triggering of apoptosis in this model (Figure S9). Also, the centroids of Gleason groups tend to move following *Proliferation*, *Migration* and *Invasion* variables. We then looked at the profiles of the phenotype scores across patients and their Gleason group and found that the density of high *Proliferation* score (close to 1, Figure 4B) tends to increase as the Gleason score increases (from low to intermediate to high). The *Apoptosis* phenotype, however, only shows a slight change in probabilities in groups with low or high Gleason scores (Figure 4C).

**Figure 4:**
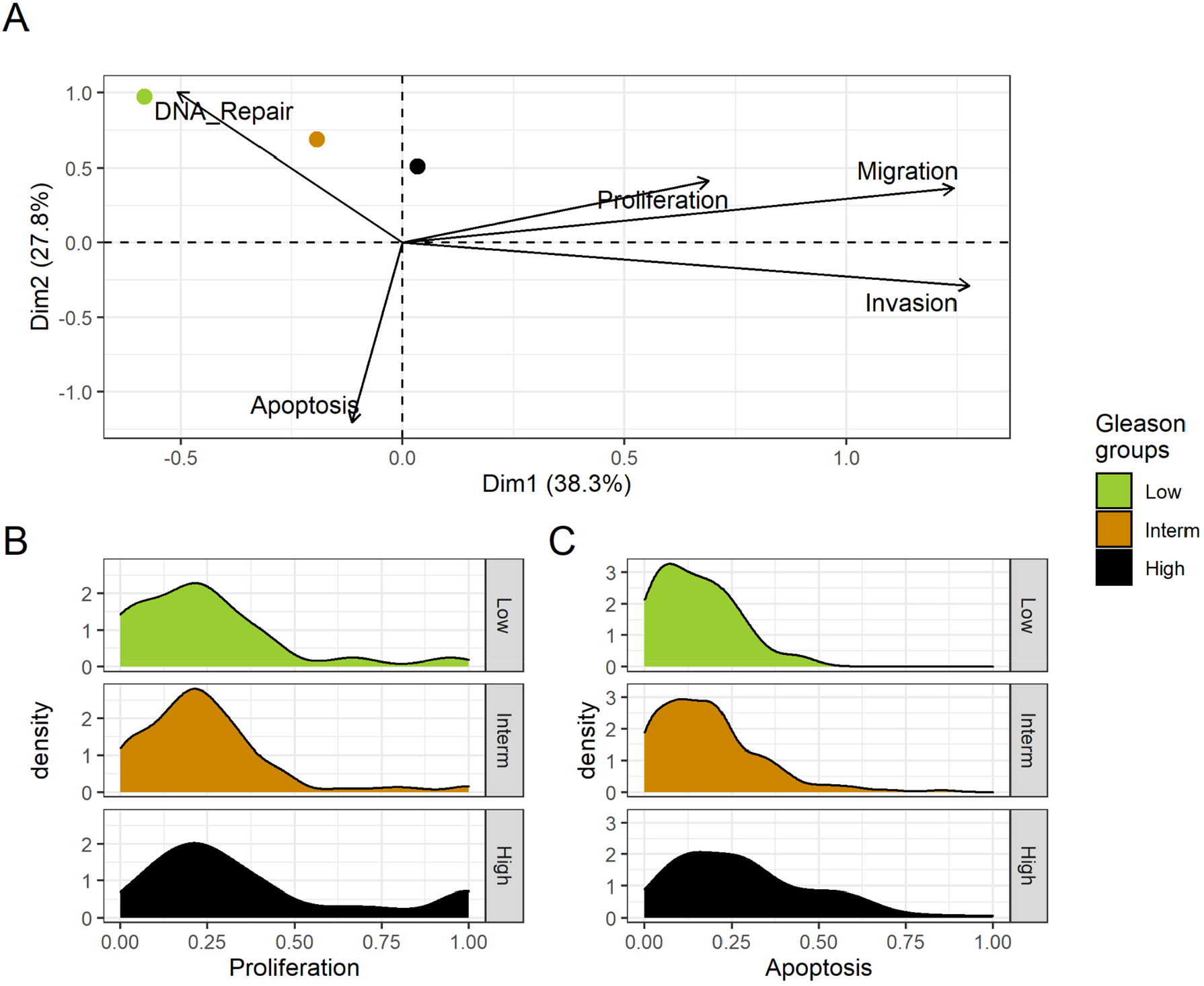
Associations between simulations and Gleason groups (GG). A) Centroids of the PCA of the samples according to their GG. The personalisation recipe used was mutations and copy number alterations (CNA) as discrete data and RNAseq as continuous data. Density plots of *Proliferation* (B) and *Apoptosis* (C) scores according to GG; each vignette corresponds to a specific sub-cohort with a fixed GG.

#### Personalised drug predictions of TCGA Boolean models

Using the 488 TCGA-patient-specific models, we looked in each patient for genes that, when inhibited, hamper *Proliferation* or promote *Apoptosis* in the model. We focused on these inhibitions as most drugs interfere with the protein activity related to these genes, even though our methodology allows us to study increased protein activity related to over-expression of genes as well (Montagud *et al*, 2017; Béal *et al*, 2019). Interestingly, we found several genes that were found as suitable points of intervention in most of the patients (MYC_MAX complex and SPOP were identified in more than 80% of the cases) (Appendix File, Figure S17 and S18), but others were specific to only some of the patients (MXI1 was identified in only 4 patients, 1% of the total, GLI in only 7% and WNT in 8% of patients). All the TCGA-specific personalised models can be found in SuppFile 3 and the TCGA mutants and their phenotype scores can be found in SuppFile 4. Furthermore, we explored the possibility of finding combinations of treatments that could reduce the *Proliferation* phenotype. To lower the computational power need, we have narrowed down the list of potential candidates to reduce *Proliferation* or increase *Apoptosis* by performing the analysis of all the single perturbations and selecting the combined perturbations of a set of selected genes that are targets of already-developed drugs relevant in cancer progression (Table 1).

**Table 1:** List of selected nodes, their corresponding genes and drugs that were included in the drug analysis of the models tailored for TCGA patients and LNCaP cell line.

| Node | Gene | Compound / Inhibitor name | Clinical stage | Source |
| --- | --- | --- | --- | --- |
| AKT | AKT1, AKT2, AKT3 | PI-103 | Preclinical | Drug Bank |
|  |  | Enzastaurin | Phase 3 | Drug Bank |
|  |  | Archexin, Pictilisib | Phase 2 | Drug Bank |
| AR | AR | Abiraterone, Enzalutamide, Formestane, Testosterone propionate | Approved | Drug Bank |
|  |  | 5alpha-androstan-3beta-ol | Preclinical | Drug Bank |
| Caspase8 | CASP8 | Bardoxolone | Preclinical | Drug Bank |
| cFLAR | CFLAR | - | - | - |
| EGFR | EGFR | Afatinib, Osimertinib, Neratinib, Erlotinib, Gefitinib | Approved | Drug Bank |
|  |  | Varlitinib | Phase 3 | Drug Bank |
|  |  | Olmutinib, Pelitinib | Phase 2 | Drug Bank |
| ERK | MAPK1 | Isoprenaline | Approved | Drug Bank |
|  |  | Perifosine | Phase 3 | Drug Bank |
|  |  | Turpentine, SB220025, | Preclinical | Drug Bank |
|  |  | Olomoucine,<br>Phosphonothreoni<br>ne |  |  |
|  | MAPK3,<br>MAPK1 | Arsenic trioxide | Approved | Drug Bank |
|  |  | Ulixertinib,<br>Seliciclib | Phase 2 | Drug Bank |
|  |  | Purvalanol | Preclinical | Drug Bank |
|  | MAPK3 | Sulindac,<br>Cholecystokinin | Approved | Drug Bank |
|  |  | 5-iodotubercidin | Preclinical | Drug Bank |
| GLUT1 | SLC2A1 | Resveratrol | Phase 4 | Drug Bank |
| HIF-1 | HIF1A | CAY-10585 | Preclinical | Drug Bank |
| HSPs | HSP90AA1,<br>HSP90AB1,<br>HSP90B1,<br>HSPA1A,<br>HSPA1B,<br>HSPB1 | Cladribine | Approved | Drug Bank |
|  |  | 17-DMAG | Phase 2 | Drug Bank |
|  |  | NMS-E973 | Preclinical | Drug Bank |
| MEK1_2 | MAP2K1,<br>MAP2K2 | Trametinib,<br>Selumetinib | Approved | Drug Bank |
|  |  | Perifosine | Phase 3 | Drug Bank |
|  |  | PD184352 (CI-1040) | Phase 2 | Drug Bank |
| MYC_MAX | complex of<br>MYC and<br>MAX | 10058-F4 (for<br>MAX) | Preclinical | Drug Bank |
| p14ARF | CDKN2A | - | - | - |
| PI3K |  | PI-103 | Preclinical | Drug Bank |
|  | PIK3CA,<br>PIK3CB,<br>PIK3CG,<br>PIK3CD,<br>PIK3R1,<br>PIK3R2,<br>PIK3R3,<br>PIK3R4,<br>PIK3R5,<br>PIK3R6,<br>PIK3C2A,<br>PIK3C2B,<br>PIK3C2G,<br>PIK3C3 | Pictilisib | Phase 2 | Drug Bank |
| ROS | NOX1,<br>NOX3,<br>NOX4 | Fostamatinib | Approved | Drug Bank |
|  | NOX2 | Dextromethorphan | Approved | Drug Bank |
|  |  | Tetrahydroisoquino<br>lines<br>(CHEMBL3733336<br>,<br>CHEMBL3347550,<br>CHEMBL3347551) | Preclinical | ChEMBL |
| SPOP | SPOP | -- | -- | -- |
| TERT | TERT | Grn163l | Phase 2 | Drug Bank |
|  |  | BIBR 1532 | Preclinical | ChEMBL |

We used the models to grade the effect that the combined treatments have in each one of the 488 TCGA-patient-specific models’ phenotypes. This list of combinations of treatments can be used to compare the effects of drugs on each TCGA patient and allows us to propose some of them for individual patients and to suggest drugs suitable to groups of patients (SuppFile 4). Indeed, the inactivation of some of the targeted genes had a greater effect in some patients than in others, suggesting the possibility for the design of personalised drug treatments. For instance, for the TCGA-EJ-5527 patient, the use of MYC_MAX complex inhibitor reduced *Proliferation* to 66%. For this patient, combining MYC_MAX with other inhibitors, such as AR or AKT did not further reduce *Proliferation* score (67% in these cases). Other patients have MYC_MAX as an interesting drug target, but the inhibition of this complex did not have such a dramatic effect in their *Proliferation* scores as in the case of TCGA-EJ-5527. Likewise, for the TCGA-H9-A6BX patient, the use of SPOP inhibitor increased *Apoptosis* by 87%, while the use of a combination of cFLAR and SPOP inhibitors further increased *Apoptosis* by 89%. For the rest of this section, we focus on the analysis of clinical groups rather than individuals.

Studying the decrease of *Proliferation*, we found that AKT is the top hit in Gleason Groups 1, 2, and 3, seconded by SPOP in Group 1, PIP3 in Group 2 and MYC_MAX in Group 3. MYC_MAX is the top hit in Group 4, seconded by AR. In regards to the increase of *Apoptosis*, SPOP is the top hit in all groups. SSH is second in Groups 1 and 2 and AKT in Group 3. It is interesting to note here that many of these genes are targeted by drugs (Table 1). Notably, AR is the target of the drug Enzalutamide, which is indicated for men with an advanced stage of the disease (Scott, 2018), or that MYC is the target of BET bromodomain inhibitors and are generally effective in castration-resistant prostate cancer cases (Coleman *et al*, 2019).

The work on patient data provided some possible insights and suggested patient- and group-specific potential targets. To validate experimentally our approach, we personalised the prostate model to different prostate cell lines where we performed drug assays to confirm the predictions of the model.

#### Personalised drug predictions of LNCaP Boolean model

We applied the methodology for personalisation of the prostate model to eight prostate cell lines available in GDSC (Iorio *et al*, 2016): 22RV1, BPH-1, DU-145, NCI-H660, PC-3, PWR-1E and VCaP (results in Appendix file and are publicly available in SuppFile 5). We decided to focus the validation on one cell line, LNCaP.

LNCaP, first isolated from a human metastatic prostate adenocarcinoma found in a lymph node (Horoszewicz *et al*, 1983), is one of the most widely used cell lines for prostate cancer studies. Androgen-sensitive LNCaP cells are representative of patients sensitive to treatments as opposed to resistant cell lines such as DU-145. Additionally, LNCaP cells have been used to obtain numerous subsequent derivatives with different characteristics (Cunningham & You, 2015).

The LNCaP personalisation was performed based on mutations as discrete data and RNA-Seq as continuous data. The resulting LNCaP-specific Boolean model was then used to identify all possible combinations of mutations (interpreted as effects of therapies) and to study the synergy of these perturbations. For that purpose, we automatically performed single and double mutant analyses on the LNCaP-specific model (knock-out and overexpression) (Montagud *et al*, 2017) and focused on the model phenotype probabilities as read-outs of the simulations. The analysis of the complete set of simulations for the 32258 mutants can be found in the Appendix File and in SuppFile 6, where the LNCaP-cell-line-specific mutants and their phenotype scores are reported for all mutants. Among all combinations, we identified the top 20 knock-out mutations that depleted *Proliferation* or increased *Apoptosis* the most. As some of them overlapped, we ended up with 29 nodes: *AKT, AR, ATR, AXIN1, Bak, BIRC5, CDH2, cFLAR, CyclinB, CyclinD, E2F1, eEF2K, eEF2, eEF2K, EGFR, ERK, HSPs, MED12, mTORC1, mTORC2, MYC, MYC_MAX, PHDs, PI3K, PIP3, SPOP, TAK1, TWIST1, and VHL*.

We used the scores of these nodes to further trim down the list to have 10 final nodes (*AKT, AR, cFLAR, EGFR, ERK, HSPs, MYC_MAX, SPOP* and *PI3K*) and added 7 other nodes whose genes are considered relevant in cancer biology, such as *AR_ERG* fusion, *Caspase8*, *HIF1*, *GLUT1, MEK1_2*, *p14ARF*, *ROS* and *TERT* (Table 1). We did not consider the overexpression mutants as they have a very difficult translation to drug uses and clinical practices.

To further analyse the mutant effects, we simulated the LNCaP model with increasing node inhibition values to mimic the effect of drugs’ dosages using a methodology we specifically developed for these purposes (PROFILE_v2). Six simulations were done for each inhibited node, with 100% of node activity (no inhibition), 80%, 60%, 40%, 20% and 0% (full knock-out) (see Methods). A nutrient-rich media with EGF was used for these simulations and we show results on three additional sets of initial conditions in the Appendix File, Figure S24: a nutrient-rich media with androgen, with androgen and EGF, and with none, that correspond to experimental conditions that are tested here. We applied this gradual inhibition, using increasing drugs’ concentrations, to a reduced list of drug-targeted genes relevant for cancer progression (Table 1). We confirmed that the inhibition of different nodes affected differently the probabilities of the outputs (Appendix File, Figure S29 and S30). Notably, *Apoptosis* score was slightly promoted when knocking out *SPOP* under all growth conditions (Appendix File, Figure S30). Likewise, *Proliferation* depletion was accomplished when *HSPs* or *MYC_MAX* were inhibited under all conditions and, less notably, when *ERK, EGFR*, *SPOP* or *PI3K* were inhibited (Appendix File, Figure S30).

Additionally, these gradual inhibition analyses can be combined to study the interaction of two simultaneously inhibiting nodes (Appendix File, Figure S31 and S32). For instance, the combined gradual inhibition of *ERK* and *MYC_MAX* nodes affects *Proliferation* score in a balanced manner (Figure 5A) even though *MYC_MAX* seems to affect this phenotype more, notably at low activity levels. By extracting subnetworks of interaction around *ERK* and *MYC_MAX* and comparing them, we found that the pathways they belong to have complementary downstream targets participating in cell proliferation through targets in MAPK and cell cycle pathways. This complementarity could explain the synergistic effects observed (Figure 5A and 5C).

**Figure 5:**
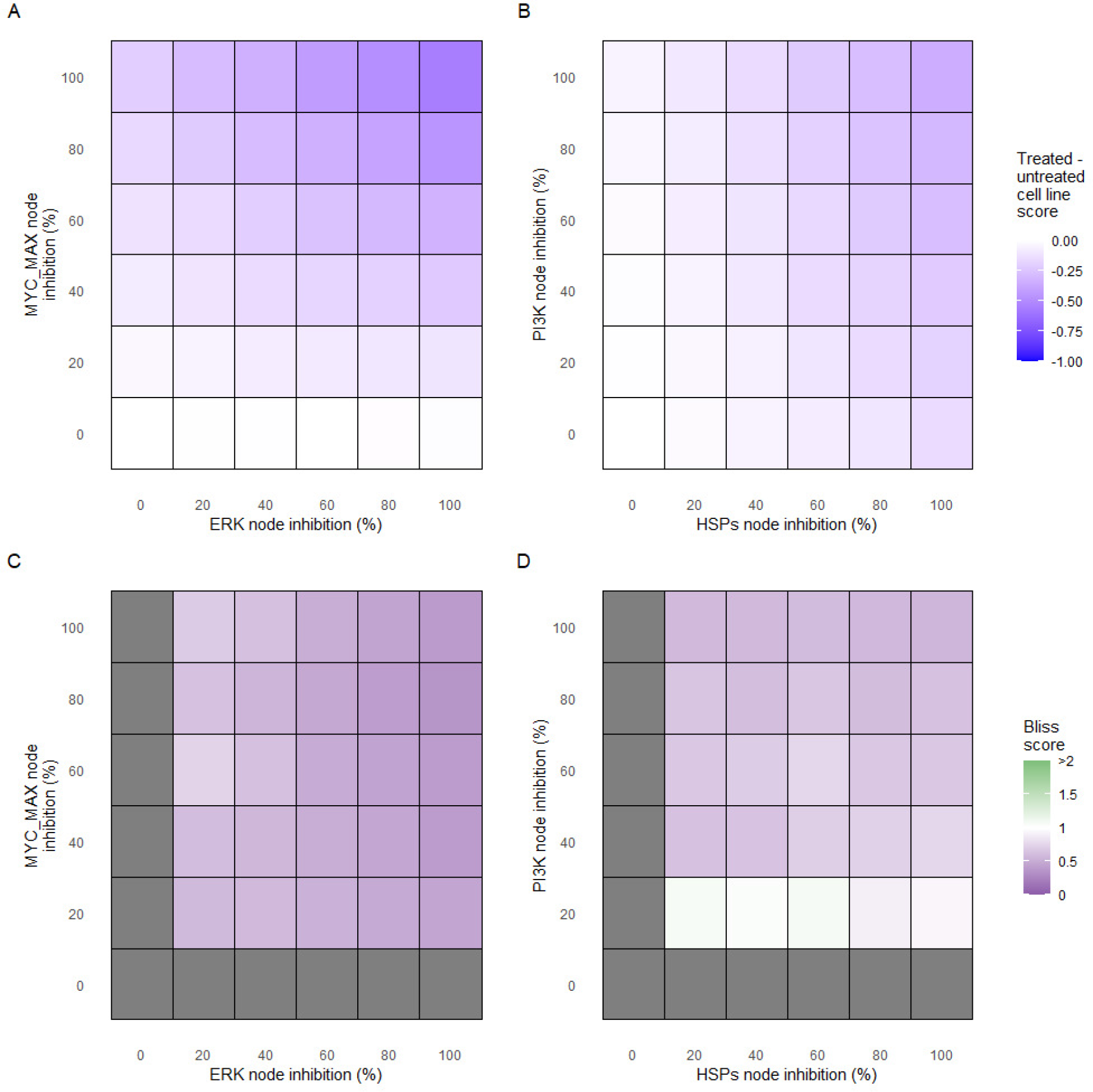
Phenotype score variations and synergy upon combined ERK and MYC_MAX (A and C) and HSPs and PI3K (B and D) inhibition under *EGF* growth condition. Proliferation score variation (A) and Bliss Independence synergy score (C) with increased node activation of nodes ERK and MYC_MAX. Proliferation score variation (B) and Bliss Independence synergy score (D) with increased node activation of nodes HSPs and PI3K. Bliss Independence synergy score < 1 is characteristic of drug synergy, grey colour means one of the drugs is absent and thus no synergy score is available.

Lastly, drug synergies can be studied using Bliss Independence using the results from single and combined simulations with gradual inhibitions. This score compares the combined effect of two drugs with the effect of each one of them, with a synergy when the value of this score is lower than 1. We found that the combined inhibition of *ERK* and *MYC_MAX* nodes on *Proliferation* score was synergistic (Figure 5C). Another synergistic pair is the combined gradual inhibition of *HSPs* and *PI3K* nodes that also affects *Proliferation* score in a joint manner (Figure 5B), with some Bliss Independence synergy found (Figure 5D). A complete study on the Bliss Independence synergy of all the drugs considered in present work on *Proliferation* and *Apoptosis* phenotypes can be found in Appendix File, Figure S33.

### Experimental validation of predicted targets

#### Drugs associated with the proposed targets

To identify drugs that could act as potential inhibitors of the genes identified with the Boolean model, we explored the drug-target associations in DrugBank (Wishart et al, 2018) and ChEMBL (Gaulton *et al*, 2017). We found drugs that targeted almost all genes corresponding to the nodes of interest in Table 1, except for cFLAR, p14ARF and SPOP. However, we could not identify experimental cases where drugs targeting both members of the proposed combinations were available (Appendix File and in SuppFile 6). One possible explanation is that the combinations predicted by the model suggest in some cases to overexpress the potential target and most of the drugs available act as inhibitors of their targets.

Using the cell-line specific models, we tested if the LNCaP cell line was more sensitive than the rest of the prostate cell lines to the LNCaP-specific drugs identified in Table 1. We compared GDSC’s Z-score of these drugs in LNCaP with their Z-scores in all GDSC cell lines (Figure 6). We observed that LNCaP is more sensitive to drugs targeting AKT or TERT than the rest of the studied prostate cell lines. Furthermore, we saw that the drugs that targeted the genes included in the model allowed the identification of cell line specificities (Appendix File). For instance, target enrichment analysis showed that LNCaP cell lines are especially sensitive to drugs targeting PI3K/AKT/MTOR, hormone-related (AR targeting) and Chromatin (bromodomain inhibitors, regulating Myc) pathways (adjusted p-values from target enrichment: 0.001, 0.001 and 0.032, respectively, Appendix File, Table S1), which corresponds to the model predictions (Table 1). Also LNCaP cell line is more sensitive to drugs targeting model-identified nodes than to drugs targeting other proteins (Figure S27, Mann-Whitney p-value 0.00041) and this effect is specific for LNCaP cell line (Mann-Whitney p-values ranging from 0.0033 to 0.38 for other prostate cancer cell lines).

**Figure 6:**
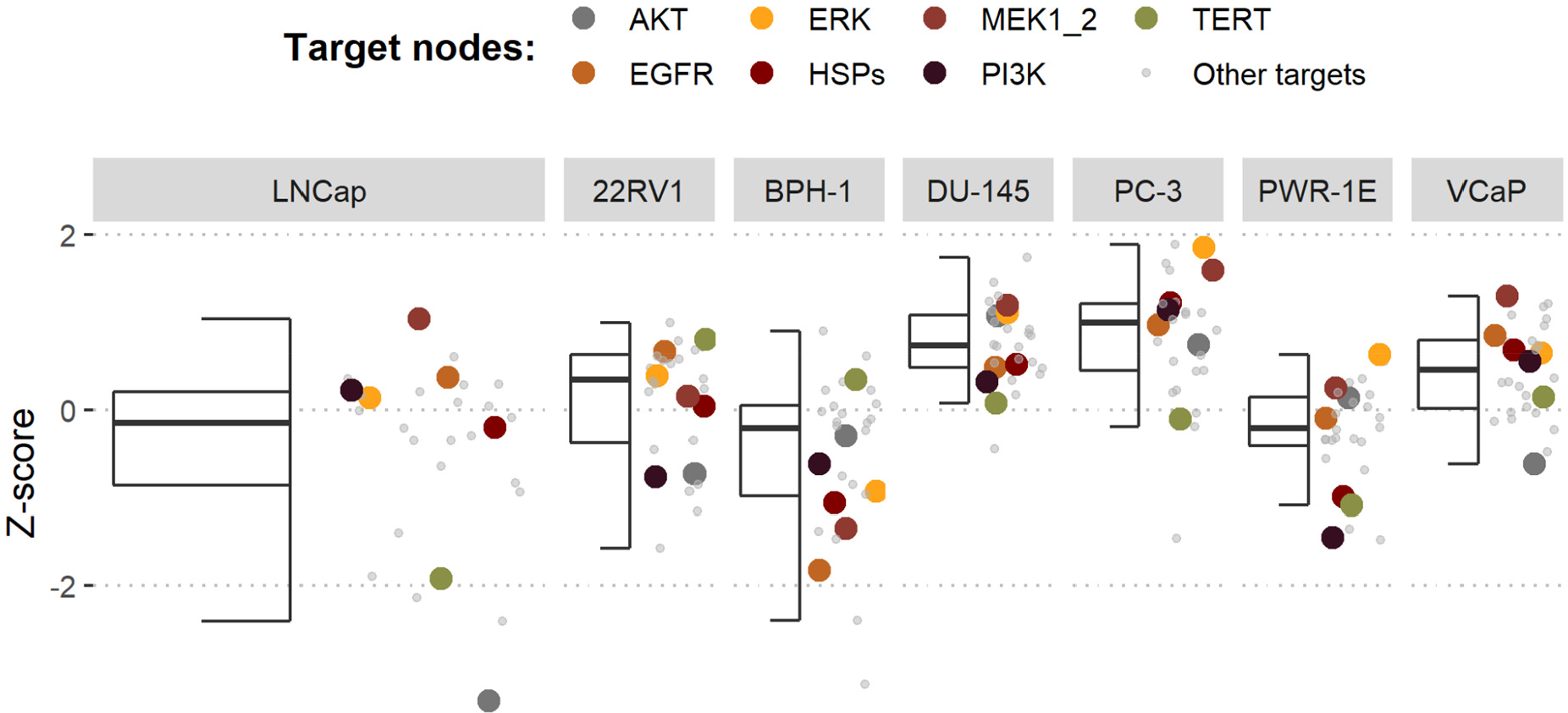
Model-targeting drugs’ sensitivities across prostate cell lines. GDSC z-score was obtained for all the drugs targeting genes included in the model for all the prostate cell lines in GDSC. Negative values means that the cell line is more sensitive to the drug. Drugs included in Table 1 were highlighted. “Other targets” are drugs targeting model-related genes that are not part of Table 1.

Overall, the drugs proposed through this analysis suggest the possibility to repurpose drugs that are used in treating other forms of cancer for prostate cancer and open the avenue for further experimental validations based on these suggestions.

#### Experimental validation of drugs in LNCaP

To validate the model predictions of the candidate drugs, we selected four drugs that target HSPs and PI3K and tested them in LNCaP cell line experiments by using endpoint cell viability measurement assays and real-time cell survival assays using the xCELLigence system (see Methods). The drug selection was a compromise between the drugs identified by our analyses (Table 1) and their effect in diminishing LNCaP’s proliferation (see previous section). In both assays, drugs that target HSP90AA1 and PI3K/AKT pathway genes retrieved from the model analyses were found to be effective against cell proliferation.

The Hsp90 chaperone is expressed abundantly and plays a crucial role in the correct folding of a wide variety of proteins such as protein kinases and steroid hormone receptors (Schopf *et al*, 2017). Hsp90 can act as a protector of less stable proteins produced by DNA mutations in cancer cells (Barrott & Haystead, 2013; Hessenkemper & Baniahmad, 2013). Currently, Hsp90 inhibitors are in clinical trials for multiple indications in cancer (Iwai *et al*, 2012; Le *et al*, 2017; Chen *et al*, 2019). The PI3K/AKT signalling pathway controls many different cellular processes such as cell growth, motility, proliferation, and apoptosis and is frequently altered in different cancer cells (Carceles-Cordon *et al*, 2020; Shorning *et al*, 2020). Many PI3K/AKT inhibitors are in different stages of clinical development and some of them are approved for clinical use (Table 1).

Notably, Hsp90 (NMS-E973,17-DMAG) and PI3K/AKT pathway (PI-103, Pictilisib) inhibitors showed a dose-dependent activity in the endpoint cell viability assay determined by the fluorescent resazurin after a 48-hour incubation (Figure 7).

**Figure 7:**
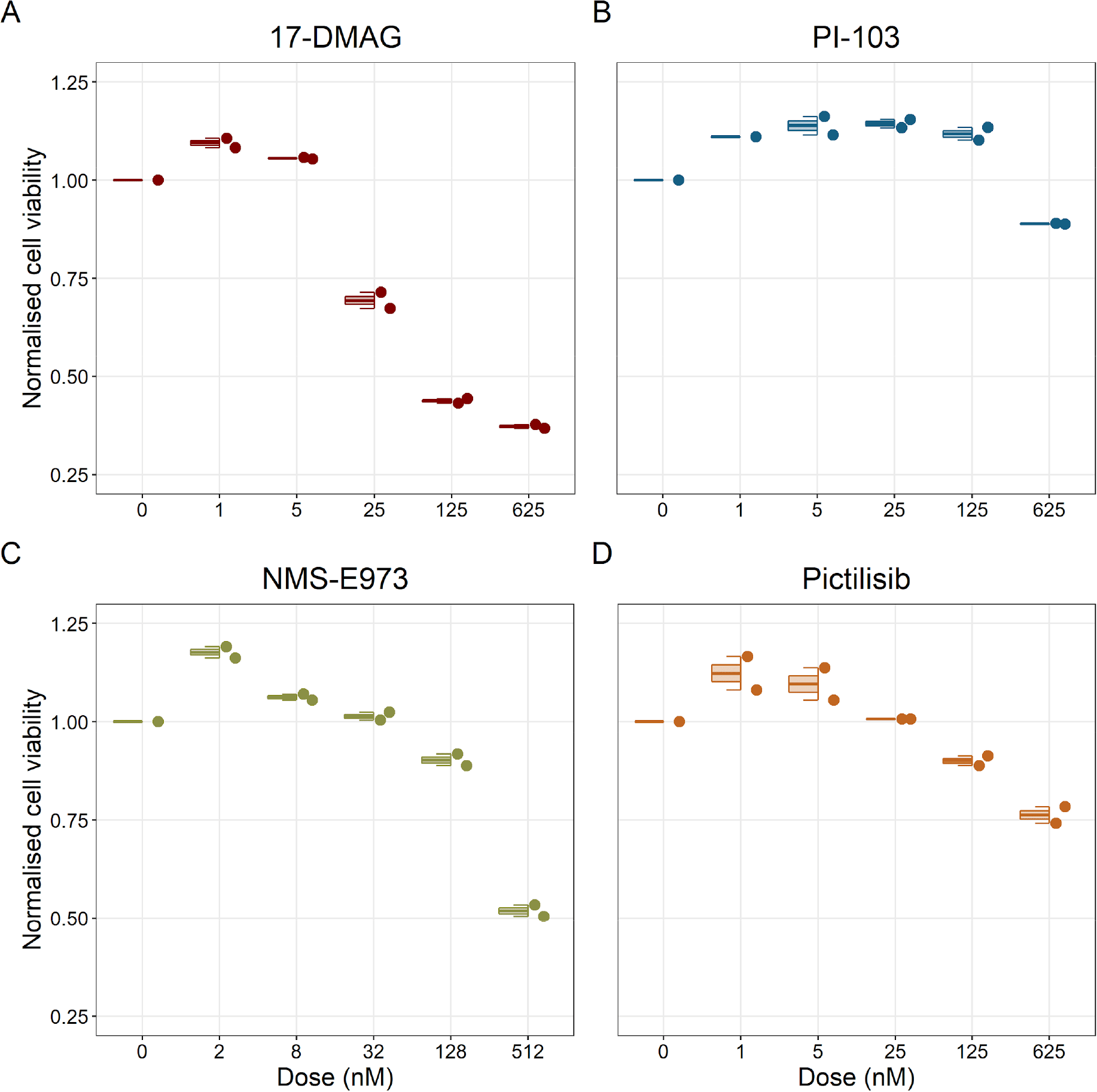
Cell viability assay determined by the fluorescent resazurin after a 48-hours incubation showed a dose-dependent response to different inhibitors. A) Cell viability assay of LNCaP cell line response to 17-DMAG HSP90 inhibitor. B) Cell viability assay of LNCaP cell line response to PI-103 PI3K/AKT pathway inhibitor. C) Cell viability assay of LNCaP cell line response to NMS-E973 HSP90 inhibitor. D) Cell viability assay of LNCaP cell line response to Pictilisib PI3K/AKT pathway inhibitor. Concentrations of drugs were selected to capture their drug-dose response curves. The concentrations for the NMS-E973 are different from the rest as this drug is more potent than the rest (see Material and methods).

We studied the real-time response of LNCaP cell viability upon drug addition and saw that the LNCaP cell line is sensitive to Hsp90 and PI3K/AKT pathway inhibitors (Figure 8 and 9, respectively). Both Hsp90 inhibitors tested, 17-DMAG and NMS-E973, reduced the cell viability 12 hours after drug supplementation (Figure 8A for 17-DMAG and Figure 8E for NMS-E973), with 17-DMAG having a stronger effect and in a more clear concentration-dependent manner than NMS-E973 (Figure 7B-D for 17-DMAG and Figure 7F-H for NMS-E973).

**Figure 8:**
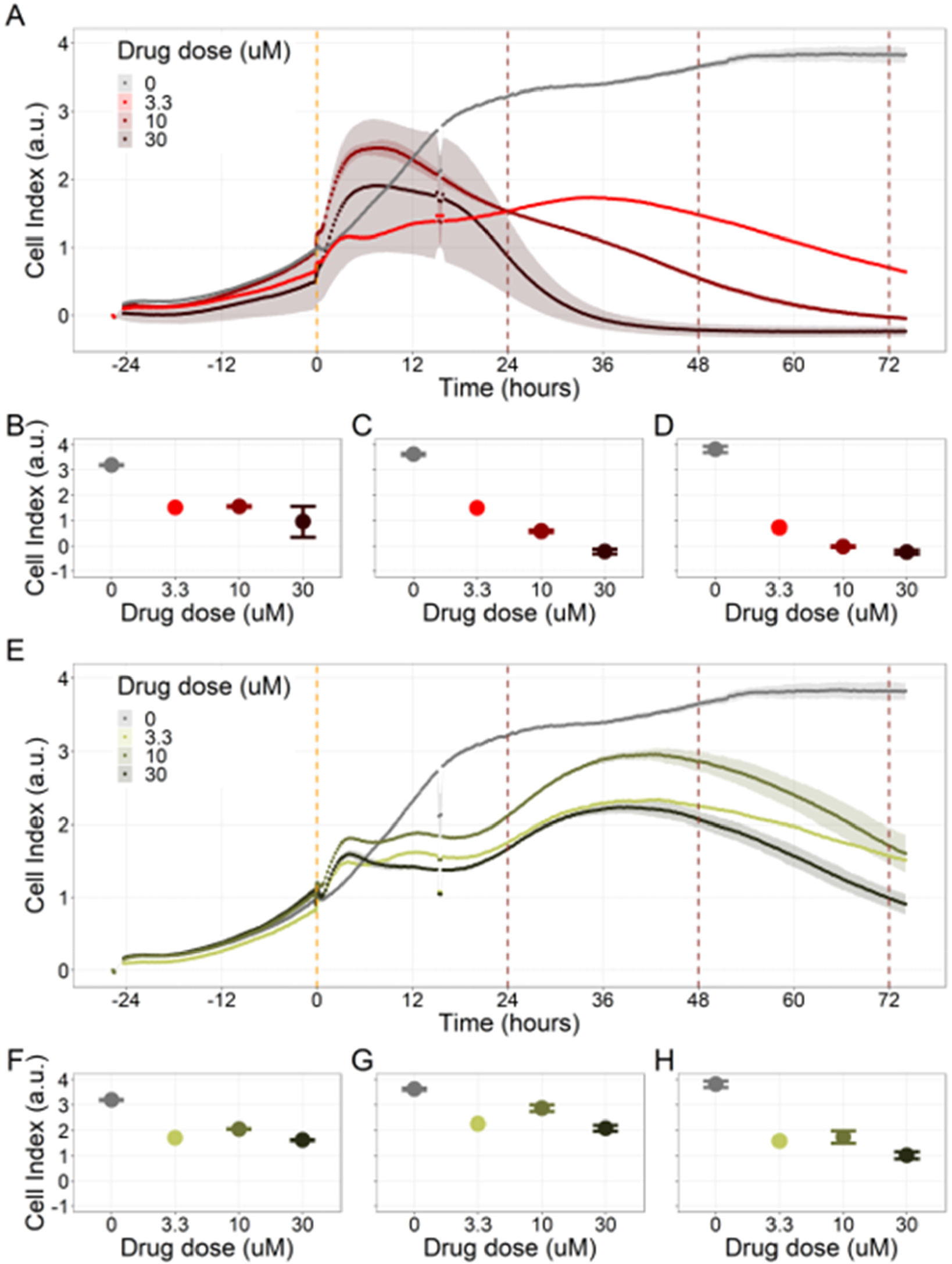
Hsp90 inhibitors resulted in dose-dependent changes in the LNCaP cell line. A) Real-time cell electronic sensing (RT-CES) cytotoxicity assay of Hsp90 inhibitor, 17-DMAG. The yellow dotted line represents 17-DMAG addition. The brown dotted lines are indicative of the cytotoxicity assay results at 24 hours (B), 48 hours (C) and 72 hours (D) after 17-DMAG addition. E) RT-CES cytotoxicity assay of Hsp90 inhibitor, NMS-E973. The yellow dotted line represents NMS-E973 addition. The brown dotted lines are indicative of the cytotoxicity assay results at 24 hours (F), 48 hours (G) and 72 hours (H) after NMS-E973 addition.

**Figure 9:**
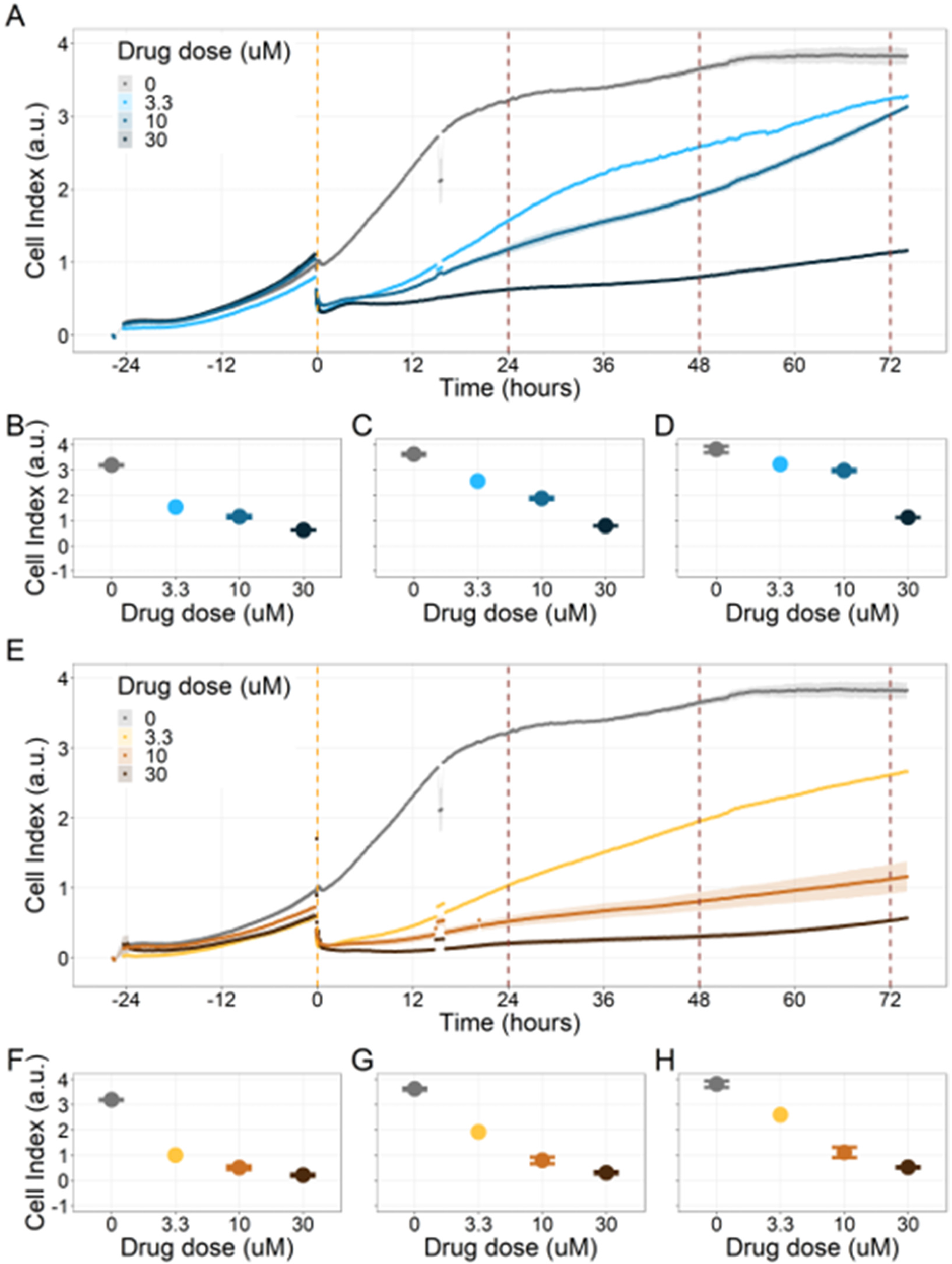
PI3K/AKT pathway inhibition with different PI3K/AKT inhibitors shows dose-dependent response in LNCaP cell line. A) Real-time cell electronic sensing (RT-CES) cytotoxicity assay of PI3K/AKT pathway inhibitor, PI-103. The yellow dotted line represents PI-103 addition. The brown dotted lines are indicative of the cytotoxicity assay results at 24 hours (B), 48 hours (C) and 72 hours (D) after PI-103 addition. E) RT-CES cytotoxicity assay of PI3K/AKT pathway inhibitor, Pictilisib. The yellow dotted line represents Pictilisib addition. The brown dotted lines are indicative of the cytotoxicity assay results at 24 hours (F), 48 hours (G) and 72 hours (H) after Pictilisib addition.

Likewise, both PI3K/AKT pathway inhibitors tested, Pictilisib and PI-103, reduced the cell viability immediately after drug supplementation (Figure 9A for Pictilisib and Figure 9E for PI-103), in a concentration-dependent manner (Figure 9B-D for Pictilisib and Figure 9F-H for PI-103). In addition, Hsp90 inhibitors had a more prolonged effect on the cells’ proliferation than PI3K/AKT pathway inhibitors.

## Discussion

Clinical assessment of cancers is moving towards more precise, personalised treatments, as the times of one-size-fits-all treatments are no longer appropriate, and patient-tailored models could boost the success rate of these treatments in clinical practice. In this study, we set out to develop a methodology to investigate drug treatments using personalised Boolean models. Our approach consists of building a model that represents the patient-specific disease status and retrieving a list of proposed interventions that affect this disease status, notably by reducing its pro-cancerous behaviours. In this work, we have showcased this methodology by applying it to TCGA prostate cancer patients and to GDSC prostate cancer cell lines, finding patient- and cell-line-specific targets and validating selected cell-line-specific predicted targets (Figure 1).

First, a prostate cancer Boolean model that encompasses relevant signalling pathways in cancer was constructed based on already published models, experimental data analyses and pathway databases (Figure 2). The influence network and the assignment of logical rules for each node of this network were obtained from known interactions described in the literature (Figure 3). This model describes the regulation of invasion, migration, cell cycle, apoptosis, androgen and growth factors signalling in prostate cancer (Appendix file).

Second, from this generic Boolean model, we constructed personalised models using the different datasets, i.e. 488 patients from TCGA and eight cell lines from GDSC. We obtained Gleason-score-specific behaviours for TCGA’s patients when studying their *Proliferation* and *Apoptosis* scores, observing that high *Proliferation* scores are higher in high Gleason groups (Figure 4). Thus, the use of these personalised models can help rationalise the relationship of Gleason grading with some of these phenotypes.

Likewise, GDSC data was used with the prostate model to obtain prostate-specific cell-line models (Figure 6). These models show differential behaviours, notably in terms of *Invasion* and *Proliferation* phenotypes (Figure S19). One of these cell-line-specific models was chosen, LNCaP, and the effects of all its genetic perturbations were thoroughly studied. We studied 32258 mutants, including single and double mutants, knock-out and over-expressed, and their phenotypes (Appendix File, Figure S25 and S26). 32 knock-out perturbations that depleted *Proliferation* and/or increased *Apoptosis* were identified and 16 of them were selected for further analyses (Table 1). The LNCaP-specific model was simulated using different initial conditions that capture different growth media’s specificities, such as RPMI media with and without androgen or epidermal growth factor (Appendix File, Figure S24).

Third, these personalised models were used to simulate the inhibition of druggable genes and proteins, uncovering new treatment’s combination and their synergies. We developed a methodology to simulate drug inhibitions in Boolean models, termed PROFILE_v2, as an extension of previous works (Béal *et al*, 2019). The LNCaP-specific model was used to obtain simulations with nodes and pairs of nodes corresponding to the genes of interest inhibited with varying strengths. This study allowed us to compile a list of potential targets (Table 1) and to identify potential synergies among genes in the model (Figure 5). Some of the drugs that targeted these genes, such as AKT and TERT, were identified in GDSC as having more sensitivity in LNCaP than in the rest of the prostate cancer cell lines (Figure 6). In addition, drugs that targeted genes included in the model allowed the identification of cell line specificities (Appendix File).

Fourth, we validated experimentally the effect of Hsp90 and PI3K/AKT pathway inhibitors on the LNCaP cell line, finding a concentration-dependent inhibition of the cell line viability as predicted, confirming the role of the drugs targeting these proteins in reducing LNCaP’s proliferation (Figure 7 and 8). Notably, these targets have been studied in other works on prostate cancer (Chen *et al*, 2019; Le *et al*, 2017).

The study presented here enables the study of drug combinations and their synergies. One reason for searching for combinations of drugs is that these have been described for allowing the use of lower doses of each of the two drugs reducing their toxicity (Bayat Mokhtari *et al*, 2017), evading compensatory mechanisms and combating drug resistances (Al-Lazikani *et al*, 2012; Krzyszczyk *et al*, 2018).

Even if this approach is attractive and promising, it has some limitations. First, the analyses performed with the mathematical model do not aim at predicting drug dosages *per se* but to help in the identification of potential candidates. The patient-specific changes in *Proliferation* and *Apoptosis* scores upon mutation are maximal theoretical yields that are used to rank the different potential treatments and should not be used as a direct target for experimental results or clinical trials. Our methodology suggests treatments for individual patients, but the obtained results vary greatly from patient to patient, which is not an uncommon issue of personalized medicine (Ciccarese *et al*, 2017; Molinari *et al*, 2018). This variability is an economical challenge for labs and companies to pursue true patient-specific treatments and also poses challenges in clinical trial designs aimed at validating the model based on the selection of treatments (Cunanan *et al*, 2017). Nowadays and because of these constraints, it might be more commercially interesting to target group-specific treatments, which can be more easily related to clinical stages of the disease.

Mathematical modelling of patient profiles helps to classify them in groups with differential characteristics, providing, in essence, a grade-specific treatment. We therefore based our analysis on clinical grouping defined by the Gleason grades, but some works have emphasized the difficulty to properly assess them (Chen & Zhou, 2016) and as a result may not be the perfect predictor for the patient subgrouping in this analysis, even though it is the only available one for these datasets. The lack of subgrouping that stratifies patients adequately may undermine the analysis of our results and could explain the *Proliferation* and *Apoptosis* scores of high-grade and low-grade Gleason patients.

Moreover, the behaviours observed in the simulations of the cell-lines-specific models do not always correspond to what is reported in the literature. The differences between simulation results and biological characteristics could be addressed in further studies by including other pathways, for example better describing the DNA repair mechanisms, or by tailoring the model with different sets of data, as the data used to personalise these models do not allow to cluster these cell lines according to their different characteristics (Appendix File, Figure S21 and S22). In this sense, another limitation is that we use static data (or a snapshot of dynamic data) to build dynamic models and to study its stochastic results. Thus, these personalised models would likely improve their performance if they were fitted to dynamic data (Saez-Rodriguez & Blüthgen, 2020) or quantitative versions of the models were built, such as ODE-based, that may capture more fine differences among cell lines.

The present work contributes to efforts aimed at using modeling (Rivas-Barragan *et al*, 2020; Eduati *et al*, 2020; Zañudo *et al*, 2017) and other computational methods (Madani Tonekaboni *et al*, 2018; Menden *et al*, 2019) for the discovery of novel drug targets and combinatorial strategies. Our study expands the prostate drug catalogue and improves predictions of the impact of these in clinical strategies for prostate cancer by proposing and grading the effectiveness of a set of drugs that could be used off-label or repurposed. The insights gained from this study present the potential of using personalised models to obtain precise, personalised drug treatments for cancer patients.

## Materials and Methods

### Data acquisition

Publicly available data of 489 human prostate cancer patients from TCGA described in (Hoadley *et al*, 2018) were used in the present work. We gathered mutations, CNA, RNA and clinical data from cBioPortal (https://www.cbioportal.org/study/summary?id=prad_tcga_pan_can_atlas_2018) for all of these samples resulting in 488 with complete omics datasets.

Publicly available data of cell lines used in the present work were obtained from Genomics of Drug Sensitivity in Cancer database (GDSC) (Iorio *et al*, 2016). Mutations, CNA and RNA data, as well as cell lines descriptors, were downloaded from (https://www.cancerrxgene.org/downloads).

All these data were used to personalise Boolean models using our PROFILE method (Béal *et al*, 2019).

### Prior knowledge network construction

Several sources were used in building this prostate Boolean model and in particular the model published by Fumiã and Martins (2013). This model includes several signalling pathways such as the ones involving receptor tyrosine kinase (RTKs), phosphatidylinositol 3-kinase (PI3K)/AKT, WNT/b-Catenin, transforming growth factor-b (TGF-b)/Smads, cyclins, retinoblastoma protein (Rb), hypoxia-inducible transcription factor (HIF-1), p53 and ataxia-telangiectasia mutated (ATM)/ataxia-telangiectasia and Rad3-related (ATR) protein kinases. The model includes these pathways as well as the substantial cross-talks among them. For a complete description of the process of construction, see Appendix File.

The model also includes several pathways that have a relevant role in our datasets identified by ROMA (Martignetti *et al*, 2016), a software that uses the first principal component of a PCA analysis to summarise the coexpression of a group of genes in the gene set, identifying significantly overdispersed pathways with a relevant role in a given set of samples. This software was applied on the TCGA transcriptomics data using the gene sets described in the Atlas of Cancer Signaling Networks, ACSN (Kuperstein *et al*, 2015) (www.acsn.curie.fr) and in Hallmarks (Liberzon *et al*, 2015) (Appendix File, Figure S1) and highlighted the signalling pathways that show high variance across all samples, suggesting candidate pathways and genes. Additionally, OmniPath (Türei *et al*, 2021) was used to extend the model and complete it connecting the nodes from Fumiã and Martins and the ones from ROMA analysis. OmniPath is a comprehensive collection of literature-curated human signalling pathways, which includes several databases such as Signor (Perfetto *et al*, 2016) or Reactome (Fabregat *et al*, 2018) and that can be queried using pypath, a Python module for molecular networks and pathways analyses.

Fusion genes are frequently found in human prostate cancer and have been identified as a specific subtype marker (The Cancer Genome Atlas Research Network, 2015). The most frequent is TMPRSS2:ERG as it involves the transcription factor ERG, which leads to cell-cycle progression. ERG fuses with the AR-regulated TMPRSS2 gene promoter to form an oncogenic fusion gene that is especially common in hormone-refractory prostate cancer, conferring androgen responsiveness to ERG. A literature search reveals that ERG directly regulates EZH2, oncogene c-Myc and many other targets in prostate cancer (Kunderfranco *et al*, 2010).

We modelled the gene fusion with activation of ERG by the decoupling of ERG in a special node *AR_ERG* that is only activated by *AR,* when the fused_event input node is active. In the healthy case, *fused_event* (that represents TMPRSS2:ERG fusion event) is fixed to 0 or inactive. The occurrence of the gene fusion is represented with the model perturbation where *fused_event* is fixed to 1. This *AR_ERG* node is further controlled by tumour suppressor NKX3-1 that accelerates *DNA_repair* response and avoids the gene fusion TMPRSS2:ERG. Thus, loss of NKX3-1 favours recruitment to the ERG gene breakpoint of proteins that promote error-prone non-homologous end-joining (Bowen *et al*, 2015).

The network was further documented using up-to-date literature and was constructed using GINsim (Chaouiya *et al*, 2012), which allowed us to study its stable states and network properties.

### Boolean model construction

We converted the network to a Boolean model by defining a regulatory graph, where each node is associated with discrete levels of activity (0 or 1). Each edge represents a regulatory interaction between the source and target nodes and is labelled with a threshold and a sign (positive or negative). The model is completed by logical rules (or functions), which assign a target value to each node for each regulator level combination (Chaouiya *et al*, 2012; Abou-Jaoudé *et al*, 2016). The regulatory graph was constructed using GINsim software (Chaouiya *et al*, 2012) and then exported in a format readable by MaBoSS software (see below) in order to perform stochastic simulations on the Boolean model.

The final model has a total of 133 nodes and 449 edges (SuppFile 1) and includes pathways such as androgen receptor and growth factor signalling, several signalling pathways (Wnt, NFkB, PI3K/AKT, MAPK, mTOR, SHH), cell cycle, epithelial-mesenchymal transition (EMT), Apoptosis, DNA damage, etc. This model has 9 inputs (*EGF, FGF, TGF beta, Nutrients, Hypoxia, Acidosis, Androgen, TNF alpha* and *Carcinogen* presence) and 6 outputs (*Proliferation*, *Apoptosis, Invasion, Migration,* (bone) *Metastasis* and *DNA repair*). This model was deposited in the GINsim Database with identifier 252 (http://ginsim.org/model/signalling-prostate-cancer) and in BioModels (Malik-Sheriff *et al*, 2019) with identifier MODEL2106070001 (https://www.ebi.ac.uk/biomodels/MODEL2106070001). SuppFile 1 is provided as a zipped folder with the model in several formats: MaBoSS, GINsim, SBML as well as images of the networks and its annotations.

### Stochastic Boolean model simulation

MaBoSS (Stoll *et al*, 2012, 2017) is a C++ software for stochastically simulating continuous/discrete-time Markov processes defined on the state transition graph (STG) describing the dynamics of a Boolean model (for more details, see (Chaouiya *et al*, 2012; Abou-Jaoudé *et al*, 2016)). MaBoSS associates transition rates to each node’s activation and inhibition, enabling it to account for different time scales of the processes described by the model. Probabilities to reach a phenotype (to have value ON) are thus computed by simulating random walks on the probabilistic STG. Since a state in the STG can combine the activation of several phenotypic variables, not all phenotype probabilities are mutually exclusive (like the ones in Appendix File, Figure S25). Using MaBoSS we can study an increase or decrease of a phenotype probability when the model variables are altered (nodes status, initial conditions and transition rates), which may correspond to the effect of particular genetic or environmental perturbation. In the present work, the outputs of MaBoSS focused on the readouts of the model, but this can be done for any node of a model.

MaBoSS applies Monte-Carlo kinetic algorithm (i.e. Gillespie algorithm) to the STG to produce time trajectories (Stoll *et al*, 2012, 2017) so time evolution of probabilities are estimated once a set of initial conditions are defined and a maximum time is set to ensure that the simulations reach asymptotic solutions. Results are analyzed in two ways: (1) the trajectories for particular model states (states of nodes) can be interpreted as the evolution of a cell population as a function of time and (2) asymptotic solutions can be represented as pie charts to illustrate the proportions of cells in particular model states. Stochastic simulations with MaBoSS have already been successfully applied to study several Boolean models (Calzone *et al*, 2010; Remy *et al*, 2015; Cohen *et al*, 2015).

### Data tailoring the Boolean model

Logical models were tailored to a dataset using PROFILE to obtain personalised models that capture the particularities of a set of patients (Béal *et al*, 2019) and cell lines (Béal *et al*, 2021). Proteomics, transcriptomics, mutations and CNA data can be used to modify different variables of the MaBoSS framework such as node activity status, transition rates and initial conditions. The resulting ensemble of models is a set of personalised variants of the original model that can show great phenotypic differences. Different recipes (use of a given data type to modify a given MaBoSS variable) can be tested to find the combination that better correlates to a given clinical or otherwise descriptive data.

In the present case, TCGA-patient-specific models were built using mutations, CNA and/or RNA expression data. After studying the effect of these recipes in the clustering of patients according to their Gleason grouping (Appendix File, Figure S8-S12), we chose to use mutations and CNA as discrete data and RNA expression as continuous data.

Likewise, we tried different personalisation recipes to personalise the GDSC prostate cell lines models, but as they had no associated clinical grouping features, we were left with the comparison of the different values for the model’s outputs among the recipes (Appendix File, Figure S20). We used mutation data as discrete data and RNA expression as continuous data as it included the most quantity of data and reproduced the desired results (Figure S20). We decided not to include CNA as discrete data as it forced LNCAP proliferation to be zero, by forcing E2F1 node to be 0 and SMAD node to be 1 throughout the simulation (for more details, refer to Appendix File).

More on PROFILE’s methodology can be found in its own work (Béal *et al*, 2019) and at its dedicated GitHub repository: https://github.com/sysbio-curie/PROFILE.

### High-throughput mutant analysis of Boolean models

MaBoSS allows the study of knock-out or loss-of-function (node forced to 0) and gain-of-function (node forced to 1) mutants as genetic perturbations and of initial conditions as environmental perturbations. Phenotypes’ stabilities against perturbations can be studied and allow to determine driver mutations that promote phenotypic transitions (Montagud *et al*, 2017).

Genetic interactions were thoroughly studied using our pipeline of computational methods for Boolean modelling of biological networks (available at https://github.com/sysbio-curie/Logical_modelling_pipeline). LNCaP-specific Boolean model was used to perform single and double knock-out (node forced to 0) and gain-of-function (node forced to 1) mutants for each one of the 133 nodes, resulting in a total of 32258 models. These were simulated under the same initial conditions, their phenotypic results were collected and a PCA was applied on the wild-type-centred matrix.

The 488 TCGA-patient-specific models were studied in a similar way, but only perturbing 16 nodes shortlisted for their therapeutic target potential (AKT, AR, Caspase8, cFLAR, EGFR, ERK, GLUT1, HIF-1, HSPs, MEK1_2, MYC_MAX, p14ARF, PI3K, ROS, SPOP and TERT). Then, the nodes that mostly contributed to a decrease of *Proliferation* (Appendix File, Figure S17) or an increase in *Apoptosis* (Appendix File, Figure S18) were gathered from the 488 models perturbed.

Additionally, the results of LNCaP model’s double mutants were used to quantify the level of genetic interactions (epistasis or otherwise (Drees *et al*, 2005)) between two model genetic perturbations (resulting from either the gain-of-function mutation of a gene or from its knock-out or loss-of-function mutation) with respect to wild type phenotypes’ probabilities (Calzone *et al*, 2015). The method was applied to the LNCaP model studying *Proliferation* and *Apoptosis* scores (Appendix File, Figure S31 and S32).

This genetic interaction study uses the following equation for each gene pairs, which is equation 2 in Calzone *et al*, (2015):

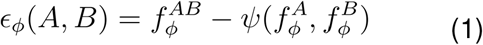

Where 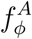 and 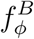 are phenotype ɸ fitness values of single gene defects, 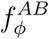 is the phenotype ɸ fitness of the double mutant, and *ψ*(*x*,*y*) is one of the four functions:

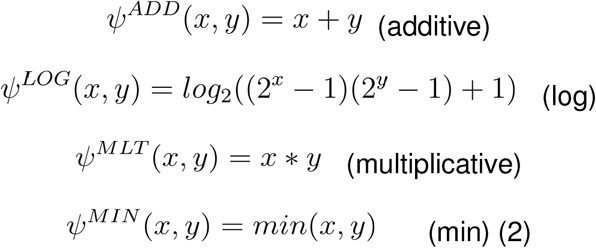

To choose the best definition of *ψ*(*x*,*y*), the Pearson correlation coefficient is computed between the fitness values observed in all double mutants and estimated by the null model (more information on (Drees *et al*, 2005)). Regarding 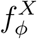 fitness value, to a given phenotype, 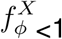 represents deleterious, 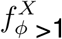 beneficial and 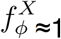 neutral mutation.

### Drug simulations in Boolean models

Logical models can be used to simulate the effect of therapeutic interventions and predict the expected efficacy of candidate drugs on different genetic and environmental backgrounds by using our PROFILE_v2 methodology. MaBoSS can perform simulations changing the proportion of activated and inhibited status of a given node. This can be determined in the configuration file of each model (see, for instance, “istate” section of the CFG files in SuppFiles 1, 3 and 5). For instance, out of 1000 trajectories of the Gillespie algorithm, MaBoSS can simulate 50% of them with an activated *AKT* and 50% with an inhibited *AKT* node. The phenotypes’ probabilities for the 1000 trajectories are averaged and these are considered to be representative of a model with a drug that half-inhibits the activity of *AKT*.

In present work, LNCaP model has been simulated with different levels of node activity, with 100% of node activity (no inhibition), 80%, 60%, 40%, 20% and 0% (proper knock-out), under four different initial conditions, a nutrient-rich media that simulates RPMI Gibco® media with DHT (androgen), with EGF, with both and with none. In terms of the model, the initial conditions are *Nutrients* is ON and *Acidosis*, *Hypoxia*, *TGF beta*, *Carcinogen* and *TNF alpha* are set to OFF. *EGF* and *Androgen* values vary upon simulations.

Drug synergies have been studied using Bliss Independence. The Combination Index was calculated with the following equation (Foucquier & Guedj, 2015):

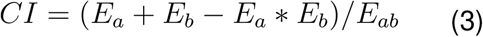

Where *E_a_* and *E_b_* is the efficiency of the single drug inhibitions and *E_ab_* is the inhibition resulting from the double drug simulations. A Combination Index (*CI*) below 1 represents synergy among drugs.

This methodology can be found in its own repository: https://github.com/ArnauMontagud/PROFILE_v2

### Identification of drugs associated with proposed targets

To identify drugs that could act as potential inhibitors of the genes identified with our models (Table 1), we explored the drug-target associations in DrugBank (Wishart et al, 2018). For those genes with multiple drug-target links, only those drugs that are selective and known to have relevance in various forms of cancer are considered here.

In addition to DrugBank searches, we also conducted exhaustive searches in ChEMBL (Gaulton *et al*, 2017) (http://doi.org/10.6019/CHEMBL.database.23) to suggest potential candidates for genes, whose information is not well documented in Drug Bank. From the large number of bioactivities extracted from ChEMBL, we filtered human data and considered only those compounds, whose bioactivities fall within a specific threshold (IC50/Kd/ Ki<100 nM).

We performed a target set enrichment analysis using the *fgsea* method (Korotkevich *et al*, 2016) from the *piano* R package (Väremo *et al*, 2013). We targeted pathway information from the GDSC1 and GDSC2 studies (Iorio *et al*, 2016) as target sets and performed the enrichment analysis on the normalised drug sensitivity profile of the LNCaP cell line. We normalised drug sensitivity across cell lines in the following way: cells were ranked from most sensitive to least sensitive (using ln(IC50) as drug sensitivity metrics), and the rank was divided by the number of cell lines tested with the given drug. Thus, the most sensitive cell line has 0, while the most resistant cell line has 1 normalised sensitivity. This rank-based metric made it possible to analyse all drug sensitivities for a given cell line, without drug-specific confounding factors, like mean IC50 of a given drug, etc.

### Cell culture method

For in vitro drug perturbation validations we used the androgen-sensitive prostate adenocarcinoma cell line LNCaP purchased from American Type Culture Collection (ATCC, Manassas, WV, USA). Cells were maintained in RPMI-1640 culture media (Gibco, Thermo Fisher Scientific, Waltham, MA, USA) containing 4.5 g/L glucose, 10% fetal bovine serum (FBS, Gibco), 1X GlutaMAX (Gibco) 1% PenStrep antibiotics (Penicillin G sodium salt, and Streptomycin sulfate salt, Sigma-Aldrich, St. Louis, MI, USA). Cells were maintained in a humidified incubator at 37 °C 5% CO2 (Sanyo, Osaka, Japan).

### Drugs used in the cell culture experiments

We tested two drugs targeted at Hsp90 and two targeted at PI3K complex. 17-DMAG is an Hsp90 inhibitor with an IC50 of 62 nM in a cell-free assay (Pacey *et al*, 2011). NMS-E973 is an Hsp90 inhibitor with DC50 of <10 nM for Hsp90 binding (Fogliatto *et al*, 2013). Pictilisib is an inhibitor of PI3Kα/δ with IC50 of 3.3 nM in cell-free assays (Zhan *et al*, 2017). PI-103 is a multi-targeted PI3K inhibitor for p110α/β/δ/γ with IC50 of 2 to 3 nM in cell-free assays and less potent inhibitor to mTOR/DNA-PK with IC50 of 30 nM (Raynaud *et al*, 2009). All drugs were obtained from commercial vendors and added to the growth media to have concentrations of 2, 8, 32, 128 and 512 nM for NMS-E973 and 1, 5, 25, 125 and 625 nM for the rest of the drugs in the endpoint cell viability and of 3.3, 10, 30 mM for all the drugs in the RT-CES cytotoxicity assay.

### Endpoint cell viability measurements

In vitro toxicity of the selected inhibitors was determined using the viability of LNCaP cells, determined by the fluorescent resazurin (Sigma-Aldrich, Germany) assay as described previously (Szebeni *et al*, 2017). Briefly, the LNCaP cells (10000) were seeded into 96-well plates (Corning Life Sciences, Tewksbury, MA, USA) in 100 μl RPMI media and incubated overnight. Test compounds were dissolved in dimethyl sulfoxide (DMSO, Sigma-Aldrich, Germany). Cells were treated with an increasing concentration of test compounds. The highest applied DMSO content of the treated cells was 0.4%. Cell viability was determined after 48 hours incubation. Resazurin reagent (Sigma–Aldrich, Budapest, Hungary) was added at a final concentration of 25 μg/mL. After 2 hours at 37°C 5%, CO2 (Sanyo) fluorescence (530 nm excitation/580 nm emission) was recorded on a multimode microplate reader (Cytofluor4000, PerSeptive Biosystems, Framingham, MA, USA). Viability was calculated with relation to blank wells containing media without cells and to wells with untreated cells.

### Real-time cell electronic sensing (RT-CES) cytotoxicity assay

Real-time cytotoxicity assay was performed as previously described (Ozsvári *et al*, 2010). Briefly, RT-CES 96-well E-plate (BioTech Hungary, Budapest, Hungary) was coated with gelatin solution (0.2% in PBS, phosphate buffer saline) for 20 min at 37 °C, then gelatin was washed twice with PBS solution. Growth media (50 μL) was then gently dispensed into each well of the 96-well E-plate for background readings by the RT-CES system prior to the addition of 50 μL of the cell suspension containing 2x10^4^ LNCaP cells. Plates were kept at room temperature in a tissue culture hood for 30 min prior to insertion into the RT-CES device in the incubator to allow cells to settle. Cell growth was monitored overnight by measurements of electrical impedance every 15 min. Continuous recording of impedance in cells was reflected by the cell index value. The next day cells were co-treated with different drugs. Treated and control wells were dynamically monitored over 72 h by measurements of electrical impedance every 5 min. Each treatment was repeated in 2 wells per plate during the experiments.

## Acknowledgements

The authors acknowledge the help provided by Jelena Čuklina at ETH Zurich, Vincent Noël at Institut Curie, Annika Meert at BSC and Aurélien Naldi at INRIA Saclay.

The authors acknowledge the technical expertise and assistance provided by the Spanish Supercomputing Network (Red Española de Supercomputación), as well as the computer resources used: the LaPalma Supercomputer, located at the Instituto de Astrofísica de Canarias and MareNostrum4, located at the Barcelona Supercomputing Center.

This work has been partially supported by the European Commission under the PrECISE project (H2020-PHC-668858), the INFORE project (H2020-ICT-825070) and the PerMedCoE (H2020-ICT-951773).

## Author contributions

Conceptualization: LC, JSR, EB

Data curation: AM, LT, PT, VS, RA

Formal Analysis: AM, JB, VS, BS, RA

Funding acquisition: LC, JSR, EB, LP, AV

Investigation: AM, JB, LT, PT, VS, RA

Methodology: AM, JB, PT, BS

Project administration: AM, LC, JSR, EB, LP

Resources: AM, PT, LT, RA, LP

Software: AM, JB

Supervision: LC, JSR, EB, LP

Validation: AM, BS, RA, LP

Visualization: AM, BS, JB, LC

Writing – original draft: AM, LC, PT, LT, JSR

Writing – review & editing: all authors

## Conflict of interest

LT is a full-time employee and shareholder of AstraZeneca. LP is a scientific advisor and RA is CEO of Astridbio Technologies Ltd. VS is a full-time employee of AstraZeneca. JSR receives funding from GSK and Sanofi and consultant fees from Travere Therapeutics. The other authors declare no conflicts of interest.

## Supplementary materials

SuppFile 1, a zipped folder with the model in several formats: MaBoSS, GINsim, SBML as well as images of the networks and its annotations.

SuppFile 2, a jupyter notebook to inspect Boolean models using MaBoSS.

SuppFile 3, a zipped folder with the TCGA-specific personalised models and their *Apoptosis* and *Proliferation* phenotype scores.

SuppFile 4, a TSV file with the *Apoptosis* and *Proliferation* phenotype scores of TCGA-patient-specific mutations.

SuppFile 5, a zipped folder with the cell-lines-specific personalised models.

SuppFile 6, a TSV file with the *Apoptosis* and *Proliferation* phenotype scores of LNCaP-cell-line specific mutations.

## Notes

https://github.com/ArnauMontagud/PROFILE_v2

