## Appendix File for "Patient-specific Boolean models of signaling networks guide personalized treatments"


**Prostate Boolean model construction** **2**

Prior knowledge network construction 2

Definition of inputs and outputs 2

Identification of new components based on literature search: 3

Identification of new components/pathways based on data analysis 4

Model extension with Omnipath via pypath 6

Model extension with the literature 7

Boolean model construction 10

**Boolean model simulation** **10**

Wild type simulation 10

Mutants simulation 11

Single mutations 12

Multiple mutations 12

**Personalised Boolean models of TCGA patients** **14**

Phenotype distribution of TCGA patients 14

Analysis of drugs that inhibit the activity of genes of TCGA patients 22

**Personalised Boolean models of prostate cell lines** **24**

**Personalised LNCaP Boolean model** **29**

High-throughput mutant analysis of LNCaP model 31

**Drug studies in prostate cell lines** **33**

Drugs associated to genes included in the model 33

Drugs associated to proposed targets of LNCaP 37

Gradual inhibition of genes in LNCaP model 38

Single inhibitions 39

Double inhibitions 41

**References** **45**

### Prostate Boolean model construction

Building the model is done in three steps :

1. Identifying signalling pathways or particular genes and proteins that are especially relevant to describe the prostate cancer tumorigenesis and tumor growth. Most of them are components that are known to be frequently altered in cancers.
2. Building a regulatory network that includes simplified representations of pathways identified as relevant for prostate cancer, as well as all individually identified genes. Each pathway is characterized by the key players that regulate it. This network takes the form of a directed graph for which positive and negative influences between components are represented.
3. From this network, a logical model is derived describing the network dynamics in specific contexts (dependent on initial conditions or perturbations). To this end, logical rules are associated with each node of the network to indicate how it is activated or inhibited by different combinations of its regulators.

#### Prior knowledge network construction

We started by using a published logical model of human signalling network (Fumiã & Martins, 2013), which is based on integrated experimental evidence of signal transduction. This model integrates major signaling pathways that have a role in regulating cell death and proliferation in many tumors. They include those involving receptor tyrosine kinase (RTKs), phosphatidylinosital 3-kinase (PI3K)/AKT, WNT/b-Catenin, transforming growth factor-b (TGF-b)/Smads, cyclins, retinoblastoma protein (Rb), hypoxia-inducible transcription factor (HIF-1), p53 and ataxia-telangiectasia mutated (ATM)/ataxia-telangiectasia and Rad3-related (ATR) protein kinases. The pathways reveal substantial cross-talks.

This initial generic network was then extended to include prostate-cancer-specific genes and proteins using several approaches presented below.

##### Definition of inputs and outputs

Our Boolean model aims at predicting prostate phenotypic behaviours for healthy and cancer cells in different ‘‘physiological’’ conditions. To account for these conditions, we considered 9 inputs that represent different physiological conditions of interest. These are *EGF, FGF, TGF beta, Nutrients, Hypoxia, Acidosis androgen, TNF alpha and Carcinogen* presence. These input nodes have no regulation and their values are fixed for each simulation, representing the cell’s microenvironmental characteristics.

For simplicity, we choose to clearly define phenotype variables as output nodes allowing the integration of multiple phenotypic signals and obtaining a 0/1 value for each phenotype. Our model has a total of 11 outputs. We define three main phenotypes representing the growing status of the cell: *Proliferation*, *Apoptosis* and *Quiescence*. *Apoptosis* is activated by Caspase 8 or Caspase 9, while *Proliferation* is activated by cyclins from the cell cycle. We define *Quiescence* as the absence of *Proliferation* and *Apoptosis* and these two, although not directly linked, are always mutually exclusive in simulations.

The proliferation output is sometimes described in already published models as specific stationary protein activation patterns, namely the following sequence of activation of cyclins: Cyclin D, then Cyclin E, then Cyclin A, then Cyclin B. This sequence can easily be detected in complex attractors in synchronous dynamics. However, since asynchronous dynamics was chosen for this work and it is more difficult to analyze complex attractors with it, we define *Proliferation* as activated by either of the four cyclins. Transient dynamics in MaBoSS simulations allow us to check the correct oscillation of cyclins.

Furthermore, we define several phenotypic outputs that are not mutually exclusive but detect the activation of some markers of cancer hallmarks: *Invasion, Migration,* (bone) *Metastasis,* and *DNA repair*.

#### Identification of new components based on literature search:

Several studies have focused on identifying main subtypes among the heterogeneous molecular abnormalities in prostate cancer. In particular, a TCGA study (The Cancer Genome Atlas Research Network, 2015) reported a comprehensive molecular analysis of 333 primary prostate carcinomas. Seven subtypes, containing 74% of these tumors, were defined by specific gene fusions (ERG, ETV1/4 and FLI1) or mutations (SPOP, FOXA1 and IDH1). Epigenetic profiles allowed us to identify a methylator phenotype in the IDH1 mutant subset. SPOP and FOXA1 mutant tumors show the highest levels of AR-induced transcripts. Lesions in the PI3K or MAPK signaling pathways are observed in 25% of the prostate cancers and DNA repair genes inactivation in 19%.

The following list of frequently mutated genes extracted from this study indicate components that could be included in the model, provided that enough information is available on their mechanistic roles:

- gene fusions: ERG, ETV1, ETV4, FLI1
- deletions: SPOP, FOXA1, IDH1, TP53, PTEN, PIK3CA, BRAF, CTNNB1, HRAS, MED12, ATM, CDKN1B, RB1, NKX3-1, AKT1, ZMYM3, KMT2C, KMT2D, ZNF595, CHD1, BRCA2, CDK12, SPINK1
- amplifications: CCND1, MYC, FGFR1, WHSC1L1.

Comparing with a published cohort of 150 castration-resistant metastatic prostate cancer samples (Robinson *et al*, 2015), the authors find a similar subtype distribution as in (Abeshouse *et al*, 2015), with increased alteration rates in the metastatic samples and more frequent amplification or mutation of AR, as well as DNA repair and PI3K pathway alterations.

Other studies such as (Altieri *et al*, 2009) focus on the role of specific pathways which play a critical role in prostate cancer maintenance, such as chaperone-mediated mitochondrial homeostasis (in particular with HSP90 found very abundant in prostate cancer), integrin-dependent cell signaling and RUNX2-regulated gene expression in the metastatic bone microenvironment.

Notably, a set of regulatory maps of signalling pathway maps and altered circuitries of various cell biological events associated with the pathogenesis of human prostate cancer have been published recently (Datta *et al*, 2016). The authors manually constructed networks based on the literature. These networks constitute an important resource for retrieving information on prostate cancer specific components. Although not exhaustive, these maps are synthetic pictures of the existing knowledge on molecular events involved in prostate cancer hallmarks.

The covered hallmarks include: (1) classical cancer hallmarks: insensitivity to anti-growth signal, self-sufficiency in growth signal, tumor promoting inflammation, genome instability, mutation and perturbation, angiogenesis, metastasis, cell death resistance, metabolic reprogramming, avoidance of immune destruction, enabling replicative immortality, tumour microenvironment; and (2) prostate cancer specific hallmarks: androgen receptor signalling androgen independence, castration resistance.

This study points toward some candidate nodes to extend our network in order to take into account, at least in a simplified way, most pathways present in the maps. In particular, it shows that the initial network obtained through combinations of published models ignore any pathways related to inflammation, metabolism, immune evasion, or the tumor microenvironment. However, the resource contains few mechanistic details for the interactions between its components, which are a mix of genes, proteins, molecules, processes and phenotypes.

Finally, among all these genes associated with prostate cancer, a subset has been chosen for further research: AR, PTEN, SPOP, TP53, EZH2, FOXA1, BRCA1, BRCA2, PIK3CA, AKT1, NCOA2, NCOR1, NCOR2, EP300, MYC, RB1, CHD1, CDKN1B, MED12, ZNF595, HOXB13.

#### Identification of new components/pathways based on data analysis

ROMA (Martignetti *et al*, 2016) is a software package written in Java for the quantification and representation of biological module activity using expression data. It uses the first principal component of a PCA analysis to summarize the coexpression of a group of genes in the gene set.

We apply ROMA analysis on the transcriptomics data of TCGA. We define gene sets as they are described in the atlas of cancer signaling networks, ACSN (Kuperstein *et al*, 2015) (www.acsn.curie.fr) and in the Hallmarks (Liberzon *et al*, 2015). ACSN is centered on signaling pathways such as DNA repair, cell death, EMT, cell adhesion, cell cycle, etc. and the Hallmarks gene sets provide a list of genes that participate in biological processes integrating information from other pathway databases.

Using ROMA, we are able to identify some pathways significantly overdispersed over the samples that should have relevant roles in prostate cancer and need therefore to be correctly described in the model.

The results show that, for ACSN database, among the 140 pathways from the database, 65 modules reveal a high variance of protein expression across all samples (Figure S1). The gene sets linked to the cell cycle seem to show a progressive activation from normal to high grade tumours, so does the DNA repair pathway with differences in the mechanisms that participate in DNA repair, whereas some gene sets such as the one related to immunosuppressive cytokine pathway show opposite behavior. We performed the same analysis with the Hallmarks database and found 16 out of the 50 pathways that showed high variance. We can confirm the role of the cell cycle in tumour progression (E2F_targets and G2_M checkpoints).

Note that in both analyses, we see that group 3 is the most heterogeneous group, with a score that does not always follow the trend of increasing or decreasing pathway scores from group 1 to group 5.

ROMA provides some hints on where to extend the network to fully grab the alterations that are found in prostate cancer patients. For instance, the Hedgehog pathway was not described in the already published logical models that we used as a starting point of this model. Moreover, both the cell cycle and the DNA repair pathways were overly simplified, and were thus extended in this version.

Some pathways related to the immune response seem to be highly represented in ACSN and would need to be included in future extended versions of the prostate network, probably in the form of interacting networks of different cell types.

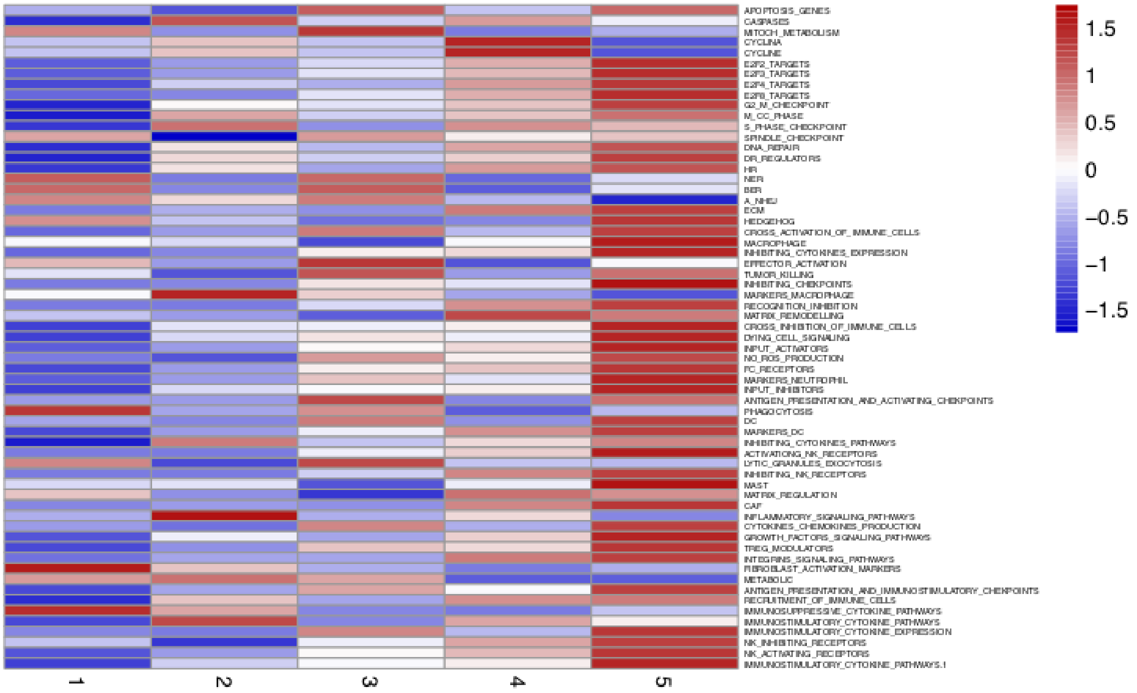


(A)

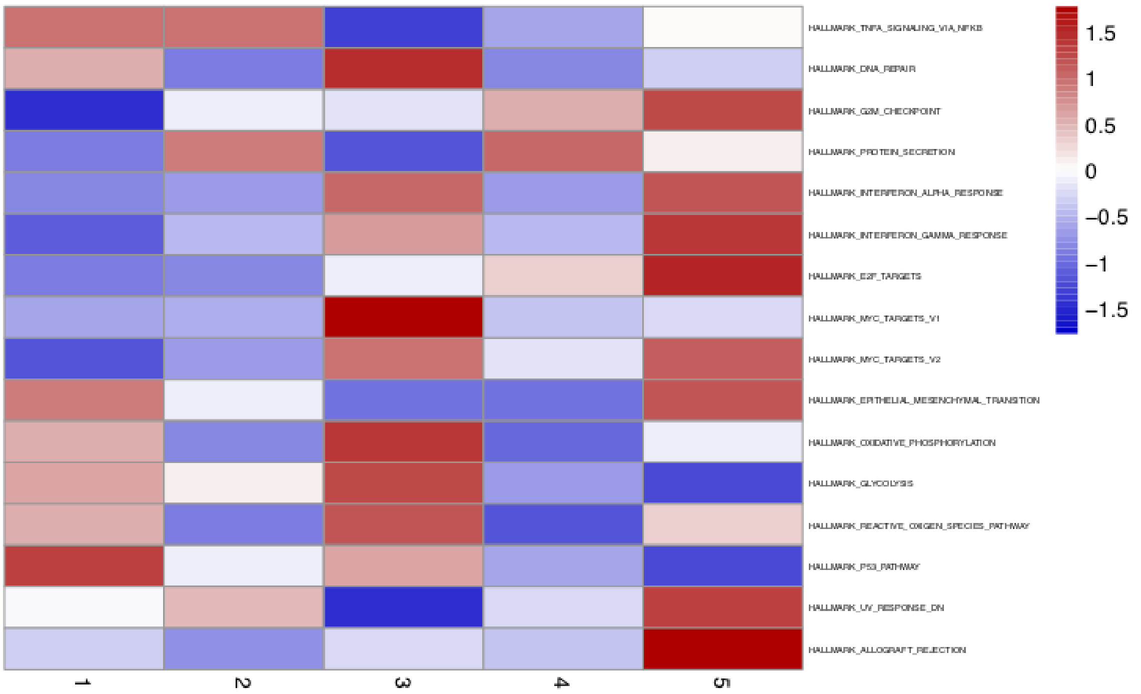


(B)

Figure S1: Mean activities by subgroups for gene modules defined from pathways described in ACSN (A) and in Hallmarks' gene sets (B) and that are significantly overdispersed over all samples. Blue indicates low pathway activity, red indicates high pathway activity.

#### Model extension with Omnipath via pypath

OmniPath (Türei *et al*, 2016) is a comprehensive collection of high confidence, literature curated, human signaling pathways. It is accompanied and developed together with Pypath, a Python module for cellular signaling pathways analysis.

Pypath is a python module used to query the content of Omnipath in order to retrieve components and interactions in the human protein-protein signaling network associated with annotations, especially sources, literature references, direction, effect signs (stimulation/inhibition) and enzyme-substrate interactions.

The development of pypath allows us to build personalised queries. For instance, existing interaction paths between a protein of interest and a list of user-defined proteins can be found, with a given size for the paths. We use this in the extension process of our network to automatically find new interactions between a new gene and the genes already included in the network. We filter the interactions found to select the ones for which the direction and sign are known.

For example, when extending the network with the chaperone protein HSP90AA1, we generate the graph displayed in Figure S2, which shows all signed directed interactions linking HSP90AA1 to the network. The associated references given as annotations are useful to check the mechanism behind each interaction and manually infer a logical rule.

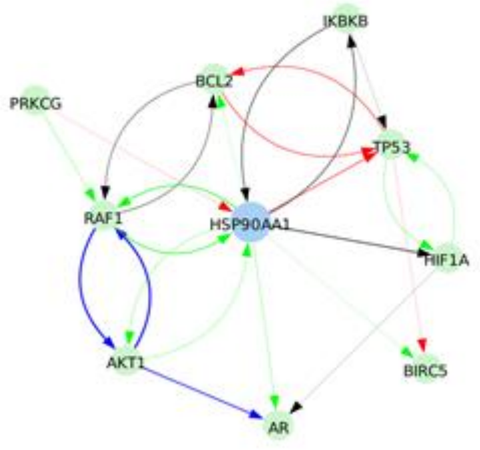


Figure S2: Signed directed interactions between HSP90AA1 and nodes already taken into account in the model.

#### Model extension with the literature

Protein-protein interactions (PPI) and signaling databases are useful to find quickly established interactions between genes and proteins. However, they are not exhaustive and in particular they often lack recent findings. It is therefore necessary to rely on manual literature search to find information on specific prostate cancer components.

The roles of the fusion gene TMPRSS2:ERG and the tumor suppressor NKX3-1 are examples where the information from databases retrieved from Omnipath or PPI databases is lacking, and for which we found additional information from the literature.

Fusion genes are frequently found in human prostate cancer and have been identified as a specific subtype marker (The Cancer Genome Atlas Research Network, 2015). The most frequent is TMPRSS2:ERG. It involves the transcription factor ERG, which leads to cell-cycle progression. ERG fuses with the AR-regulated TMPRSS2 gene promoter to form an oncogenic fusion gene that is especially common in hormone-refractory prostate cancer, conferring androgen responsiveness to ERG. This fusion is not found with Pypath, nor is any target of ERG (Figure S3A). However, literature search reveals that ERG directly regulates EZH2, oncogene c-Myc and tumor suppressor NKX3-1 and many other targets in prostate cancer (Kunderfranco *et al*, 2010).

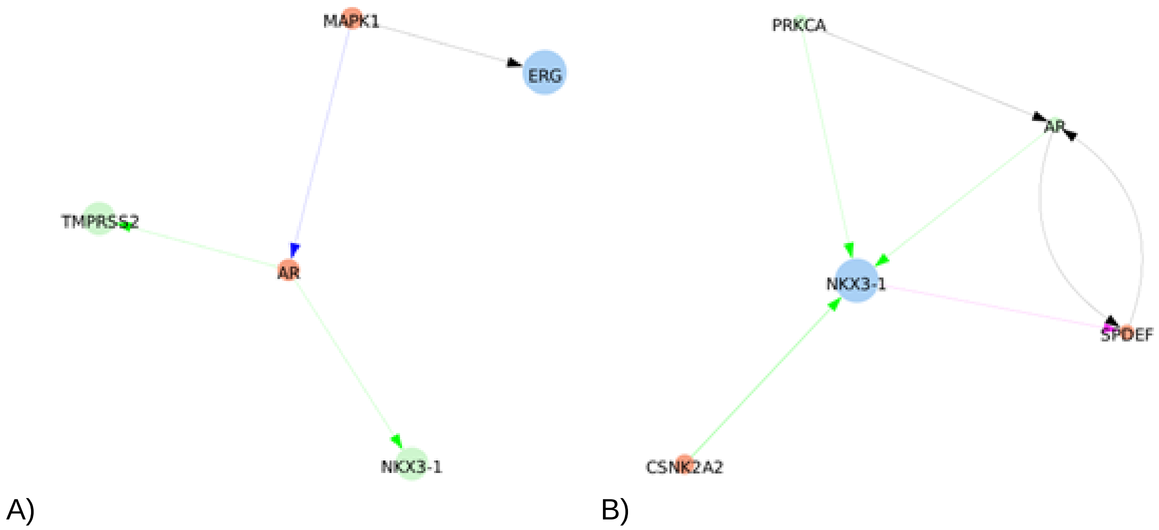


Figure S3: shortest paths found between ERG and TMPRSS2 or NKX3-1 by Pypath: no direct interaction is found.

We model the gene fusion with an activation of ERG by the decoupling of ERG in a special node AR_ERG that is only activated by AR & fused_event node. In the healthy case, fused_event (that represents TMPRSS2) is fixed to 0 or inactive. The occurrence of the gene fusion is represented with the model perturbation where fused_event is fixed to 1. Moreover, ERG expression has a major impact on cell invasion and epithelial-mesenchymal transition (EMT) through the upregulation of the FZD4 gene, a member of the frizzled family of receptors. In our model, we choose for simplicity to consider ERG as a marker of EMT, with a direct activation of the output node EMT by ERG (Adamo & Ladomery, 2016).

NKX3-1 has been identified as a tumor suppressor for prostate cancer. Since it is frequently mutated, it should be included in the model. Some of its regulations can be found with Pypath (Figure S3B), in particular its activation by AR and PKC. However, its role is not identified. The literature search highlighted its role in accelerating the DNA repair response and in particular in avoiding the gene fusion TMPRSS2:ERG. NKX3-1 binds to AR at the ERG gene breakpoint and inhibits both the juxtaposition of the TMPRSS2 and ERG gene loci and also their recombination, by influencing the recruitment of proteins that promote homology-directed DNA repair. Thus, loss of NKX3-1 favors recruitment to the ERG gene breakpoint of proteins that promote error-prone non-homologous end-joining (Bowen *et al*, 2015).

We therefore add the absence of the node NKX3-1 as a new requirement for the activation of ERG by AR and TMPRSS2 in the model. The effect of the gene fusion can be seen in combination with the perturbation that maintains NKX3-1 to the null level.

In contrast with these examples where some knowledge can be retrieved from the literature, some new nodes can not be included in the model in a satisfactory manner, because of missing information about their regulation or role. High-throughput studies have allowed us to identify genes with mutations or expression levels associated with prostate cancer progression or prognosis. But for many of them, the precise mechanisms behind this association remains to be elucidated.

For example, IDH1 (isocitrate dehydrogenase 1) exhibits a recurrent mutation in 1% of primary prostate cancers that defines a specific subtype (The Cancer Genome Atlas Research Network, 2015). This mutant status is associated with a DNA hypermethylation phenotype. Despite a lack of detailed mechanisms linking this gene to the regulation network, we can still reflect a candidate association in the model by including IDH1 as regulated by mTOR and MEK1_2, whose absence (level 0) induces the activation of the output node *Hypermethylation*. The regulation of both new nodes IDH1 and Hypermethylation should be refined when new knowledge is found.

In some cases, we cannot provide any link for a new node, either to an existing node or to a phenotypic output, even qualitatively. For example, ZNF595 has been linked to prostate cancer progression. However, this gene encodes a protein belonging to the Cys2His2 zinc finger protein family, whose members function as transcription factors that can regulate a broad variety of developmental and cellular processes. This knowledge is not detailed enough to add this node in the model yet. However, future mutation data from prostate cancer samples, associated with clinical data, will allow us to test several hypotheses.

This model includes several signaling pathways as well as the substantial cross-talks among them. These pathways range from receptors such as receptor tyrosine kinase (RTKs), androgen receptor (AR) and growth factors pathways (EGF, FGF, TGF-β); downstream gene regulation pathways such as phosphatidylinositol 3-kinase (PI3K)/AKT, Wnt/β-Catenin, NFkB, MAPK, mTOR, SHH, MYC, ETS1, p53, hypoxia-inducible transcription factor (HIF-1) and Smad pathways; cell cycle descriptions with cyclins, E2F1, retinoblastoma protein (Rb) and p21; epithelial-mesenchymal transition (EMT) and migration-related genes; DNA damage and apoptosis-related genes; as well as prostate cancer characteristic genes such as p53, ataxia-telangiectasia mutated (ATM)/ataxia-telangiectasia and Rad3-related (ATR) protein kinases, NKX3.1, TMPRSS2 and TMPRSS2:ERG fusion.

A complete list of the references for all the nodes and edges included in the model can be found in the XLS file of SuppFile 1.

#### Boolean model construction

We converted our network to a Boolean model by defining a regulatory graph, where each node represents a regulatory component and is associated with discrete levels of activity (0, 1 and further integers when justified). Each edge represents a regulatory interaction between the source and target nodes and is labelled with a threshold and a sign (positive or negative). The model is completed by logical rules (or functions), which assign a target value to each node for each regulator level combination. A node that has no regulator is denoted as *input* and a node that doesn't regulate another node is denoted as *output*. *Input* represent different physiological initial conditions and *outputs* represent biological read-outs. The resulting dynamics can be represented in terms of a state transition graph (STG), where the nodes denote the states of the system (i.e. vectors giving the levels of activity of all the variables) and the arcs represent state transitions (i.e. changes in variable values, according to the corresponding logical functions) (for more details, see (Chaouiya *et al*, 2012; Abou-Jaoudé *et al*, 2016)).

When concurrent variable changes are enabled at a given state, the resulting state transition depends on the chosen updating assumptions. Numerous studies use the fully synchronous strategy where all variables are updated through a unique transition. This assumption leads to relatively simple transition graphs and deterministic dynamics. The proportion of initial conditions leading to given attractors is measured as the attractor landscape (Helikar *et al*, 2008; Fumiã & Martins, 2013; Cho *et al*, 2016). However, the synchronous updating assumption approximation often leads to spurious cyclic attractors. On the other hand, the fully asynchronous updating assumption considers separately all possible transitions and therefore allows the consideration of alternative dynamics in the absence of kinetic data. The resulting dynamics has a branching structure which makes it more difficult to evaluate. In this project, we consider asynchronous dynamics mixed with stochastic simulations. The regulatory graph was constructed using GINsim software (Chaouiya *et al*, 2012) and then exported in a format readable by MaBoSS software (see below) in order to perform stochastic simulations on the Boolean model.

The final model accounts for 133 nodes and 449 edges (Figure 1 and SuppFile 1) and includes pathways such as androgen receptor and growth factor signalling, several signalling pathways (Wnt, NFkB, PI3K/AKT, MAPK, mTOR, SHH), cell cycle, epithelial-mesenchymal transition (EMT), Apoptosis, DNA damage, etc. This model has 9 inputs (*EGF, FGF, TGF beta, Nutrients, Hypoxia, Acidosis, Androgen, TNF alpha* and *Carcinogen* presence) and 6 outputs (*Proliferation*, *Apoptosis, Invasion, Migration,* (bone) *Metastasis* and *DNA repair*).

### Boolean model simulation

#### Wild type simulation

Our prostate Boolean model recapitulates known phenotypes of prostate cells by stochastic simulations in each of the studied “physiological” conditions. The model can be considered as a model of healthy prostate cells when no mutants or fused genes are present, called wild-type model in present work. These healthy cells mostly exhibit quiescence in absence of any input. Because the initial conditions of all components of the model are set to random values and input nodes are OFF, there is a possibility to transiently activate the pathways but not to maintain them, and all pathways are eventually turned off.

Our prostate Boolean model was simulated using MaBoSS and asynchronous updates and recapitulates known phenotypes of prostate cells under physiological conditions (Main text, Figure 2). Model states distribution at the end of the simulation with growth factors, *Nutrients* and *Androgen* as inputs can be seen in Figure 2B. Note that some outputs are not mutually exclusive, therefore the presence of cells with *Invasion* and *Proliferation*. In Figure 2C, the same model with cell death factors ON.

In proliferating conditions, transient probabilities of the cyclins can be used to check that the order of activations of these nodes in the paths leading to the cyclic attractor is consistent with a proper cell cycle progression (Figure S4).

These analyses can be performed using model files from SuppFile 1 and the jupyter notebook from SuppFile 2.

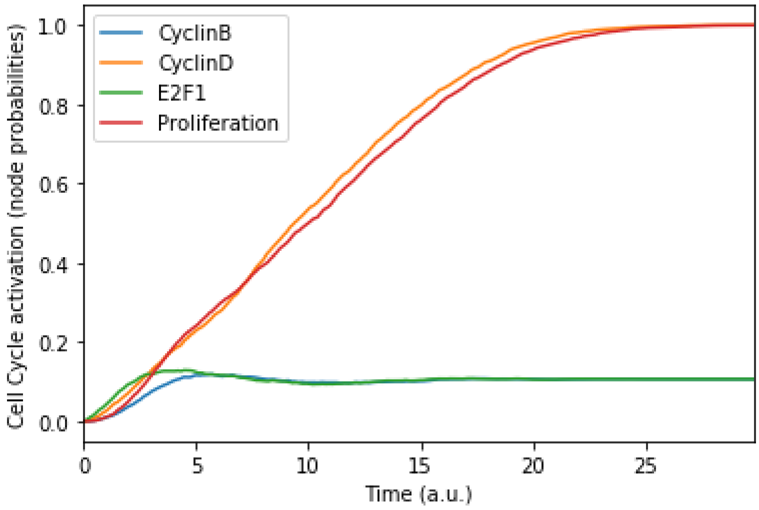


Figure S4: Mean probabilities of the nodes characterising the cyclins and proliferation, with nutrients and growth factors as inputs. We choose initial states for the nodes involved in the cell cycle that correspond to quiescence (cyclins OFF, cell cycle inhibitors Rb and p21 ON), in order to visualise the order of activation of the cyclins: first Cyclin D, then Cyclin B. The mean probabilities reach asymptotic levels because of the desynchronisation of stochastic trajectories in the population.

### Mutants simulation

A mutant in the logical framework is simulated by setting the node corresponding to the gene mutated to 0 in the case of loss of function and to 1 in the case of gain of function. The effect of a mutation is assessed, like the change of initial conditions, by comparing the mutant's probabilities of reaching a phenotype with respect to the wild-type model. Therefore, mutations change the model phenotypes: *Apoptosis, Proliferation, Invasion, Migration,* (bone) *Metastasis* and *DNA repair.*

#### Single mutations

The single mutations of some of the main nodes of the network show some changes in the probabilities of reaching the phenotypes when compared to wild type conditions.

The examples on Figure S5 show that a loss-of-function mutation of FOXA1 in proliferative conditions (nutrients and growth factors) results in the activation of migration and invasion but not metastasis. A loss-of-function mutation of TP53 in the same condition with the addition of carcinogen does not lead to loss of the apoptosis induced by DNA damage because of the activation of caspase 3 pathway.

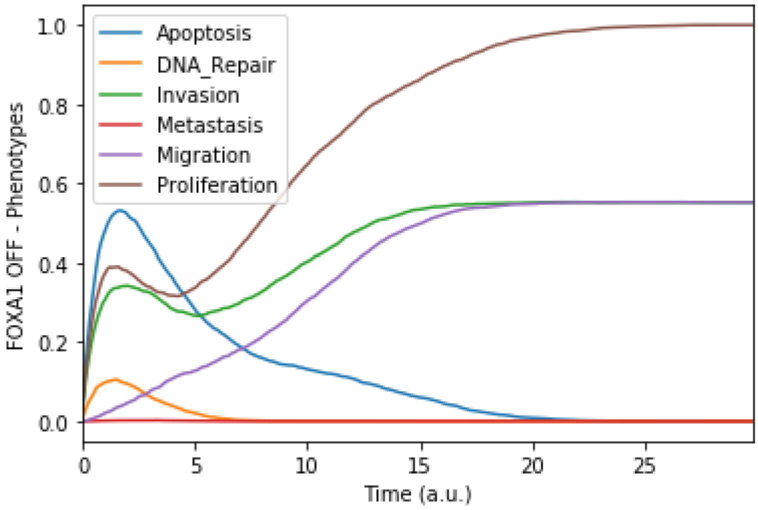


A)

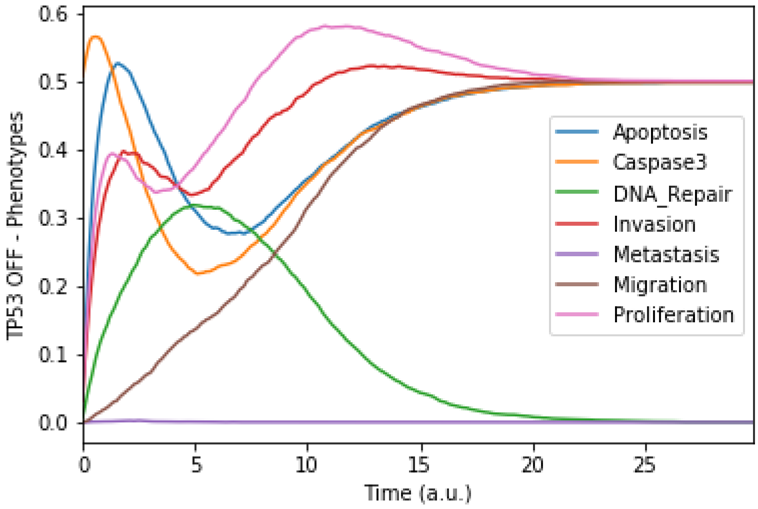


B)

Figure S5: Mean probabilities in simulations of mutated models. A) Loss-of-function mutation of FOXA1. B) Loss-of-function mutation of TP53.

#### Multiple mutations

Cancer progression is characterized by the accumulation of genetic alterations that affect multiple pathways in the signaling network. The logical model allows to easily simulate all possible combinations of mutations and study the potential redundancy or synergy of alteration effects and the importance of order. An example of double mutation is shown in Figure S6, where the combination of the gene fusion AR:ERG and the loss-of-function of NKX3-1 activates bone metastasis signals in proliferative conditions with androgen induction.

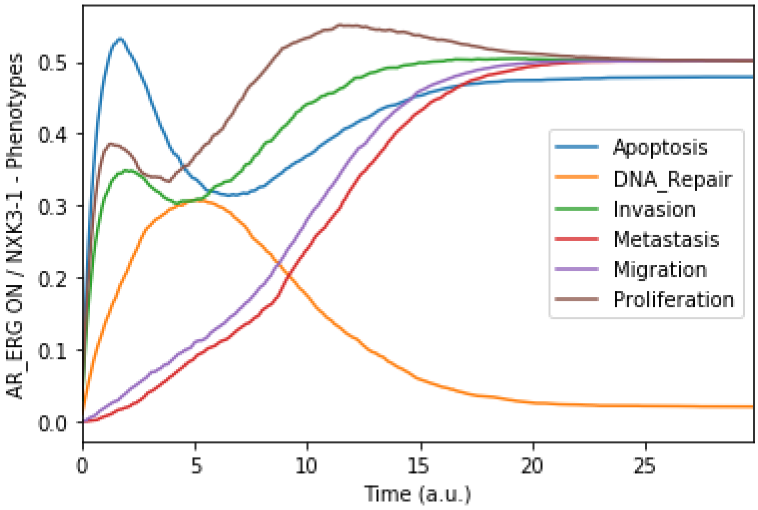


Figure S6: Mean probabilities in simulations of the model with a multiple simulation: the gene fusion TMPRSS2:and a loss-of-function of NKX3-1.

The model allows to study easily all possible associations of mutations to assess synergies or redundancies. It can also reproduce sets of mutations observed in tumors. Different sequences of possible acquired mutations can be simulated and compared to what is already known about patients harboring these mutations.

### Personalised Boolean models of TCGA patients

Our prostate Boolean model was tailored to a set of 488 TCGA prostate cancer patients using our PROFILE personalisation method (Béal *et al*, 2019). The distribution of the 488 patients' Gleason score can be seen in Figure S7. The prostate cancer patients recipe that has a better correlation with their Gleason score was using mutations and copy number alterations (CNA) as node activity status and RNA as initial conditions and transition rates (Figure S8-S12). All the 488 TCGA prostate cancer patients models can be found in MaBoSS format in SuppFile 3, TCGA-specific personalised models.

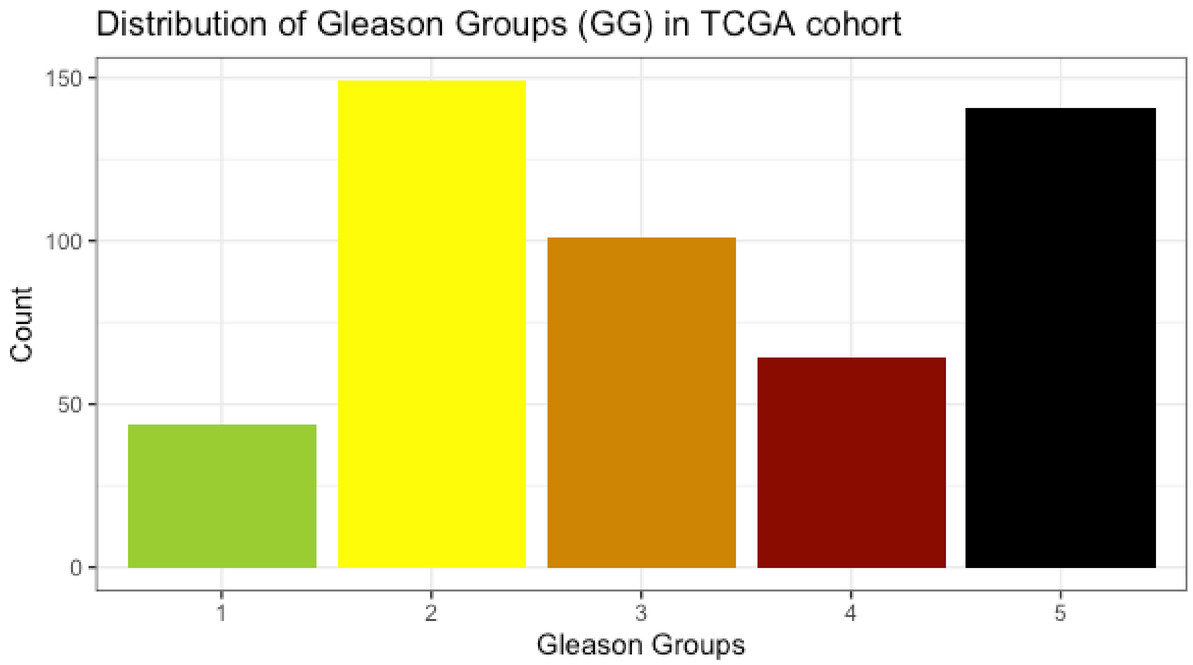


Figure S7: Distribution of 488 TCGA prostate cancer patients' samples per Gleason group.

#### Phenotype distribution of TCGA patients

One of the quality checks performed in PROFILE is to build models using different recipes, i.e., using different data to modify different model variables, and to compare them to some clinical grouping or expression signature to rank them and select the most performing one. In our case, we used five different recipes (only mutations, mutations and CNA, mutations and RNA data, mutations, CNA and RNA data and only RNA data), we grouped the patients by their GG (either 3- or 5-stage) and studied the distributions of the different phenotypes scores: *Apoptosis* (Figure S8), *DNA repair* (Figure S9), *Invasion* (Figure S10), *Migration* (Figure S11) and *Proliferation* (Figure S12). Finally, we chose the recipe that uses mutations, CNA and RNA data as it included the most quantity of data and reproduced desired results (SuppFile 3, TCGA-specific personalised models).

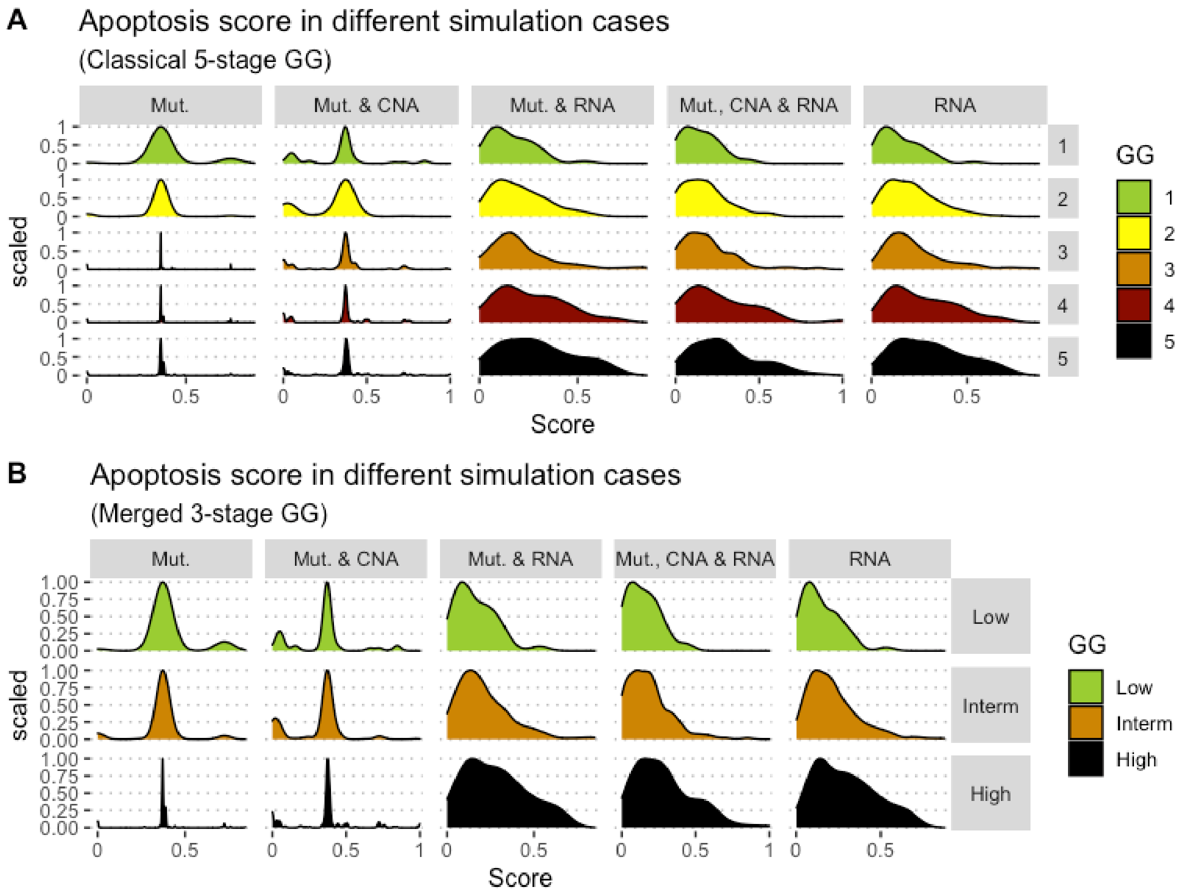


Figure S8: Associations between simulations and Gleason groups (GG). Distribution histograms of *Apoptosis* scores according to GG in 3 groups (A) and 5 groups (B). Columns correspond to different personalisation recipes (see (Béal *et al*, 2019) for more details).

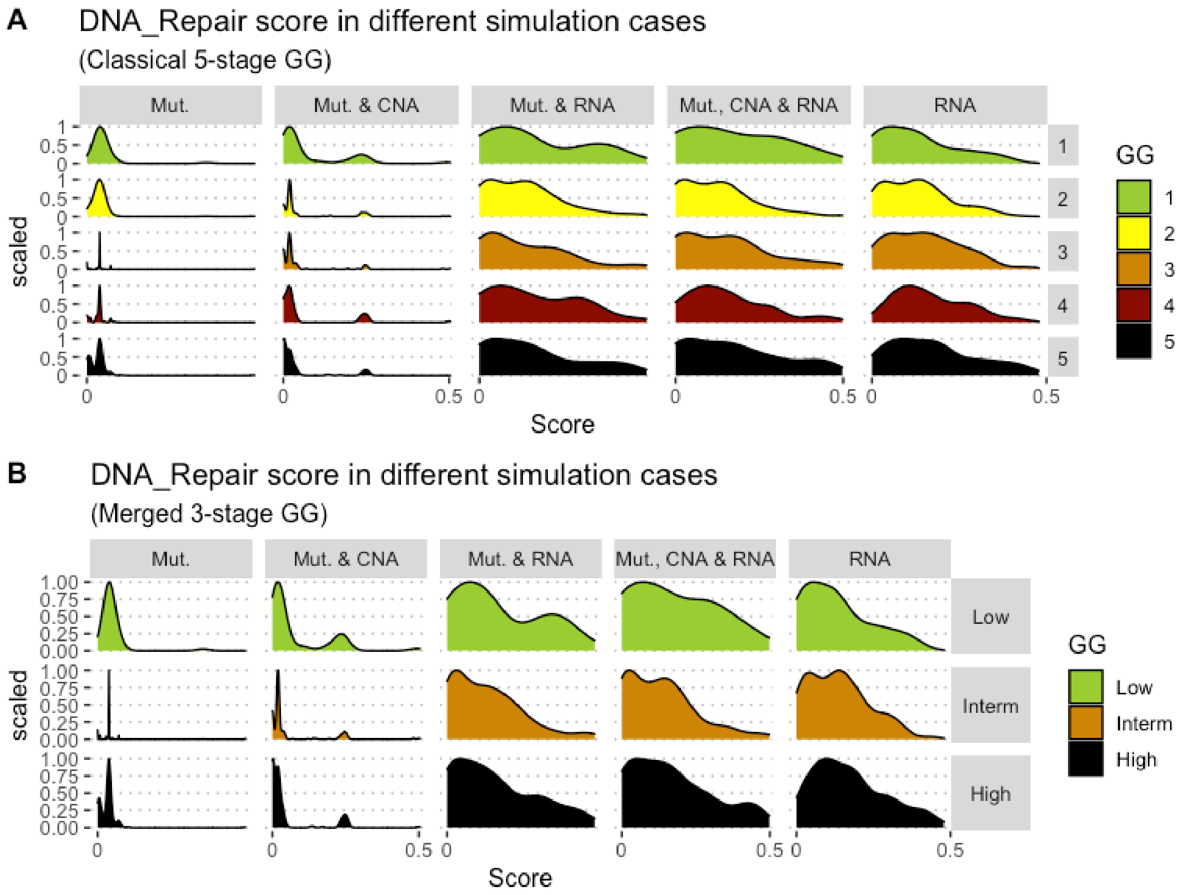


Figure S9: Associations between simulations and Gleason groups (GG). Distribution histograms of *DNA_repair* scores according to GG in 3 groups (A) and 5 groups (B). Columns correspond to different personalisation recipes (see (Béal *et al*, 2019) for more details).

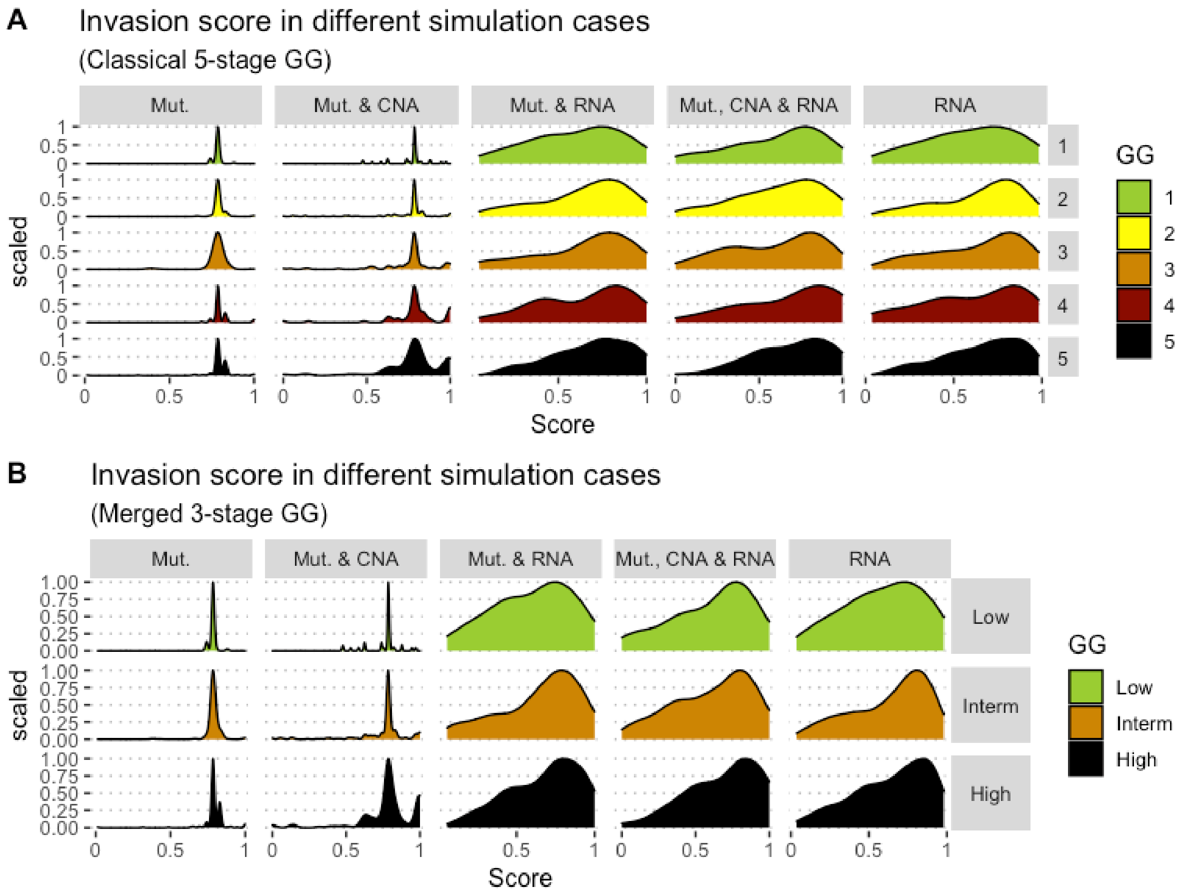


Figure S10: Associations between simulations and Gleason groups (GG). Distribution histograms of *Invasion* scores according to GG in 3 groups (A) and 5 groups (B). Columns correspond to different personalisation recipes (see (Béal *et al*, 2019) for more details).

**A**

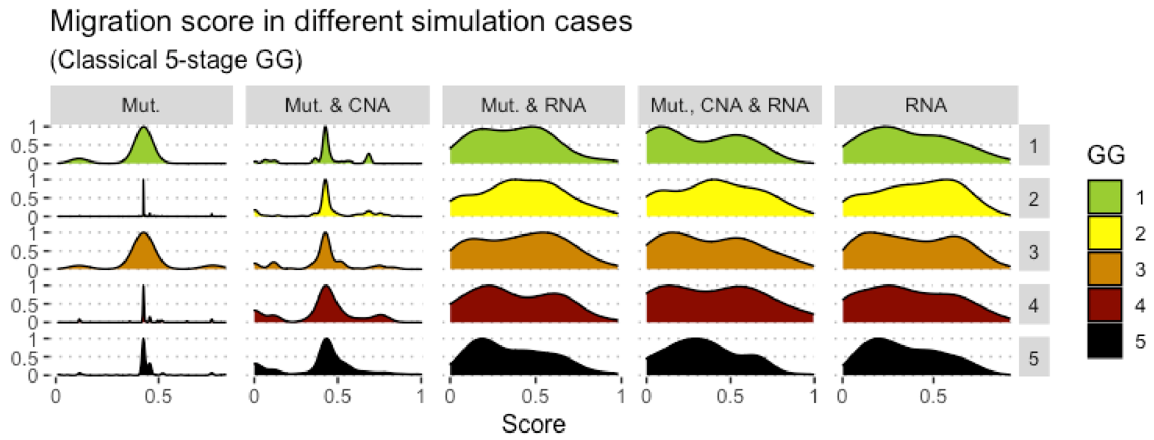


**B**

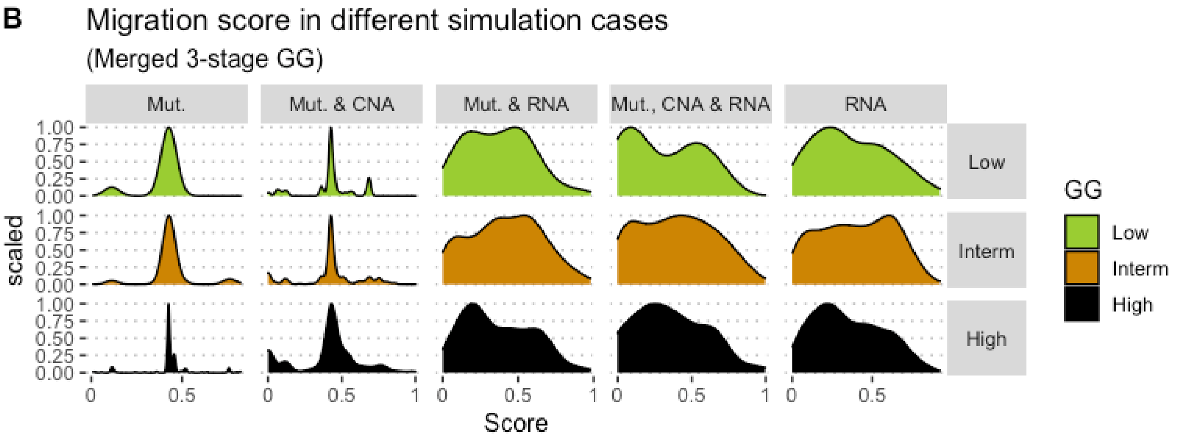


Figure S11: Associations between simulations and Gleason groups (GG). Distribution histograms of *Migration* scores according to GG in 3 groups (A) and 5 groups (B). Columns correspond to different personalisation recipes (see (Béal *et al*, 2019) for more details).

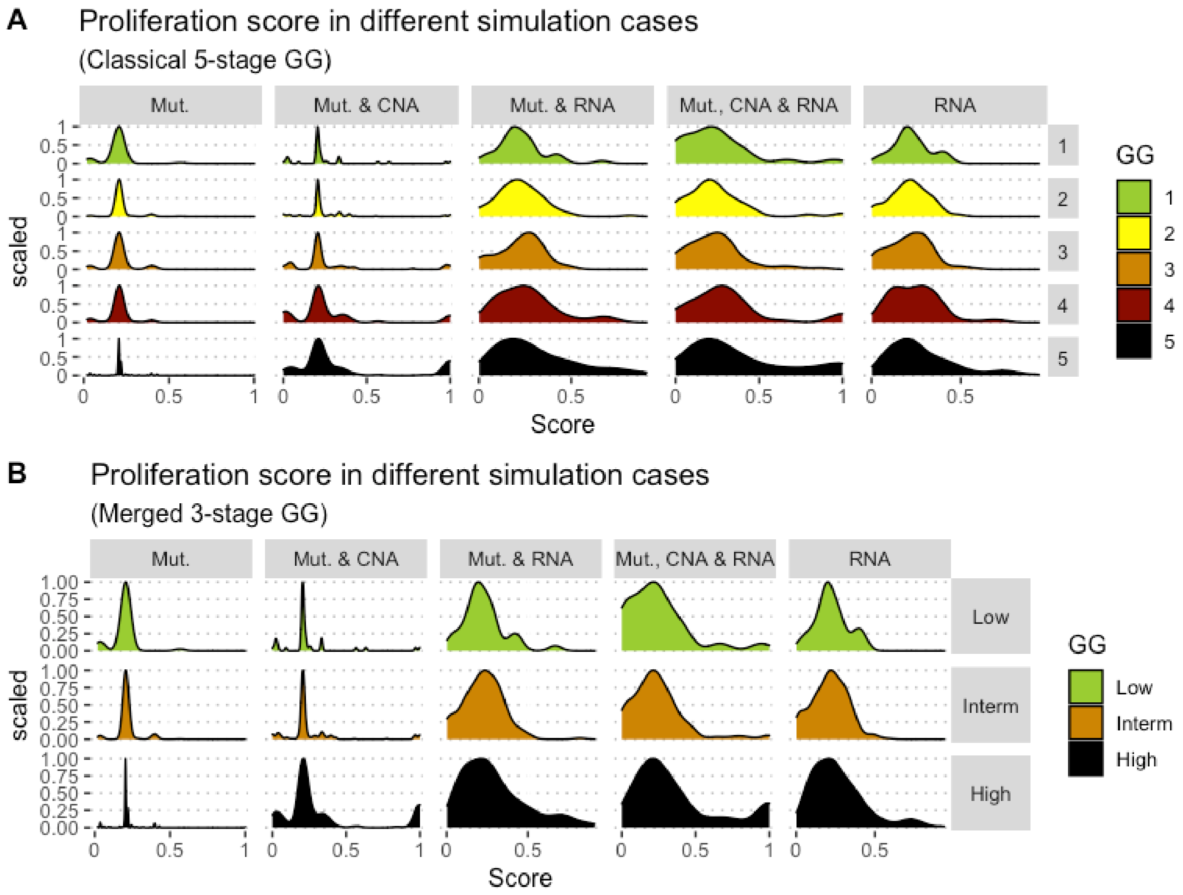


Figure S12: Associations between simulations and Gleason groups (GG). Distribution histograms of *Proliferation* scores according to GG in 3 groups (A) and 5 groups (B). Columns correspond to different personalisation recipes (see (Béal *et al*, 2019) for more details).

Next, we took the personalised models that used mutations, CNA and RNA data and performed a PCA analysis on the 488 TCGA patients (SuppFile 3) and their 5 phenotype scores that result from simulating them using MaBoSS. For these PCA, we grouped the patients by 3-stages GG (Figure S13) and 5-stages GG (Figure S14). In addition and for the sake of clarity, we reduced each of these groups to their barycenter (Figure S15 for 3-stages GG and Figure S16 for 5-stages GG), where we can see that higher GG move towards Proliferation, Invasion and Migration variables.

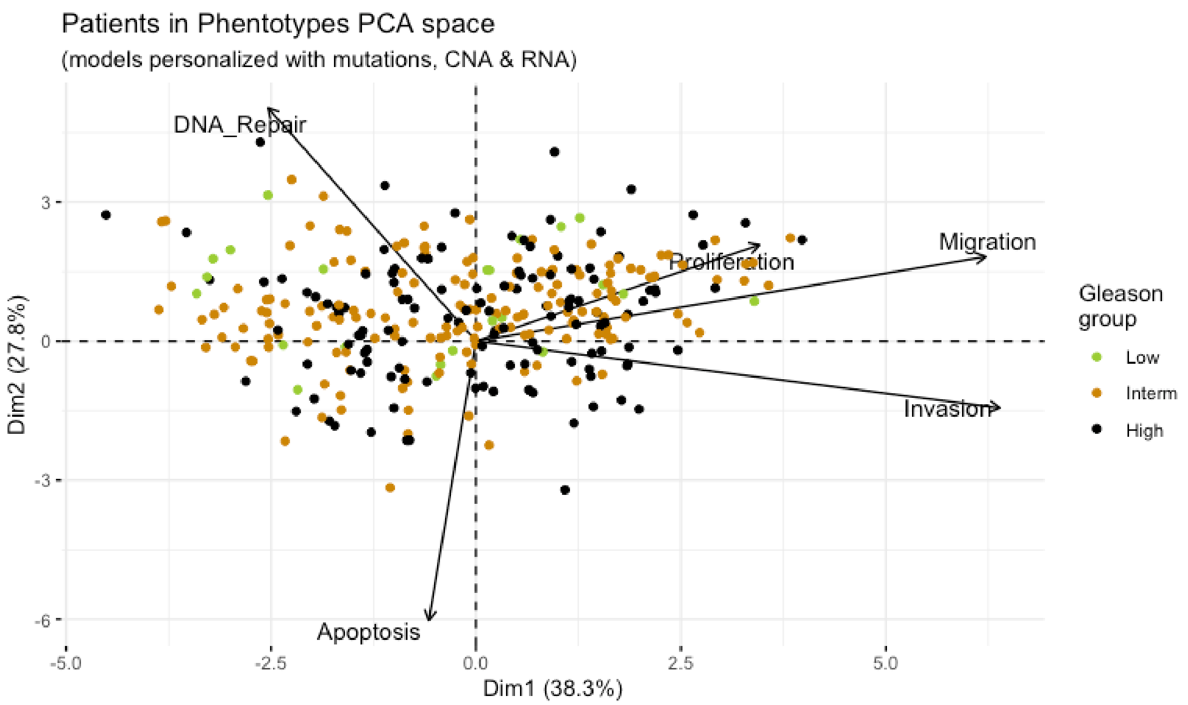


Figure S13: Principal Component Analysis of all 488 TCGA patients in 3 Gleason Groups using the vectors of all 5 phenotypes from the model.

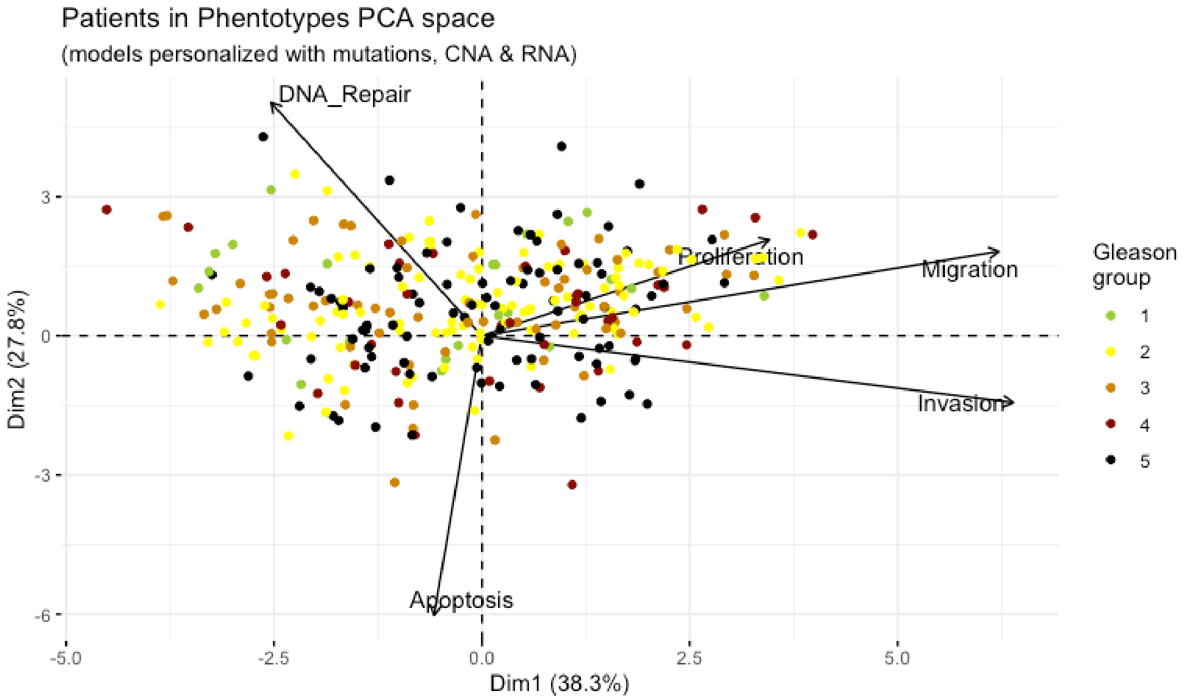


Figure S14: Principal Component Analysis of all 488 TCGA patients in 5 Gleason Groups using the vectors of all 5 phenotypes from the model.

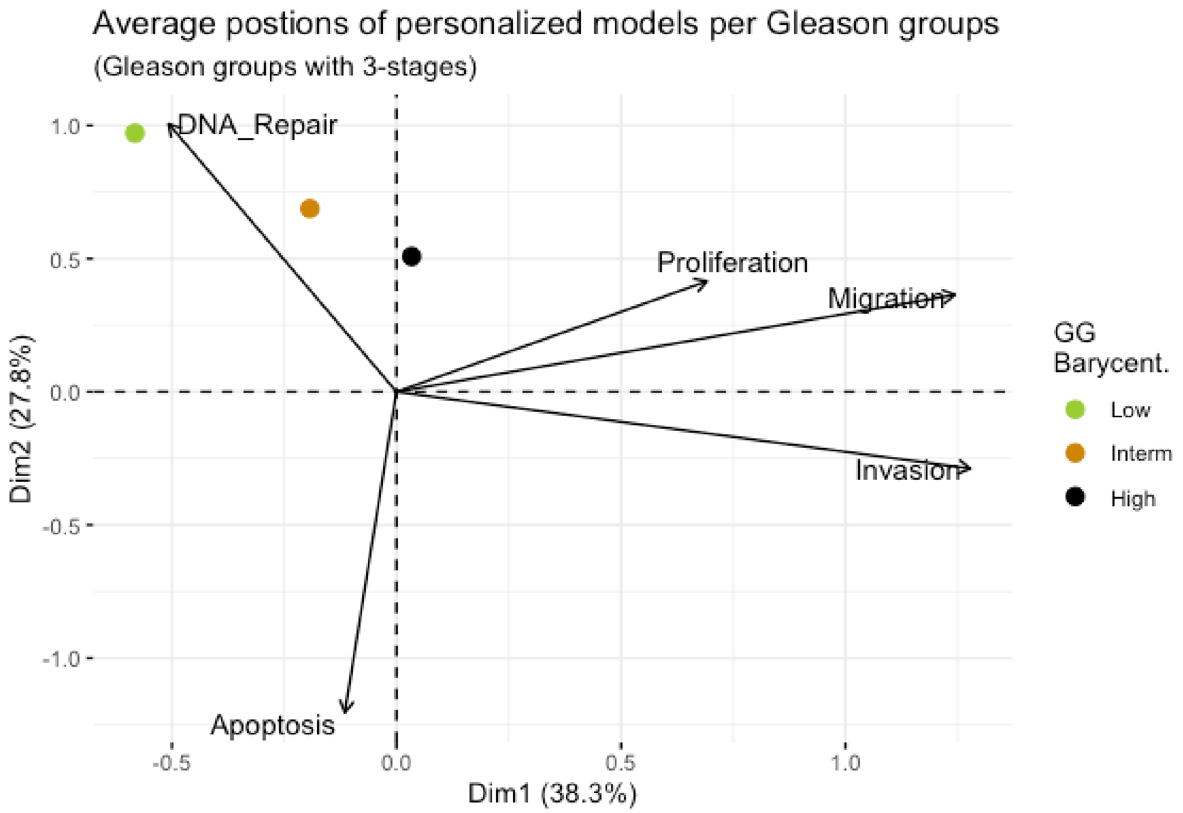


Figure S15: Principal Component Analysis' barycenters of all 488 TCGA patients grouped in 3 Gleason Groups using the vectors of all 5 phenotypes from the model. This is the same figure as Figure 3 in the main text.

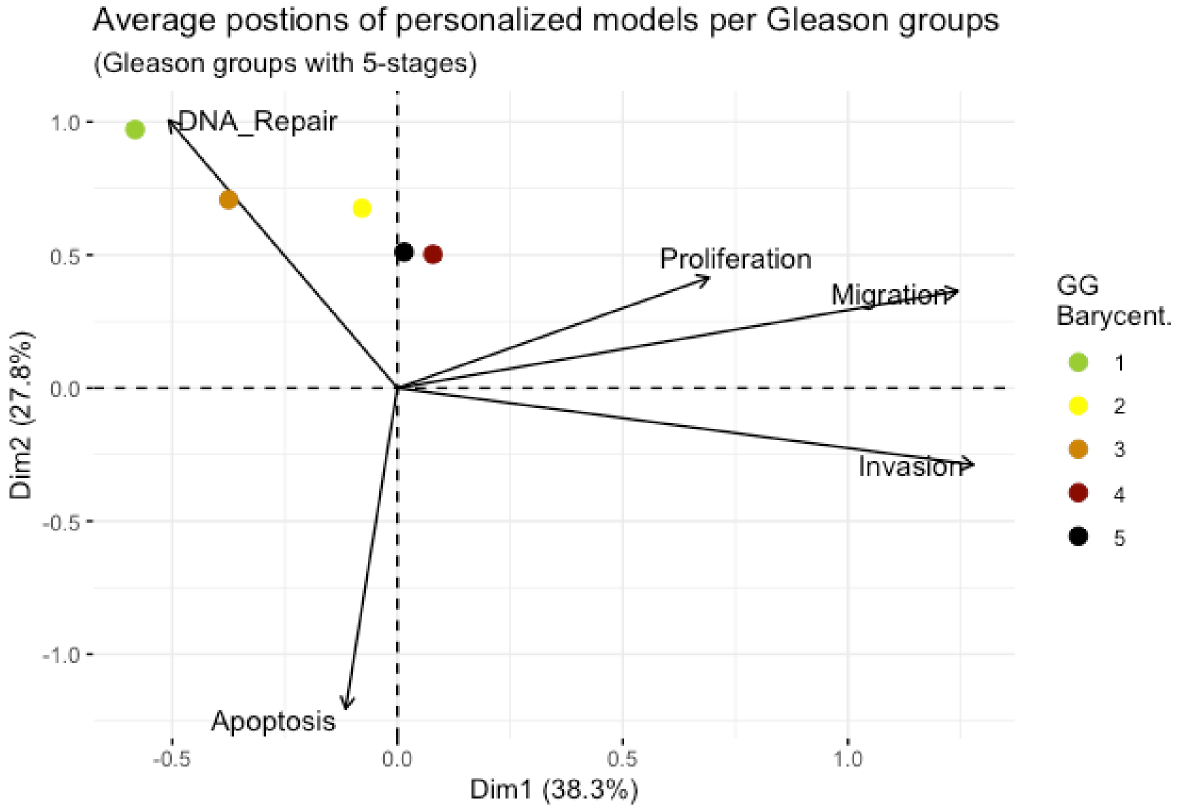

Figure S16: Principal Component Analysis' barycenters of all 488 TCGA patients grouped in 5 Gleason Groups using the vectors of all 5 phenotypes from the model.

### Analysis of drugs that inhibit the activity of genes of TCGA patients

Using our pipeline of tools (Montagud *et al*, 2017), we performed the analysis of all single perturbations that reduce *Proliferation* or increase *Apoptosis* together with the combined perturbations of a set of selected genes that are targets of already-developed drugs relevant in cancer progression (Table 1). Then, we aggregated the results of the 488 patients to identify which inhibitions affected *Proliferation* (Figure S17) and *Apoptosis* (Figure S18) the most in this cohort.

Interestingly, we found several genes that were found as suitable points of intervention in most of the patients (MYC_MAX complex and SPOP were identified in more than 80% of the cases) (Figure S17 and S18), but others were specific to only some of the patients (MXI1 was identified in only 4 patients, 1% of the total, GLI in only 7% and WNT in 8% of patients).

The inactivation of some of the targeted genes had greater effect in some patients than in others, suggesting the possibility for the design of personalised drug treatments (Main text). Nevertheless, knowing that some treatments that inhibit one gene are already able to reduce Proliferation phenotypes considerably, we explored the possibility of finding combinations of treatments that could lead to the same types of outcomes. One reason for searching for coupled drugs is that these combinations allow the use of lower doses of each of the two drugs and thus reduce their toxicity. It is important to note, though, that the analyses performed with the mathematical model do not aim at predicting drug dosages per se but to help in the identification of potential candidates.

The exhaustive search for combinations of drugs for each patient of the cohort requires an extensive amount of computation time (9 days and 7 hours on a personal computer or 3 hours on 20 nodes with 48 CPUs each, per model) as all variables of the model are automatically overexpressed and inhibited, one by one and in pairs, leading to a vast amount of simulations. For this reason, we have narrowed the list of potential candidates to reduce *Proliferation* or increase *Apoptosis* by performing the analysis of all single perturbation and selecting the combined perturbations of a set of selected genes that are targets of already-developed drugs relevant in cancer progression (Main text, Table 1).

We used the models to grade the effects that the combined treatments would have in each one of the 488 TCGA-patient-specific models. The resulting list of combinations vary greatly from patient to patient, making it infeasible economically for labs and companies to pursue true patient-specific treatments. It also poses challenges in clinical trial designs aimed at validating the model based on the selection of treatments. Because of these constraints, it is more interesting commercially to target group-specific treatments, which can be more easily related to clinical stages of the disease. Mathematical modelling of patient profiles would then help to classify them in these groups, providing, in essence, a grade-specific treatment.

The TCGA mutants and their normalised phenotype scores in regards to the WT model can be found in SuppFile 4.

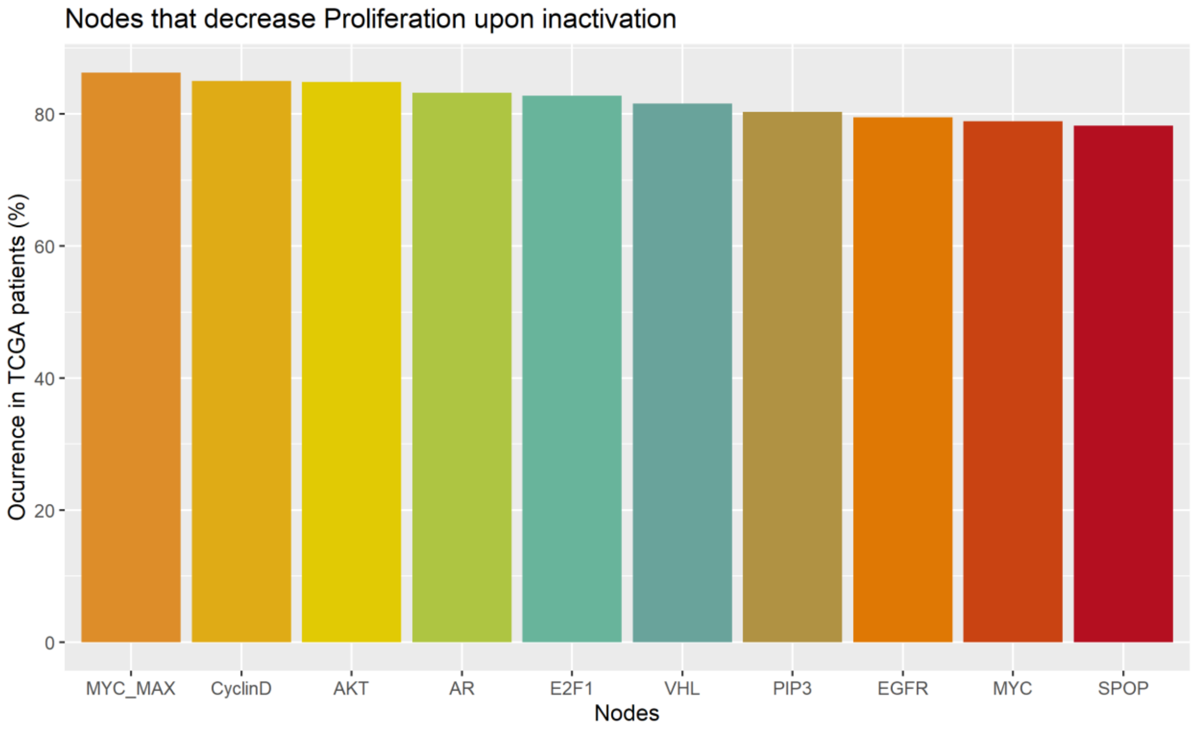


Figure S17: Nodes in the Boolean model that hamper *Proliferation* upon inactivation aggregating all patient-specific results.

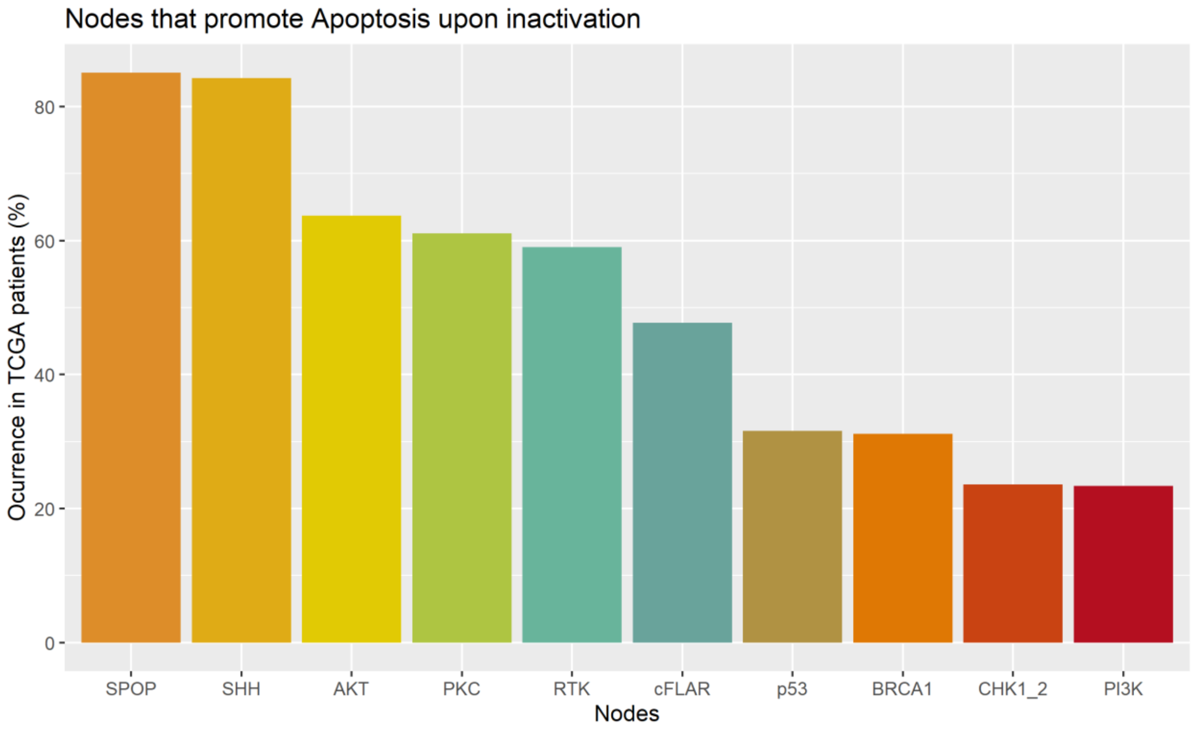


Figure S18: Nodes in the Boolean model that promote *Apoptosis* upon inactivation aggregating all patient-specific results.

### Personalised Boolean models of prostate cell lines

We tailored our generic prostate model to 8 prostate-specific cell lines: 22RV1 (Sramkoski *et al*, 1999), BPH-1 (Hayward *et al*, 1995), DU-145 (Stone *et al*, 1978), LNCaP-Clone-FGC (Horoszewicz *et al*, 1983), NCI-H660 (Johnson *et al*, 1989; Lai *et al*, 1995; Castoria *et al*, 2011), PC-3 (Kaighn *et al*, 1979), PWR-1E (Webber *et al*, 1996) and VCaP (Korenchuk *et al*, 2001). These cell lines had available datasets in the GDSC resource (Iorio *et al*, 2016) and these were used to personalize models using our PROFILE framework (Béal *et al*, 2019) and using mutation data as discrete data and RNA as continuous data (Figure S19).

We simulated the prostate-cell-line-specific models under random initial conditions and observed that they generated distinctive phenotype probabilities and captured some of the differences described in literature (Figure S19). For instance, it has been described that PC-3 cell line has high migratory potential compared to DU-145 cells, which have a moderate migratory potential, and to LNCaP cells, which have low migratory potential (Cunningham & You, 2015). In our simulations, we capture that PC-3 has greater invasiveness, migration and proliferation than DU-145. However, the invasiveness and proliferation potential of LNCaP found is much higher than PC3. Note that these results come from a collection of datasets from GDSC and a Boolean model that includes a subset of the interactions of 312 proteins. Distortions from real-life behaviour are expected and will be the focus of further research, such as the high LNCaP invasiveness or the lack of difference of the benign cell lines (BPH-1 and PWR-1E) with the rest of the cell lines.

As we did for the TCGA patients' study, we tried different personalisation recipes to personalise these cell lines, but as they had no associated clinical features, we were left with the comparison of the different values for the model's outputs among the recipes. We chose the aforementioned recipe as it included two different data types (RNAseq and mutations) and reproduced desired results (Figure S19). Nevertheless, we could have considered using mutation and CNA as discrete data and RNA as continuous data, but the inclusion of CNA data forced LNCaP proliferation to be zero (Figure S20). This is due to the fact that CNA data used as discrete data forces several nodes to be active or inactive throughout the simulation, as if they were mutants. Notably, CNA data forces E2F1 node to be 0, which forces Cyclin B to be 0 and it forces SMAD node to be 1, which forces MYC_MAX node to be 0 and p21 node to be 1, forcing Cyclin D to be 0. Without either Cyclin B or D, the model cannot activate the *Proliferation* node.

In addition, we wanted to study these different personalisation recipes to try to better match simulated phenotypes and cell-line phenotypes described experimentally, but we had similar results (Figures S20). Furthermore and due to the mismatches of cell line models with their described biology characteristics, we went back to the source data to study if these mismatches were something we could correct on the model or a problem of the dataset we used to personalise the model. We performed principal component analysis (PCA, using FactoMineR R package (Lé *et al*, 2008)) (Figure S21) on the dataset used to personalise the models: an RNAseq dataset of 111 genes. We found that the cell lines do not cluster by their characteristics: DU-145, an invasive cell line, is close to BPH-1 and PWR-1E, non-invasive cell lines.

Furthermore, we digged into the pathways that are characteristic of each of these cell lines by using single sample GSEA (using ssGSEA 2.0 R package (Krug *et al*, 2019)) (Figure S22) on the same RNA dataset using the Hallmarks molecular signatures. We found that out of the 50 Hallmarks, 21 have an overlap of more than 5 genes with the model's genes. Thus, we set to cluster the cell lines by using the signatures of each one of them in these 21 pathways. The results are quite telling of the lack of clear clustering of these cell lines with their different characteristics (Figures S21 and S22): invasive and non-invasive cell lines have similar signature values in EMT or G2M checkpoint pathways, BPH-1 clusters with NCI-H660 and PWR-1E with DU-145, etc.

All in all, it is unrealistic to expect that a model of different cellular behaviours will match all biological aspects and characteristics as models are, by definition, abstractions of reality (Rosenblueth & Wiener, 1945; Korzybski, 1995). For instance, if one were to match the cell lines' doubling times, of which *Proliferation* phenotype should be a good proxy (St John *et al*, 2012; Cunningham & You, 2015), such a study would need a deeper understanding of the cell's biology, the modelling of many more processes, with many more parameters, and a more complete simulation framework both multi-scaled and finer-grained, which is beyond the scope of the present work.

All these cell-line-specific personalised models are publicly available in SuppFile 5.

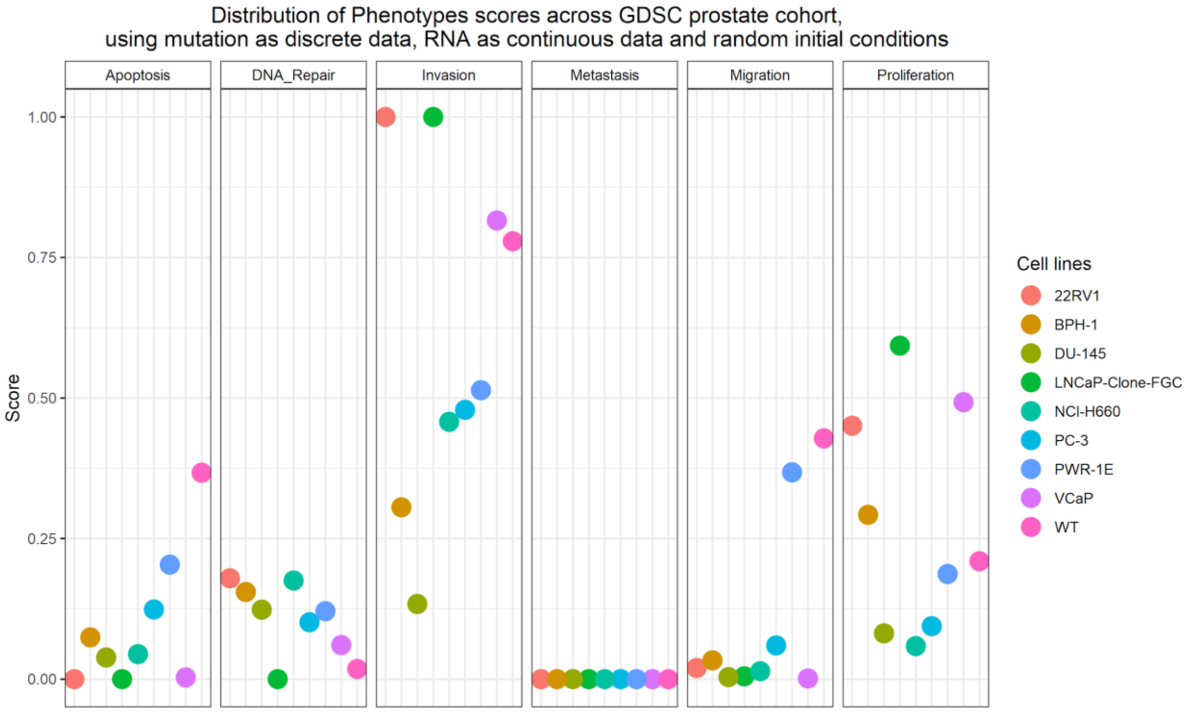


Figure S19: Phenotype simulation results across GDSC prostate-cell-line-specific Boolean models' simulation with random initial conditions. WT stands for wild type model, the original prostate model with no personalisation.

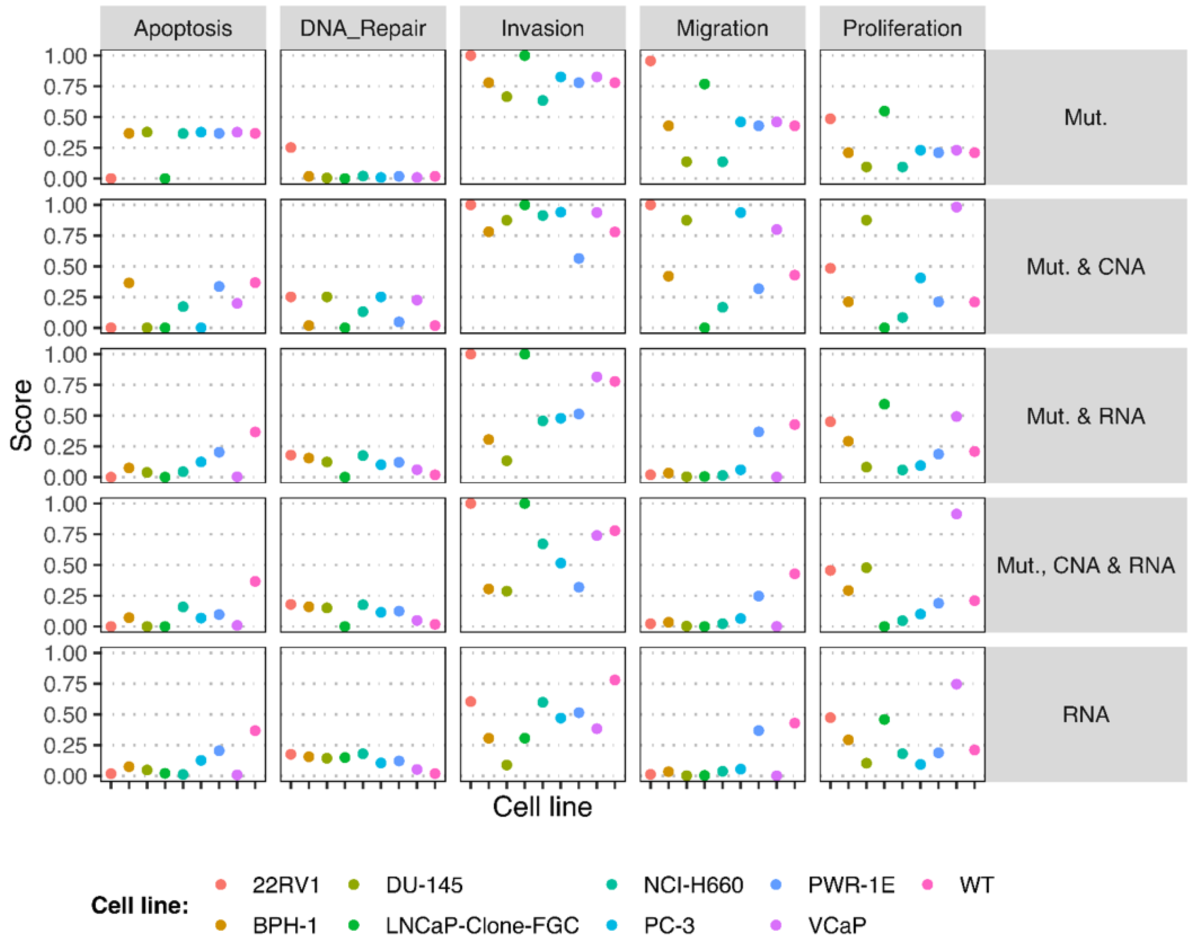


Figure S20: Phenotype simulation results across GDSC prostate-cell-line-specific Boolean models' simulation with random initial conditions under different personalisation recipes. Mutatiosn and CNA are always considered as discrete data and RNA expression is always considered as continuous data.

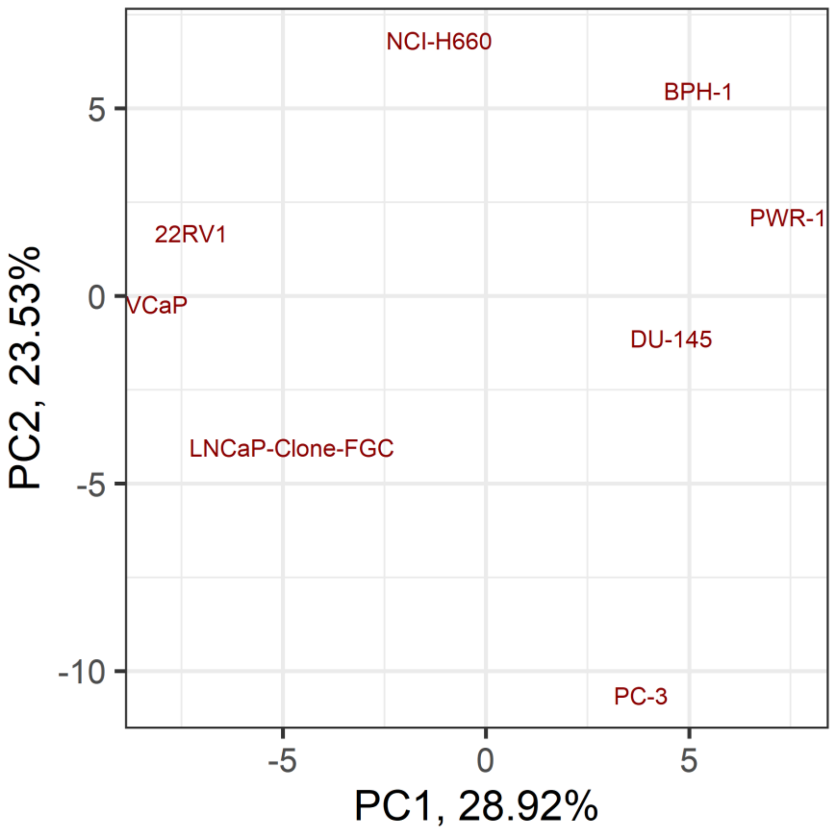


Figure S21: Principal Component Analysis (PCA) of the RNA dataset used to tailor the prostate cell lines.

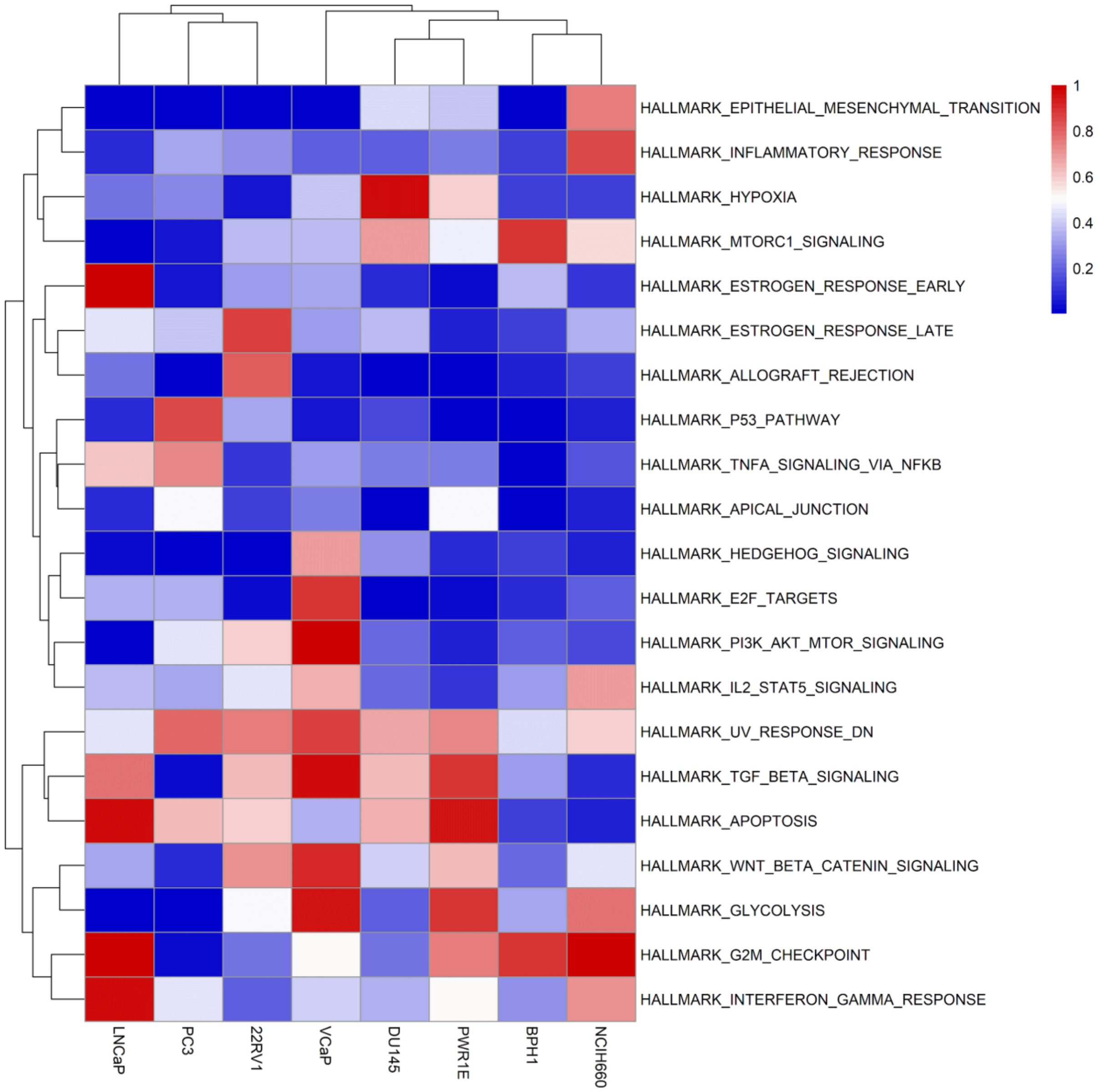


Figure S22: Results of the ssGSEA performed on the RNA dataset used to tailor the prostate cell lines.

### Personalised LNCaP Boolean model

LNCaP model was selected to study its genetic interactions and its uses for drug discovery. The simulation of the LNCaP-specific model under random initial conditions leads to 4 most probable phenotypes: Invasion-Migration, Invasion-Migration-Proliferation, Invasion-Proliferation and Invasion. Using MaBoSS software, we were able to assign probabilities to each one of these phenotypes (Figure S23 and SuppFile 1 and 2).

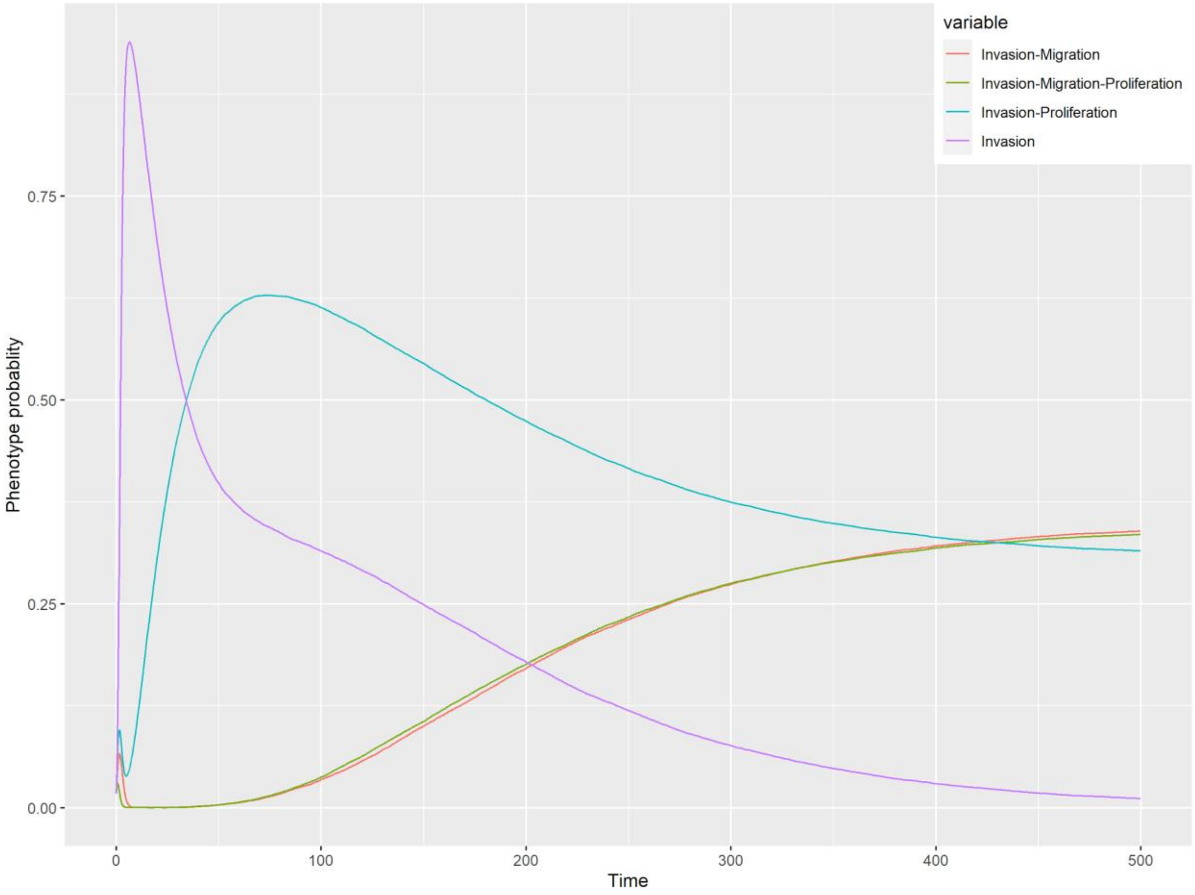


Figure S23: Phenotype probabilities of LNCaP model under random initial conditions.

Additionally, we studied the LNCaP model under four different growth conditions that could be reproduced in experiments. These are a nutrient-rich media that mimics the RPMI supplemented with glucose and fetal bovine serum with additional androgen, EGF, both or none (Figure S24).

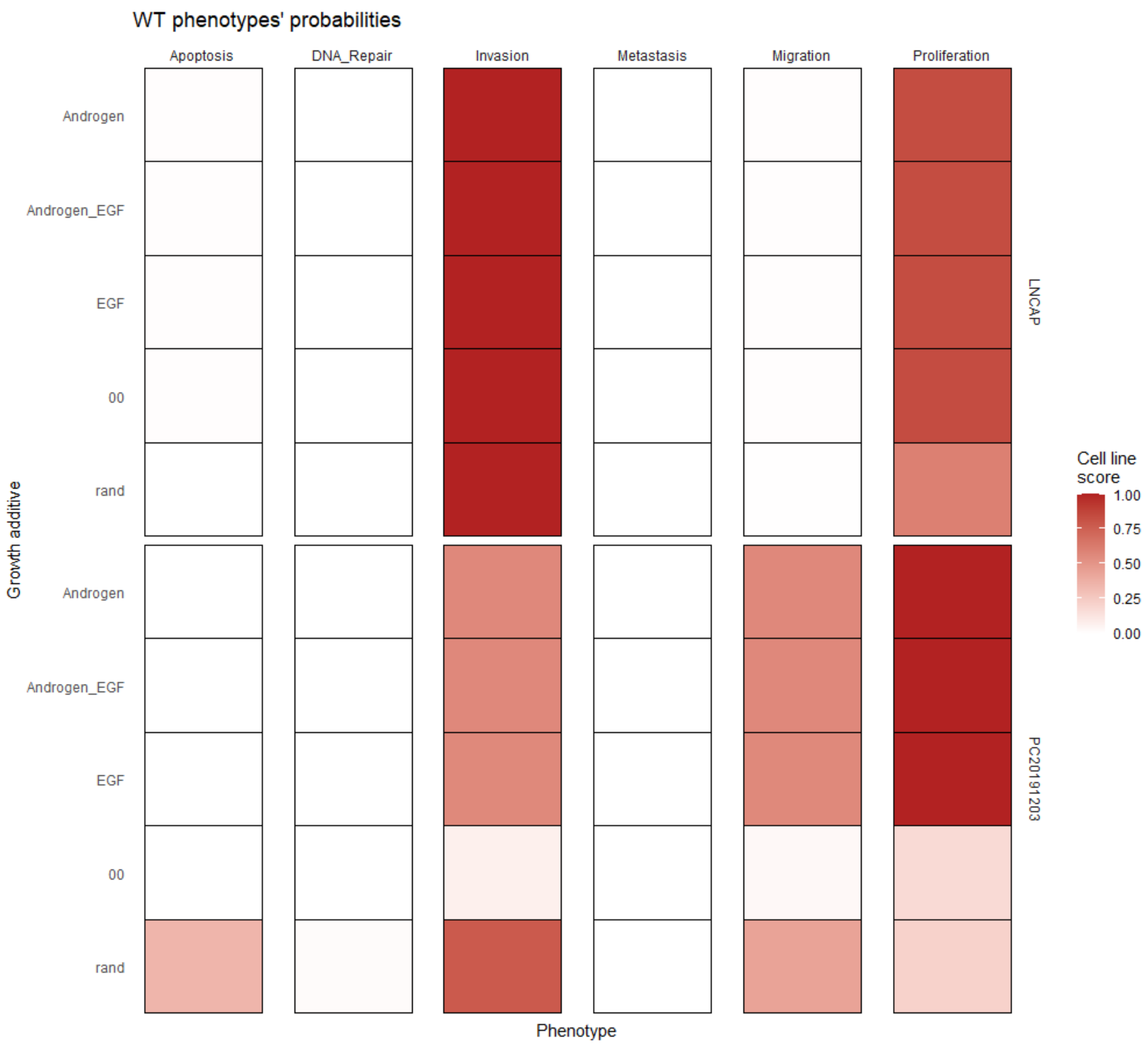


Figure S24: WT and LNCaP-specific model phenotype probability variations under four different growth conditions. AR stands for androgen presence, EGF for EGF presence, and 00 for lack of any of both.

### High-throughput mutant analysis of LNCaP model

A mutant in the logical framework is simulated by setting the node corresponding to the gene mutated to 0 in the case of loss of function and to 1 in the case of gain of function. The effect of a mutation is assessed, likewise to the change of initial conditions, by comparing the mutant's probability of reaching a phenotype with respect to the wild type model. Therefore, mutations change the model phenotypes' probabilities and this can be compared to the wild type model.

The logical model allows to easily simulate all possible combinations of mutations and study the potential redundancy or synergy of these perturbations. To perform this, tools like our high-throughput mutant analysis pipeline (Montagud *et al*, 2017) are ideally suited. This pipeline of tools was applied to the LNCaP-specific model in order to study all single and double mutants of LNCaP model (32258 mutants) and their probabilities of reaching all the phenotypes of the model.

The double mutants of the high-throughput mutant analysis were used to identify genetic interaction relationships, such as epistasis, among the single mutants. Phenotype probabilities' variations of all 32258 models were compared to the wild type model and were used to identify relevant combinations of perturbations that affect phenotypes of interest (Figure S25) and single phenotypes (Figure S26). In these figures, a PCA was applied to the matrix of the 7 phenotype probabilities of the 32258 mutants and was then normalised with the PCA values of the wild type. The result is a PCA centered around the wild type using the phenotypes as variables, where the distance between a given point and the wild type orange point at the center is representative of the distance in the phenotype scores among them.

We were particularly interested in identifying KO and OE mutants that depleted *Proliferation* and/or increased *Apoptosis* with regard to the wild type LNCaP model. Using MaBoSS, we were able to quantify and rank the effect of all the 32258 mutants on the probabilities of reaching Proliferation and Apoptosis (SuppFile 6).

The double mutants that mostly depleted *Proliferation* were combinations of p21_oe, MXI1_oe, HIF1_oe, AR_ko and E2F1_ko. Likewise, double mutants that mostly increased *Apoptosis* were combinations of GLI_oe, Caspase3_oe, Caspase8_oe, Caspase9_oe and PTCH1_oe. The single mutants that mostly depleted *Proliferation* were HIF_oe, MXI_oe, p21_oe, Caspase3_oe and Caspase8_oe. Likewise, those that mostly increased *Apoptosis* were GLI_oe, Caspase3_oe, Caspase8_oe, Caspase9_oe and SMO_oe

It was in our interest to identify drugs that could inhibit some of these genes, thus, we filtered these lists to find the best single KO mutations. We found that the single KO that mostly depleted *Proliferation* were AR_ko, VHL_ko, AKT_ko, E2F1_ko, PIP3_ko, EGFR_ko, PI3K_ko, CDH2_ko, TWIST1_ko, ERK_ko. Likewise, the single KO that mostly increased *Apoptosis* were AKT_ko, AR_ko, ERK_ko, cFLAR_ko, SPOP_ko, PIP3_ko, PI3K_ko, EGFR_ko, HSPs_ko and ATR_ko. Another knockout, p53_ko, was identified in our analysis, but was later discarded upon closer analysis. From topological analyses, p53 deletion should increase *Proliferation*, as p21, a cyclin inhibitor, is therefore not transcribed. Nevertheless, p53 has a dual effect on *Apoptosis* in the network: p53 activates CytC and Apaf1, which activate *Apoptosis*, but p53 also inhibits BIRC5, an activator of *Apoptosis*. The model should be closely inspected to correct this mismatch in future works. In any case, the effects of p53's mutations are not further analysed in present work, nor their results are further discussed.

We gathered the 20 top nodes from each of those lists and ended with 29 nodes that could be knocked out to deplete *Proliferation* and/or increase *Apoptosis* (AKT, AR, ATR, AXIN1, Bak, BIRC5, CDH2, cFLAR, CyclinB, CyclinD, E2F1, eEF2, eEF2K, EGFR, ERK, HSPs, MED12, mTORC1, mTORC2, MYC, MYC_MAX, PHDs, PI3K, PIP3, SPOP, TAK1, TWIST1, VHL). We used this ranking, the genes corresponding to these nodes and known drugs that target these genes to shortlist potential therapeutic target candidates tailored to LNCaP cell line (Main text, Table 1).

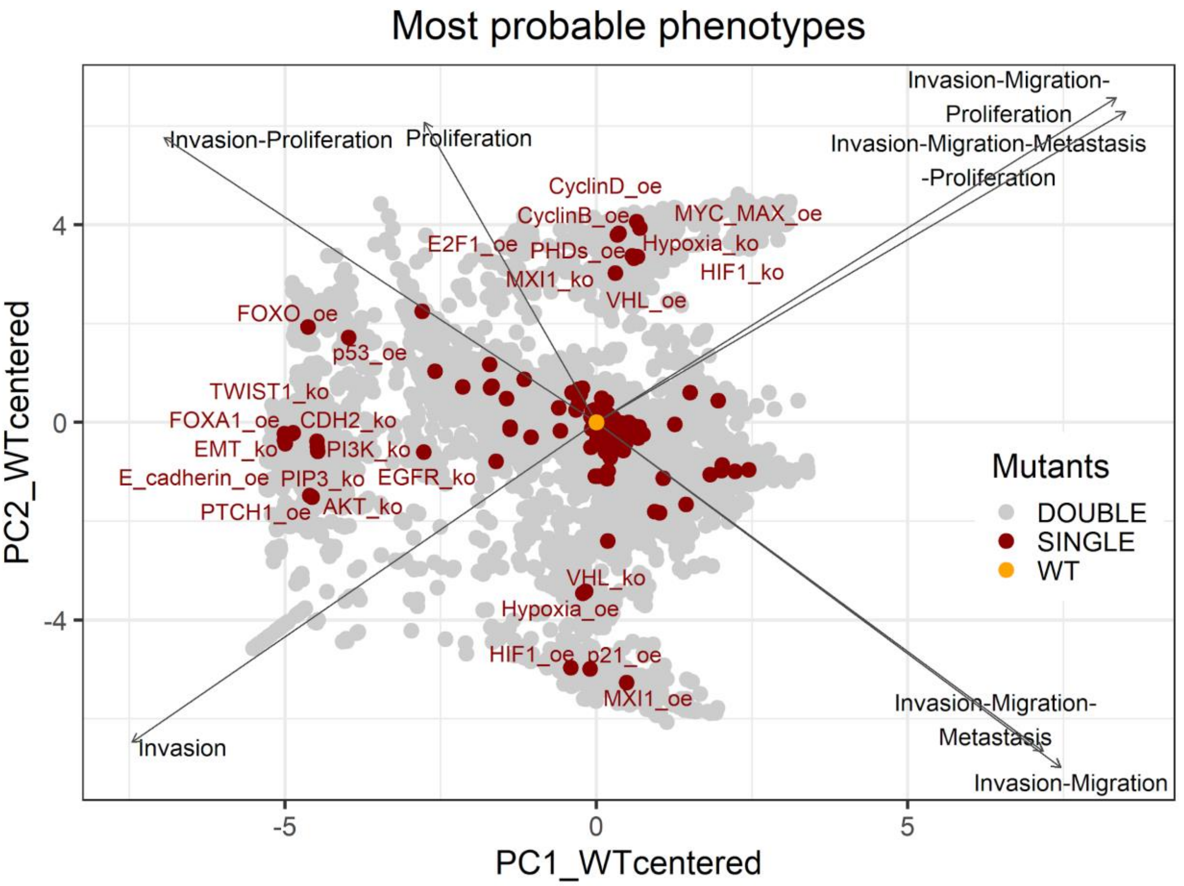


Figure S25: PCA of the 32258 single and double LNCaP model mutants with combination of phenotypes.

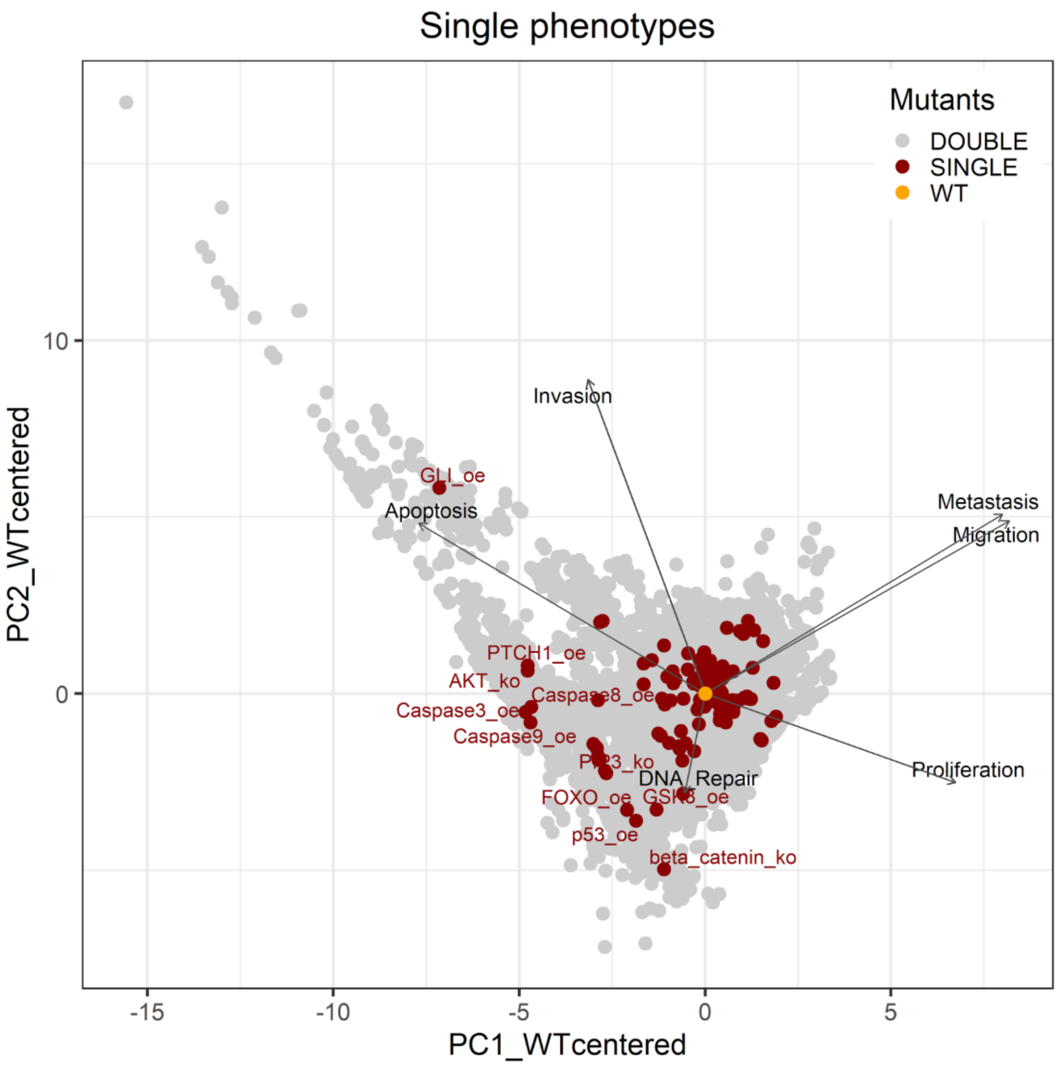


Figure S26: PCA of the 32258 single and double LNCaP model mutants with the decomposition in single phenotypes.

### Drug studies in prostate cell lines

#### Drugs associated to genes included in the model

We tested if the drugs that targeted the genes included in the model allowed us to identify cell line specificities. We analysed drug sensitivity data from GDSC1 and GDSC2 studies (Iorio *et al*, 2016) and for each drug we calculated a normalised sensitivity of the 8 prostate cell lines considered in present study (22RV1, BPH-1, DU-145, LNCaP-Clone-FGC, NCI-H660, PC-3, PWR-1E and VCaP). We normalised drug sensitivity across cell lines in the following way: cells were ranked from most sensitive to least sensitive using ln(IC50) as drug sensitivity metrics, and the rank was divided by the number of cell lines tested with the given drug. Thus, the most sensitive cell line scored 0, while the most resistant cell line scored 1 normalised sensitivity. This rank-based metric made it possible to analyse all drug sensitivities for a given cell line, without drug specific confounding factors, such as the mean IC50 of a given drug, or others.

We observed that cell lines described as resistant (DU-145 and PC-3) have a skewed distribution towards least sensitive values (Figure S27, panels D and E), while cell lines such as LNCaP have a skewed distribution towards more sensitive values (Figure S27, panel A). Meaning that the drugs that target the genes in the personalised model are not very effective against the resistant cell lines, but that LNCaP is significantly more sensitive to these. Additionally, we found that BPH-1 is generally sensitive to all drugs, let them be model-specific or not (Figure S27, panel C). For the other cell lines, there is no significant difference between model-specific drugs or not.

In addition, we performed a target set enrichment analysis using the *fgsea* method (Korotkevich *et al*, 2016) from the *piano* R package (Väremo *et al*, 2013). Again, we targeted pathway information from the GDSC1 and GDSC2 studies (Iorio *et al*, 2016) as target sets, and performed the enrichment analysis on the aforementioned normalised drug sensitivity profile of the LNCaP cell line. This target enrichment analysis showed that LNCaP cell lines are especially sensitive to PI3K/AKT/MTOR, hormone related (AR targeting) and Chromatin targeting (bromodomain inhibitors, regulating Myc) drugs (Table S1, adjusted p-values from target enrichment: 0.001, 0.001 and 0.037, respectively), which corresponds to the model predictions (Main text, Table 1).

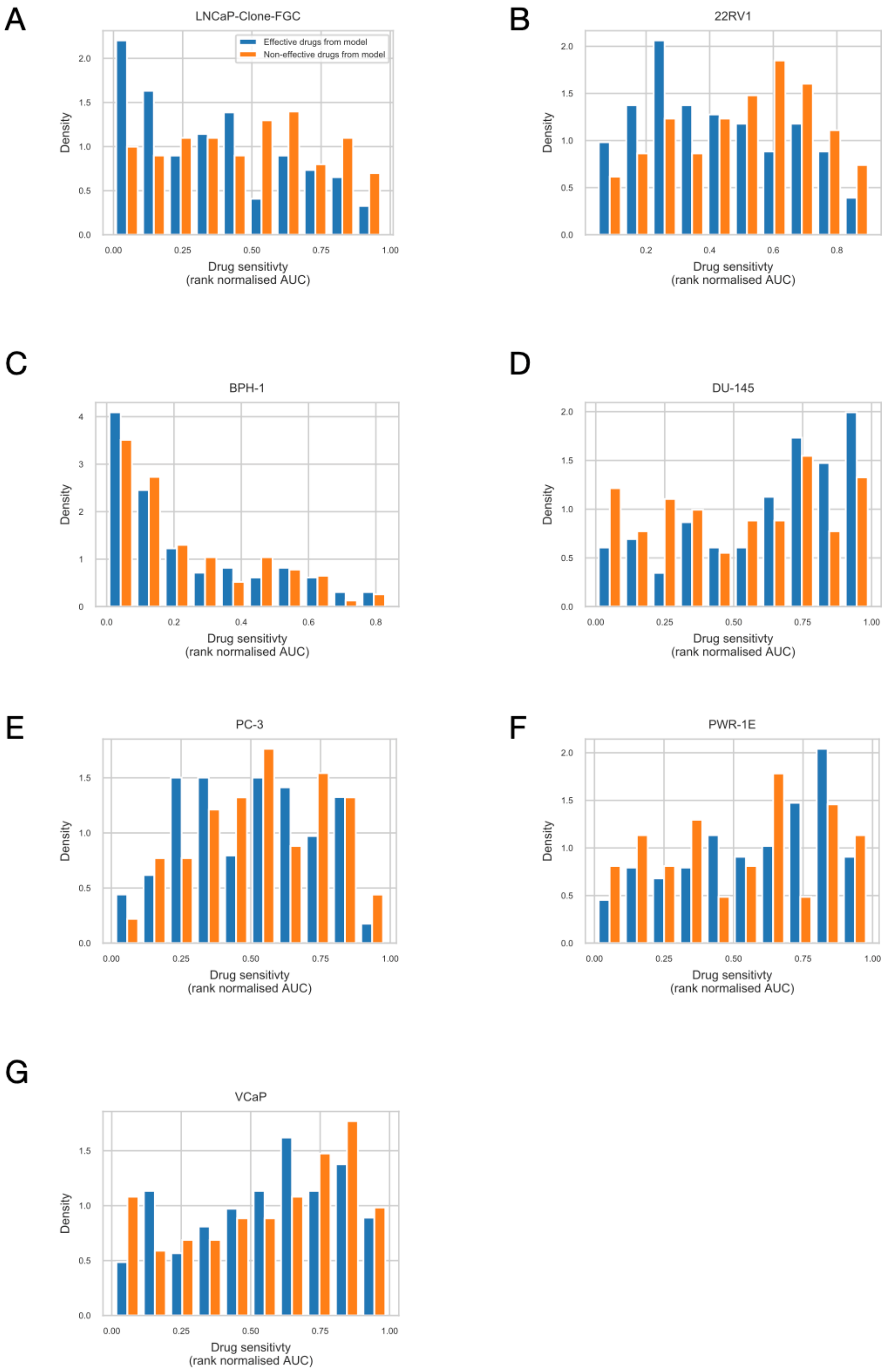


Figure S27: Drug sensitivity of the 7 prostate cell lines. Rank normalised drug sensitivity (0: most sensitive; 1: most resistant, based on GDSC AUC drug sensitivity metric) for each GDSC drug across prostate cancer cell lines. Drugs are grouped to be predicted effective drugs based on the LNCaP Boolean model (orange) and predicted ineffective drugs (blue). Mann-Whitney U p-values for differences between the rank normalised drug sensitivity between predicted effective and ineffective drugs: A) LNCaP p = 0.00041 (more sensitive to LNCaP-model-predicted drugs). B) 22RV1 p = 0.0033 (more sensitive to LNCaP-model-predicted drugs). C) BPH-1 p = 0.31. D) DU-145 p = 0.0026 (more resistant to LNCaP-model-predicted drugs). E) PC-3 p = 0.15. F) PWR-1E p = 0.075. G) VCaP p=0.38.

Table S1: Target enrichment for LNCaP-specific drug sensitivities. Drugs were sorted based on rank normalised drug sensitivity 0: most sensitive, 1 most resistant, based on GDSC AUC drug sensitivity metric for LNCaP). Target pathway enrichment analysis was performed based on the pathway membership of drug targets. Direction represents whether pathway targeting drugs were enriched in sensitive or resistant drugs.

| Drug target pathway | p-value | adj. p-value | direction |
|---|---|---|---|
| **PI3K/MTOR signaling** | 0.00011563 | **0.0011106** | **sensitive** |
| **Hormone-related** | 0.00014808 | **0.0011106** | **sensitive** |
| **Chromatin other** | 0.0065661 | **0.03283** | **sensitive** |
| **Chromatin histone methylation** | 0.01216 | **0.045601** | **sensitive** |
| p53 pathway | 0.079554 | 0.23866 | sensitive |
| DNA replication | 0.10466 | 0.26164 | sensitive |
| WNT signaling | 0.13583 | 0.29107 | sensitive |
| Unclassified | 0.20391 | 0.38233 | sensitive |
| Genome integrity | 0.54186 | 0.90311 | sensitive |
| Cytoskeleton | 0.63153 | 0.93981 | sensitive |
| Other, kinases | 0.81647 | 0.93981 | sensitive |
| RTK signaling | 0.85985 | 0.93981 | sensitive |
| Other | 0.87572 | 0.93981 | sensitive |

| | | | |
|---|---|---|---|
| Protein stability and degradation | 0.88166 | 0.93981 | sensitive |
| EGFR signaling | 0.93981 | 0.93981 | sensitive |
| Apoptosis regulation | 0.96036 | 0.96036 | resistant |
| Chromatin histone acetylation | 0.73164 | 0.83616 | resistant |
| JNK and p38 signaling | 0.63484 | 0.83616 | resistant |
| IGF1R signaling | 0.23538 | 0.37662 | resistant |
| Cell cycle | 0.19382 | 0.37662 | resistant |
| Metabolism | 0.053352 | 0.14227 | resistant |
| Mitosis | 0.027536 | 0.11014 | resistant |
| **ERK MAPK signaling** | 0.00050075 | **0.004006** | resistant |

#### Drugs associated to proposed targets of LNCaP

We wanted to test if the LNCaP cell line is more sensitive than the rest of the prostate cell lines to the LNCaP-specific drugs identified in Table 1 from the main text. We compared GDSC's Z-score of these drugs in LNCaP with their Z-scores in all GDSC cell lines (Figure 5). We observed that LNCaP is more sensitive to drugs targeting AKT or TERT than the rest of the studied prostate cell lines. In Figure S28 we can observe that trend in comparison to the other prostate cell lines and to the rest of the GDSC cell lines. In addition, we see that AKT sensibility in LNCaP is one of the highest in GDSC records.

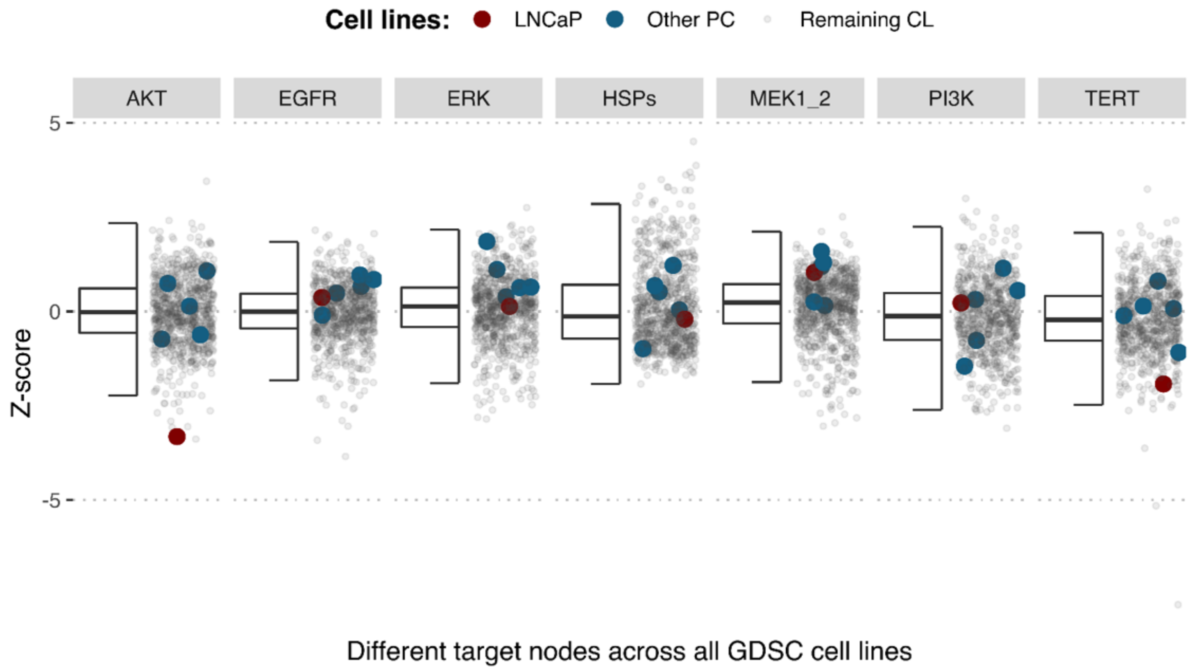


Figure S28: Model-targeting drugs' sensitivities across prostate cell lines. GDSC Z-score was obtained for all the drugs targeting genes included in the model for all the prostate cell lines in GDSC. LNCaP is highlighted in red, the other 7 prostate cell lines in blue and the rest of the GDSC cell line are coloured in grey.

### Gradual inhibition of genes in LNCaP model

All these inhibitions were performed using our PROFILE_v2 framework (https://github.com/ArnauMontagud/PROFILE_v2) that allow to study the effect of single and double mutations (knock-out and overexpression) in the phenotypes' probabilities using MaBoSS as well as to study the Bliss Independence synergy score of these combinations.

### Single inhibitions

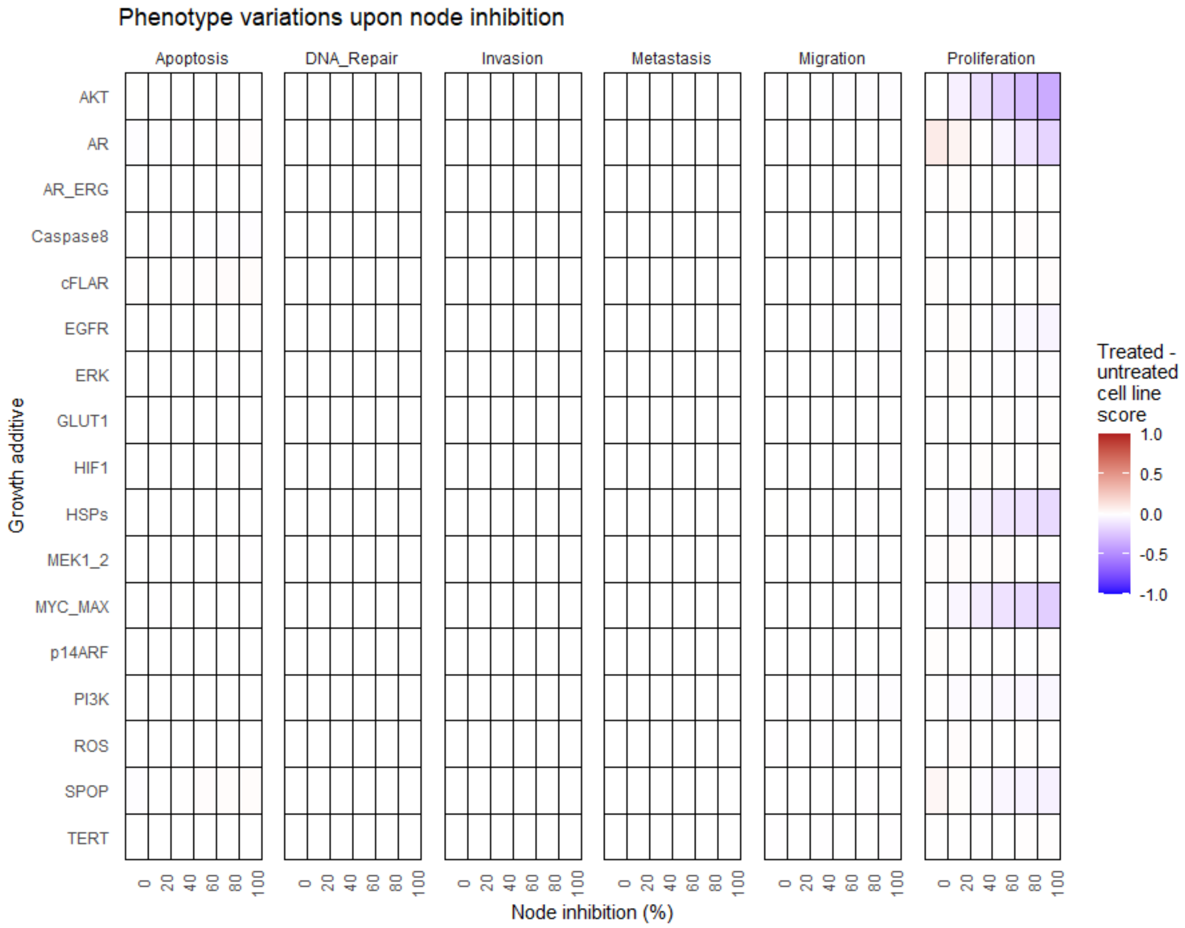


Figure S29: Phenotype score variations upon nodes inhibition under *EGF* growth condition. Values of the scores are depicted with a colour gradient.

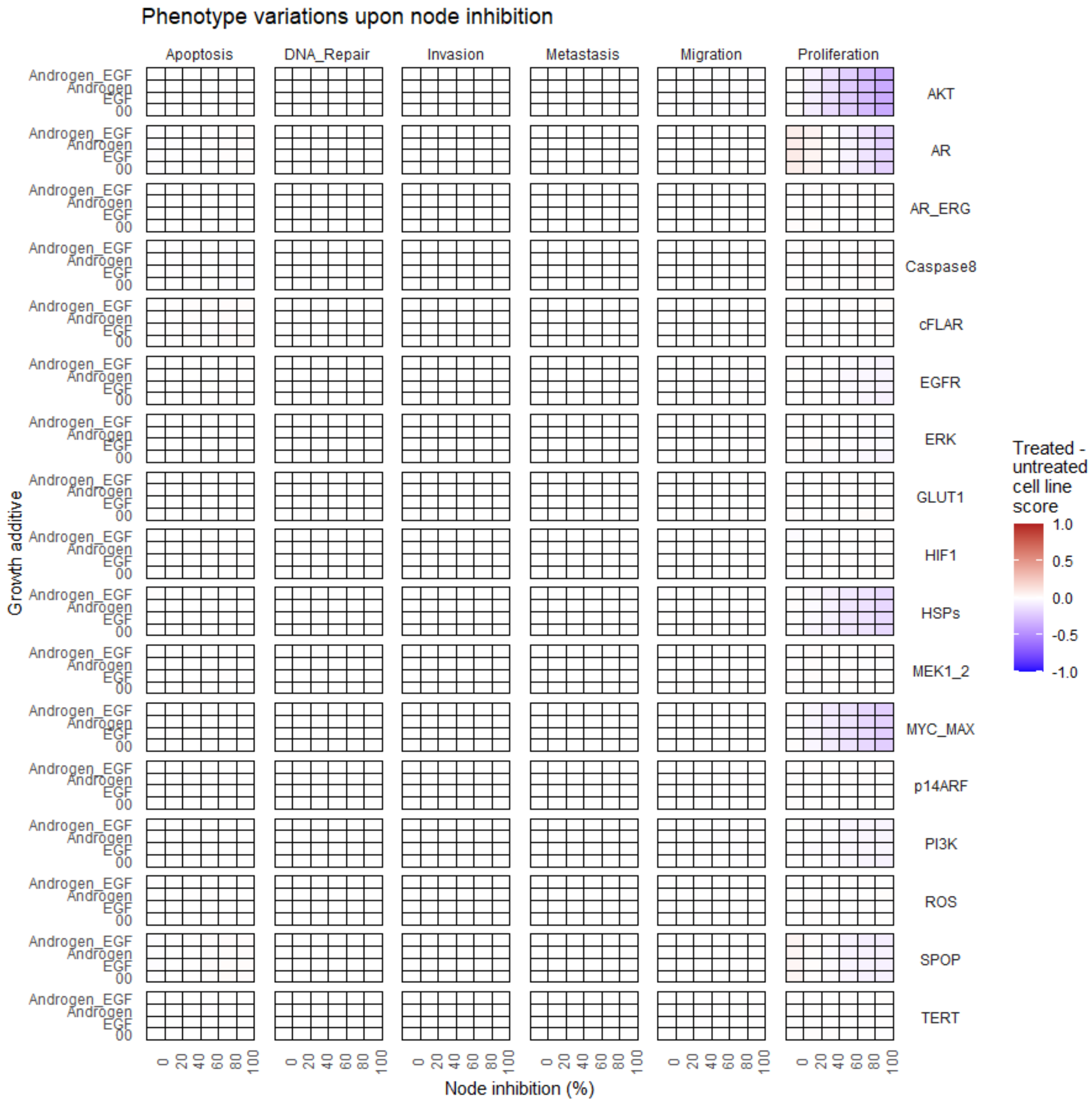


Figure S30: Phenotype score variations upon nodes inhibition under *AR, EGF, 00* and *AR_EGF* growth conditions. Values of the scores are depicted with a colour gradient.

### Double inhibitions

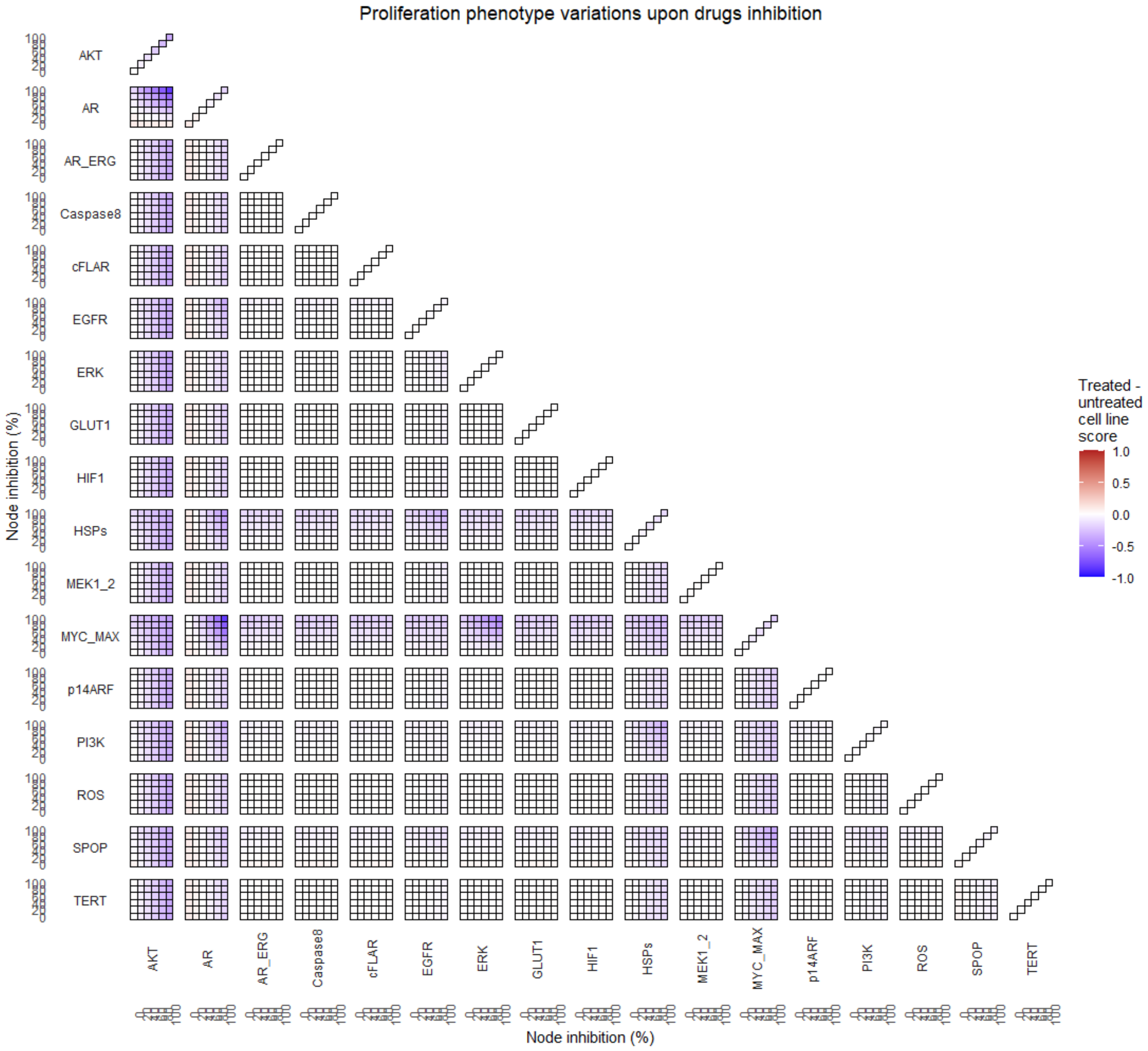


Figure S31: Proliferation phenotype score variations upon combined nodes inhibition under *EGF* growth condition. Figure 4A is a closer look at ERK and MYC_MAX combination and Figure 4B at HSPs and PI3K combination.

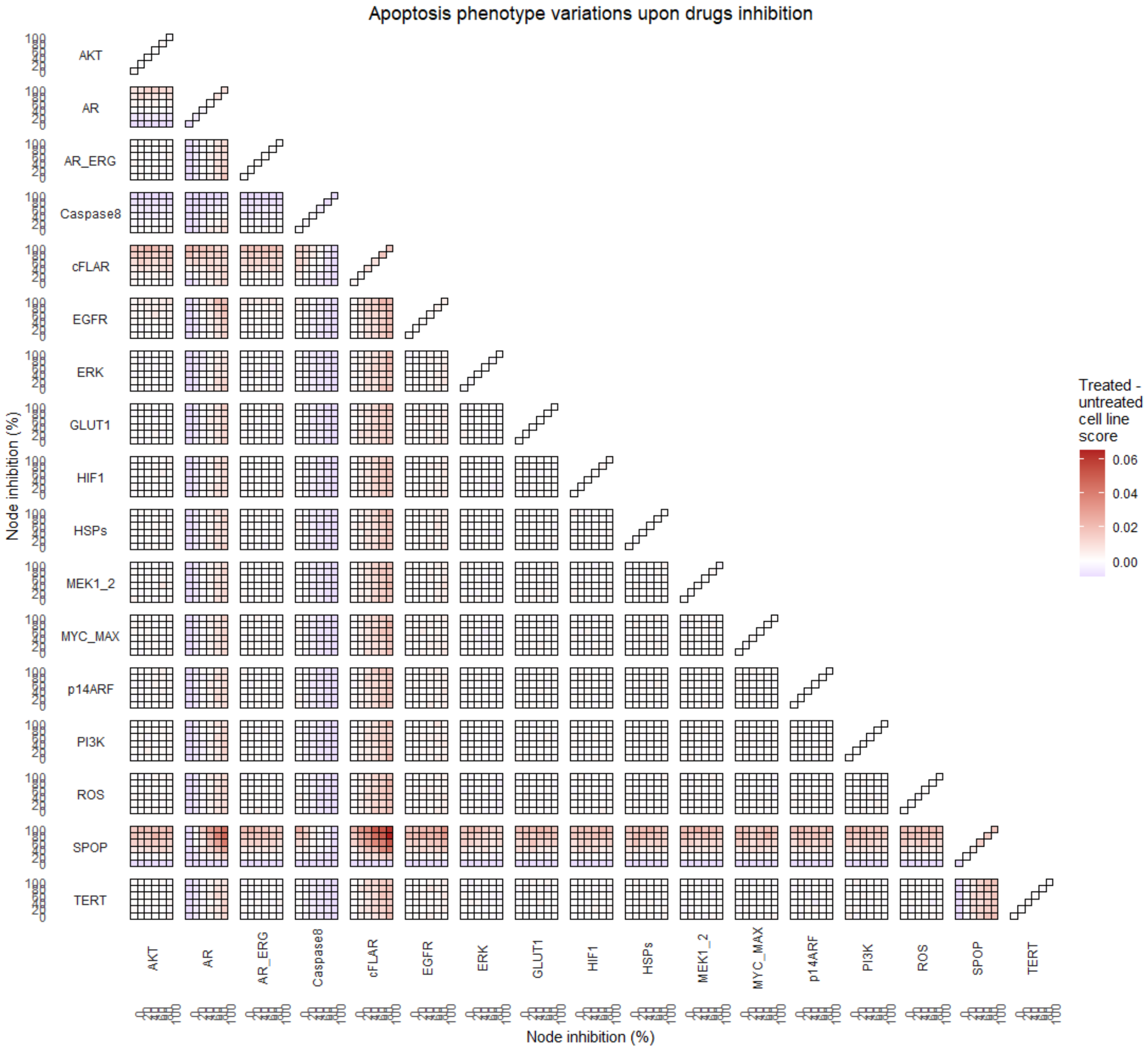


Figure S32: Apoptosis phenotype score variations upon combined nodes inhibition under *EGF* growth condition.

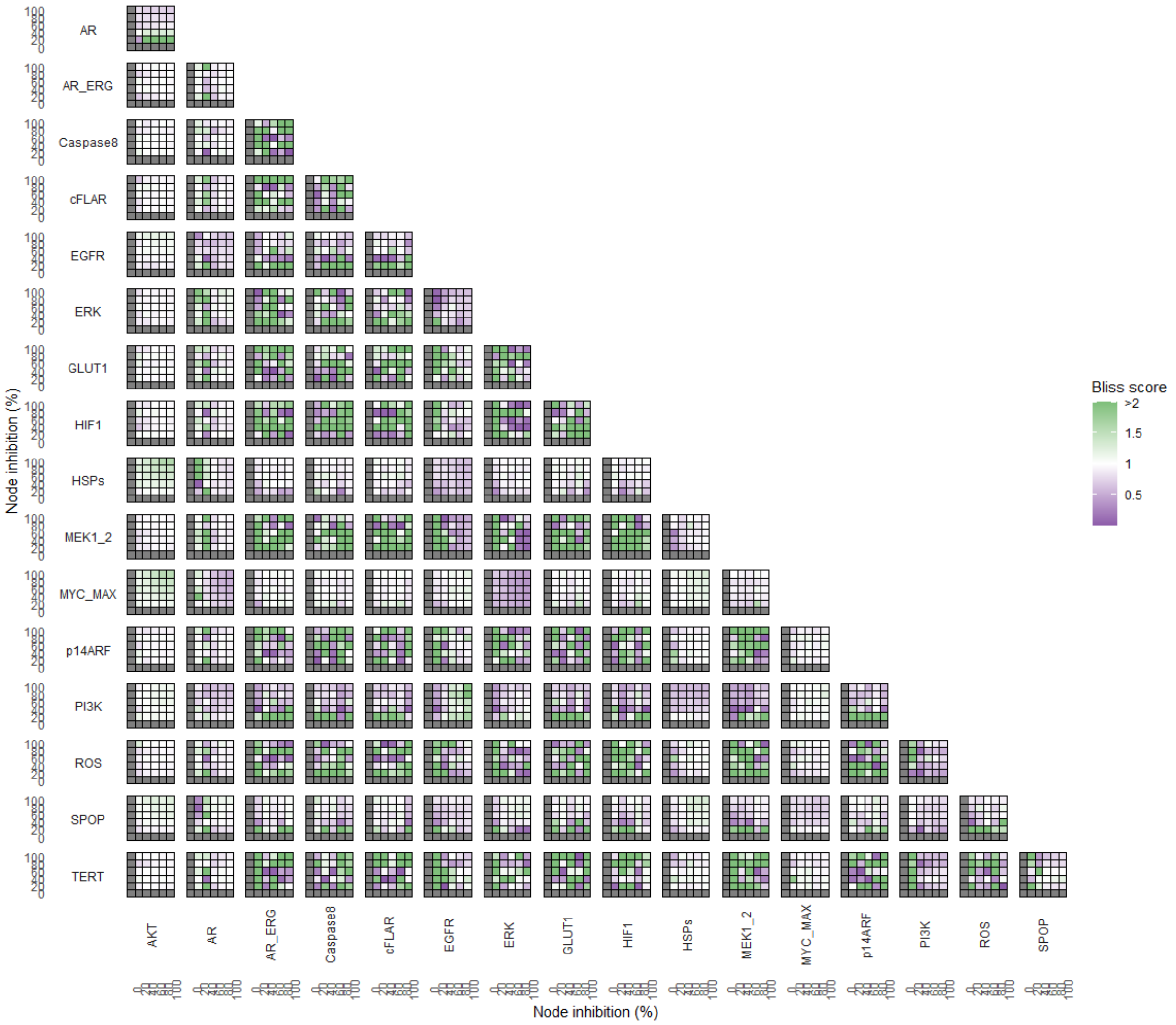


Figure S33: Bliss Independence synergies scores variations in Proliferation phenotype upon combined nodes inhibition under *EGF* growth conditions. Bliss Independence synergy score < 1 is characteristic of drug synergy. Figure 4C is a closer look at ERK and MYC_MAX combination and Figure 4D at HSPs and PI3K combination, grey colour means one of the drugs is absent and thus no synergy score is available..

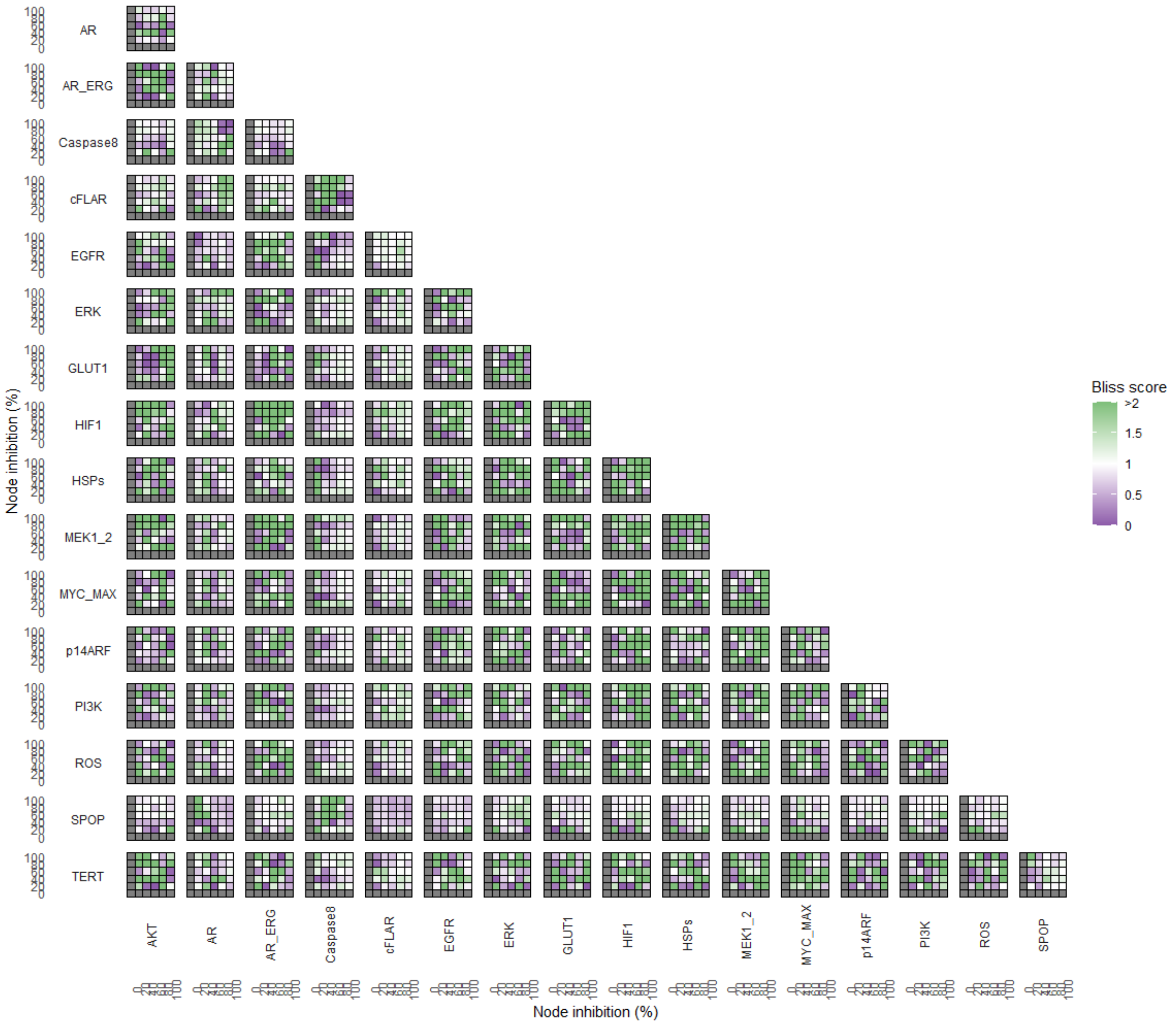


Figure S34: Bliss Independence synergies scores variations in Apoptosis phenotypes upon combined nodes inhibition under *EGF* growth conditions. Bliss Independence synergy score < 1 is characteristic of drug synergy, grey colour means one of the drugs is absent and thus no synergy score is available.
